# Joint Generation of Protein Sequence and Structure with RoseTTAFold Sequence Space Diffusion

**DOI:** 10.1101/2023.05.08.539766

**Authors:** Sidney Lyayuga Lisanza, Jake Merle Gershon, Sam Tipps, Lucas Arnoldt, Samuel Hendel, Jeremiah Nelson Sims, Xinting Li, David Baker

**Author notes:** May 8, 2023. Equal contribution (Sidney Lyayuga Lisanza, Jake Merle Gershon, Sam Tipps, Lucas Arnoldt, Samuel Hendel).

## Abstract

Protein denoising diffusion probabilistic models (DDPMs) show great promise in the *de novo* generation of protein backbones but are limited in their inability to guide generation of proteins with sequence specific attributes and functional properties. To overcome this limitation, we develop ProteinGenerator, a sequence space diffusion model based on RoseTTAfold that simultaneously generates protein sequences and structures. Beginning from random amino acid sequences, our model generates sequence and structure pairs by iterative denoising, guided by any desired sequence and structural protein attributes. To explore the versatility of this approach, we designed proteins enriched for specific amino acids, with internal sequence repeats, with masked bioactive peptides, with state dependent structures, and with key sequence features of specific protein families. ProteinGenerator readily generates sequence-structure pairs satisfying the input conditioning (sequence and/or structural) criteria, and experimental validation showed that the designs were monomeric by size exclusion chromatography (SEC), had the desired secondary structure content by circular dichroism (CD), and were thermostable up to 95°C. By enabling the simultaneous optimization of both sequence and structure, ProteinGenerator allows for the design of functional proteins with specific sequence and structural attributes, and paves the way for protein function optimization by active learning on sequence-activity datasets.

## 1 Main

Protein function arises from a complex interplay of sequence and structural features, hence designing new protein functions requires reasoning over both sequence and structure space. Many protein design methods sample structures and sequences in separate steps, typically by generating protein backbones first and using inverse folding methods to generate sequences. Traditional methods like Rosetta flexible backbone protein design^1^ alternate between structure and sequence design, while recent deep learning based approaches typically generate backbones first and then use sequence design methods such as ProteinMPNN to identify sequences that fold into a given backbone^2–5^. Among the latter class of approaches, denoising diffusion probabilistic models^6^ (DDPMs), which have shown considerable promise in continuous data domains allow for the generation of protein backbones subject to a wide range of structural constraints^7–9^. DDPMs approximate the probability density function over a data distribution by learning to denoise samples corrupted with Gaussian noise, enabling the generation of high-quality samples from a Gaussian prior; they have been explored less in categorical domains such as text and protein sequences^2, 3, 5, 10^. Simultaneous generation of sequence and structure could have advantages over methods that alternate between optimization in the two domains independently by enabling coordinated guidance with both sequence and structural features. Hallucination approaches that apply activation-maximization to structure prediction networks^11–13^ can generate sequence-structure pairs without additional training, but these solutions can be adversarial, require a large number of steps to converge, and robust experimental success requires subsequent sequence design on the hallucinated backbones^3^.

We reasoned that diffusion approaches could be powerful for simultaneous generation of sequence and structure while avoiding the adversarial solutions of activation maximization, and set out to develop a diffusion model which jointly generates sequence-structure pairs and can be guided by constraints in both domains. We hypothesized that RoseTTAfold’s ability to simultaneously generate protein sequences and structures, as illustrated by RoseTTAFold Joint Inpainting^11^, could be adapted for diffusive generation of coherent sequence-structure pairs by finetuning to recover noised native protein sequences while imposing a loss on structure prediction accuracy, and that such a DDPM could be readily guided by constraints in both domains.

### 1.1 DDPM Implementation

We chose to implement diffusion in sequence space by representing amino acid sequences as scaled one hot tensors where true values are set to 1 and all other values set to -1, allowing progressive corruption with Gaussian noise N(*µ* = 0*, σ* = 1) ^14, 15^. This approach is advantageous over other categorical diffusion methods, where diffusion occurs within a learned embedding space of text^16, 17^, because it simplifies the use of raw sequence based classifiers for guidance. To finetune RoseTTAFold we input the protein sequences progressively noised according to a square root schedule^16^, the corresponding time step, and optional structural information. We task the model to generate ground truth sequence-structure pairs by applying a categorical cross entropy loss to the predicted sequence (relative to the ground truth sequence) and FAPE structure loss on the predicted structure. Self-conditioning^14^, which allows the model to condition on its previous prediction, was employed to improve training and inference performance. Protein generation begins with an Lx20 dimensional sequence of Gaussian noise, and at each timestep (**x_t_**) the model predicts **x_0_** from **x_t_**, after which **x_0_**is noised to **x_t-1_** (Figure 1A, top panel). Conditioning information (guidance) can be combined with **x_0_** to guide the model towards a constrained sequence space using activity data, sequence specific potentials, secondary structure features, and more (Figure 1A, bottom)^18^.

**Figure 1:**
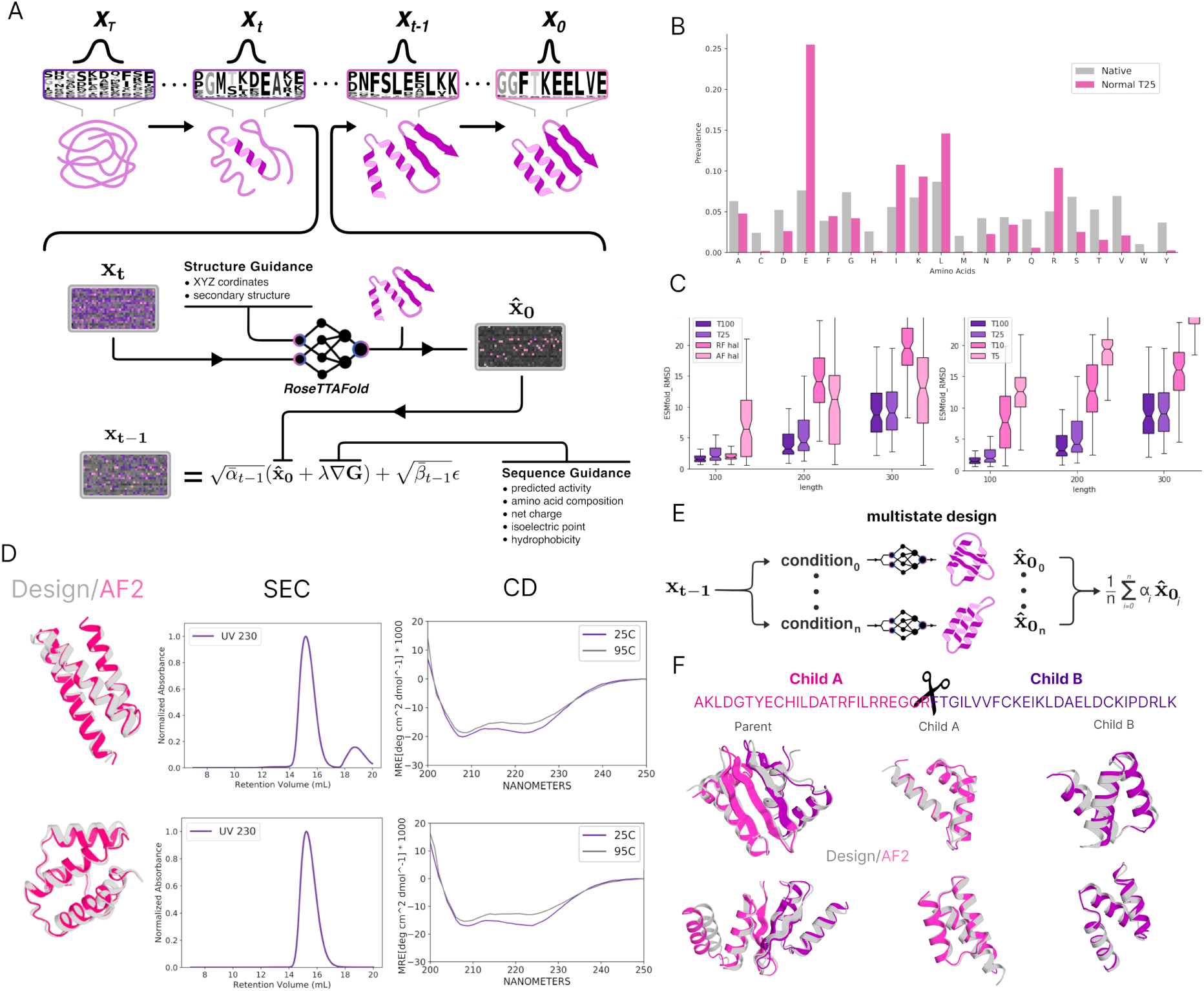
Overview of ProteinGenerator. (A) Inference schematic indicating how a noised sequence **x_t_** is passed through the model with structural conditioning and updated to **x_t-1_**for the next pass. At each step in the diffusion process **x_0_**is predicted from **x_t_**and guidance can be added to the predicted **x_0_**prior to scaling with noise. This process is repeated for T steps as the sequence-structure pair converges on a high confidence solution. (B) Sampling from a Gaussian distribution yields sequences that approach that of native sequences. (C) Unconditional designs have higher ESMfold pLDDT and lower ESMfold RMSD to design compared other joint models: RosettaFold (RF) and Alphafold2 (AF) hallucination. (D) Experimental validation of unconditional designs: Design (grey) and AlphaFold2 (pink) models of unconditionally generated proteins. Size exclusion chromatography and circular dichroism experiments show these designs are soluble, monodispersed, and thermostable to 95°C. (E) Multistate guidance allows for the design of a single sequence with a variety of structural conditioning states to converge on a single sequence predicted to adopt multiple states. (F) Use of multistate guidance to generate sequences which upon fragmentation switch from alpha/beta to all alpha secondary structure. Parent design (left) switches secondary structure when split into two child proteins (right). Designed structures of parent and children are shown in grey and AlphaFold2 predictions are overlaid in pink/purple.

### 1.2 Unconditional generation

Starting with a sequence of Gaussian noise, the model generates sequence-structure pairs with amino acid compositions similar to those of native proteins (Figure 1B, left). The generated sequences and structures are internally consistent: AlphaFold2 and ESMFold predictions of the structures adopted by the generated sequences are very close to the generated structures (Figure 1C, S1B) and confident (Figure S1A). Sampling from different noise distributions resulted in different amino acid frequencies and secondary structure compositions in the generated outputs^19^ (Figure S1A, S2, S3, S4). Samples of unconditionally generated designs with 100aa, 200aa or 300aa length can be found in Figures S5, S6 and S7. For the longer lengths the success rate of ProteinGenerator in generating sequences that fold to the designed structures is lower than that of the RoseTTAfold based structure diffusion method RFdiffusion^8^ followed by ProteinMPNN^3^; this may reflect intrinsic differences between diffusion in sequence and structure space, or arise from differences in model training.

For experimental characterization, we unconditionally generated 70-80 residue proteins, filtered for high AF2 confidence (pLDDT > 90) and AF2 RMSD to design < 2Å (Table S4). A second subset with high ProteinGenerator confidence (model pLDDT > 90) and AF2 RMSD to design < 2Å, but low AF2 confidence (AF2 pLDDT < 80) was tested as well (Table S4). Synthetic genes encoding the designs were transformed into E. coli, and the proteins were expressed and purified using nickel-NTA chromatography. Of the 42 proteins tested, 34 were soluble and monomeric by size exclusion chromatography (SEC) and circular dichroism (CD) experiments showed they had the anticipated secondary structure and were stable up to 95°C (Figure 1D).

### 1.3 Conditioning on Single or Multi-state Structural Information

ProteinGenerator can be conditioned on either explicit 3D coordinates or on secondary structure as described by per-residue DSSP features. *In silico* tests show that when conditioned on 3D structural motif information the model generates proteins accurately recapitulating these motifs (Figure S8). During training coordinates for structural motifs are provided 40% of the time either as continuous spans of 4-9 residues or as 5-10 sparse residues, distant in sequence space. For lower resolution secondary structure guidance, DSSP^20^ features were specified on a per-residue level 25% of the time and masked between 0% and 90% (randomly sampled). This allows guidance of a part or all of a structure towards a specific secondary structure type or fold, which the ProteinGenerator does quite well (Figure 1F, 3A).

Designing an amino acid sequence that can adopt distinct structural conformations upon an external trigger is a challenging task, as the energy landscape must contain two discrete minima with free energy differences small enough for a trigger to induce state switching^21^. We reasoned that ProteinGenerator was well equipped for this task because of its understanding of sequence-structure relationships and its ability to apply constraints in both domains.

We experimented with going beyond single-state structural specification by seeking to condition on distinct structural features of two different states. We applied multistate conditioning to design fold switching proteins by inputting the same sequence with two (or more) input sets of structural constraints at each step and averaging the output logits (Figure 1E). This allows the model to search sequence space for high-confidence solutions that satisfy all constraints. We used this approach to generate designs consisting of two fragments separated by a protease cleavage site which adopt different secondary structures following: in the intact parent state beta strand conditioning is used in the region flanking the cleavage site, while alpha helical conditioning is used for the two resulting subsequences. Logits from the parent and children sequences are averaged together at each step to arrive at a single sequence. As designed, AF2 structure predictions for the intact parent sequence have beta sheets in this region, whereas the two fragments (predicted independently) are entirely helical; the 3D structures of both the intact parent and the two children are very close to the design models (Figure 1F). A similar multistate approach can be applied to design monomers that adopt multiple oligomeric states and other conformationally switching systems.

### 1.4 Sequence guidance with amino-acid based potentials

An advantage of diffusion in sequence space is that sequence-based guiding functions can be readily implemented and applied. As a first test of this, we sought to design proteins with high frequencies of specific amino acids conferring structural or functional properties (cysteines can form disulfide bonds to make stable proteins, tryptophans possess spectroscopic properties, and histidines can confer pH sensitivity). Given a specification of the desired fraction of a given amino acid, at each denoising step positions are ranked based on the extent to which the output logits favor the amino acid, and the desired fraction are biased further in this direction (Figure 2A). We found this allowed more fine grained control in generating sequences than imposing a global bias towards a particular amino acid. We used this procedure to generate proteins with high frequencies (20%) of tryptophan, cysteine, valine, histidine, and methionine one at a time (Figure 2B, S9). We obtain compositionally biased protein sequences composed of nearly 20% of the desired amino acids that are strongly predicted to adopt the corresponding structures.

**Figure 2:**
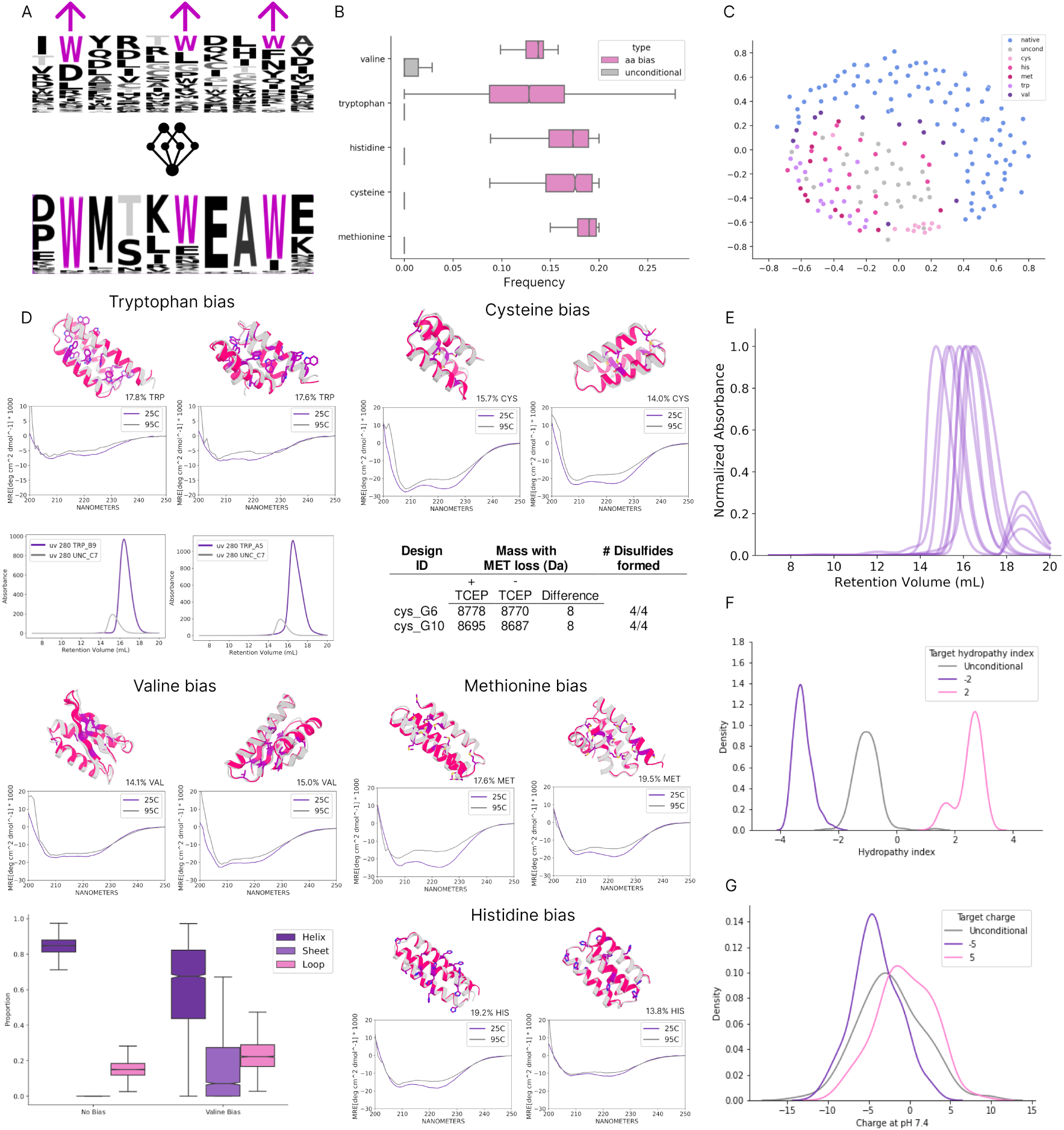
Generation of folded proteins with target sequence compositions. (A) Amino acid compositional bias: Applying an amino acid compositional bias generates proteins with desired sequence composition (tryptophan bias shown as an example). (B) Generation of compositionally biased proteins using ProteinGenerator. Bar plot displaying the average amino acid composition of designed proteins when guiding for 20% amino acid of interest compared to unconditional generation. (C) Sequence diversity representation: Multidimensional scaling of compositionally biased protein sequences occupy distinct spaces. (D) Experimentally validated amino acid compositionally biased proteins: Proteins generated with upweighted tryptophan, cysteine, valine, methionine, and histidine are predicted with high confidence by AlphaFold2 (pink), match the design model (grey), and are thermostable up to 95°C by circular dichroism. Proteins designed with high tryptophan bias show five-fold higher absorbance at 280nm compared to unconditionally generated proteins. Experimental validation of cysteine-rich proteins under reducing and non-reducing conditions confirms the presence of designed disulfide bonds by mass spectrometry. Proteins designed with high valine bias show increased beta strand propensity compared to unconditionally generated designs. (E) Size exclusion chromatography overlay: Proteins designed with sequence potentials are soluble and monodisperse by size exclusion chromatography. (F) Hydrophobic sequence potential: Biasing the sequence away or toward hydrophobic amino acids results in a shift in the distribution of hydrophobicity scores for the output sequences. (G) Net charge sequence potential: Resulting distribution of net charges when guidance is used to bias sequences toward +/- 5 net charge.

To evaluate the compositionally biased designs experimentally, we generated 70 to 80 residue proteins with different amino acids upweighted, filtered on AF2 pLDDT > 90 and AF2 RMSD to design < 2Å, and experimentally characterized the top 96 designs (Table S4). Of the characterized designs, SEC traces indicated the proteins were monomeric for 4/5 upweighted cysteine proteins, 8/19 upweighted tryptophan proteins, 19/22 upweighted valine proteins, 10/12 upweighted histidine proteins, and 10/10 upweighted methionine proteins. CD spectra were obtained for a subset of the monomeric designs, and in all cases indicated secondary structure was consistent with the designed structure (Figure 2D,E). Guiding for high cysteine content at the sequence level resulted in the formation of 3 to 5 disulfide bonds per protein without any structural conditioning as indicated by mass spectrometry in the presence and absence of the reducing agent TCEP at 50mM (Figure 2D, Table S2). Proteins designed with upweighted tryptophans exhibited high absorbance at 280 nm, and proteins with upweighted valine exhibited higher beta sheet content by CD (Figure 2D middle, right). These results indicate the model understands general sequence to structure relationships beyond the typical sequence space of native proteins (Figure 2C).

We next explored the generation of proteins with prespecified charge composition, isoelectric points and hydrophobicity which can influence solubility, activity, subcellular location^22^, pharmacokinetic clearance, and retention^23^. Biasing away from hydrophobic amino acids can lead to better expression and solubility^24^, and designing towards hydrophobic interfaces is advantageous for protein-protein interactions^25^. We implemented sequence based potentials to guide^18^ the diffusive process towards these characteristics to enable fine-tuned control over physical properties of the output sequence. This approach enabled the design of proteins with a range of user-defined hydrophobicities (Figure 2F) and isoelectric points (Figure 2G).

### 1.5 Scaffolding bioactive sequences

The design of proteins with activities conditional on an outside input is of considerable general interest, and could enable generation of therapeutics with spatial and temporal control^26^. As a first exploration of the use of ProteinGenerator for such proteins, we sought to scaffold bioactive peptide sequences within an inert protein cage. Unlike our previous LOCKR^27, 28^ sensor system, in which the bioactive sequence must be in a helical conformation and make specific interactions with the caging scaffold, the generality of ProteinGenerator requires only that the sequence of the bioactive peptide be specified–neither the structure this adopts nor the structure of the overall cage need be decided on in advance.

ProteinGenerator is able to generate structures containing peptide sequences corresponding to known lytic peptides and the designed sequences are confidently predicted to adopt the designed structures (Figure 3B). We used this approach to scaffold a bioactive peptide in the terminus of a protein that can be conditionally released upon proteolytic cleavage of a terminal loop (Figure 3C). We specified the sequence, not the structure, of the bioactive segment, and used DSSP conditioning to force the cleavage motif to be in a loop. We chose to scaffold the pore forming peptide melittin^29^ currently being explored as a cancer therapy^30^. Starting with the melittin sequence and a flanking cleavage site, we generated an additional 125 residues to scaffold the peptide into a globular protein. Melittin-scaffolded proteins generated by the model were in agreement with AlphaFold2 models (AF2 pLDDT > 85, AF2 RMSD < 2Å) (Figure 3C). We obtained synthetic genes encoding 12 proteins scaffolding melittin and found that 9/12 were monodisperse by SEC and had the correct secondary structure by CD (Figure 3C).

**Figure 3:**
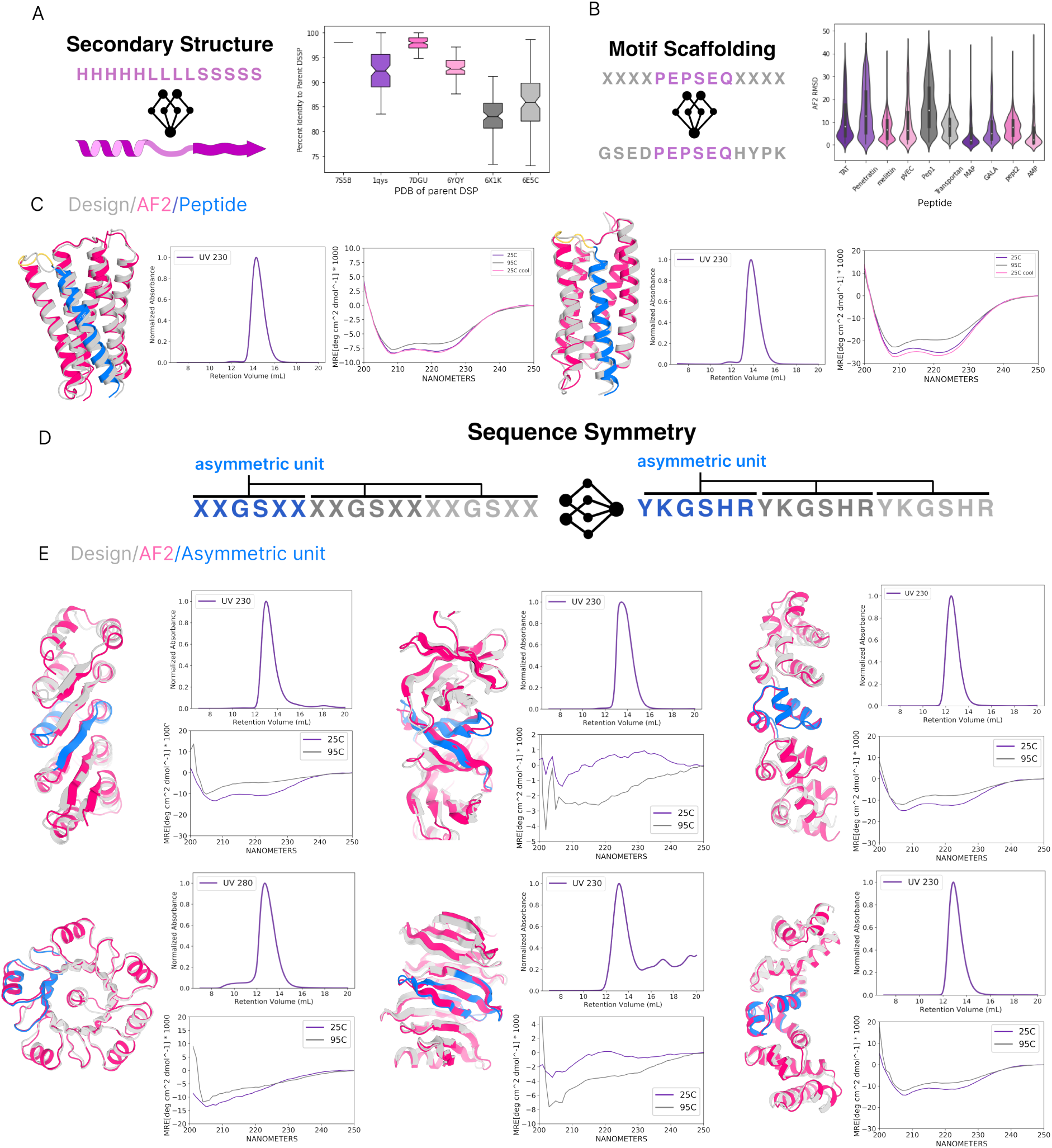
Scaffolding bioactive peptides and designing repeat proteins with ProteinGenerator. (A) Secondary structure conditioning: Protein generation with secondary structure conditioning recapitulates the DSSP of the target protein. (B) Unstructured (sequence only) motif scaffolding: Sequence motif scaffolding of unstructured bioactive peptides yields designs with low AlphaFold2 RMSD to design. (C) Melittin scaffold designs: Scaffolding melittin yields designs (grey) that are corroborated by AlphaFold2 (pink), are soluble and monodispersed by size exclusion chromatography, and thermostable to 95°C by circular dichroism. (D) Sequence symmetry: Sequence repeat symmetry is applied at inference time by symmetrizing update (xt-1) sequences to generate tandem repeat proteins. (E) Experimentally characterized symmetric repeat proteins: Designed repeat proteins (grey) with secondary structure conditioning are corroborated by AlphaFold2 (pink), are soluble and monodispersed by size exclusion chromatography, and are thermostable up to 95°C by circular dichroism.

### 1.6 Generation of sequence repeat proteins

Repeat proteins containing tandem copies of a sequence-structure unit are ubiquitous in nature and play central roles in molecular recognition and signaling^31^. Previous work in designing repeat proteins has required extensive pre specification of structural features^32^. We reasoned ProteinGenerator could be used to readily generate repeat proteins given only the sequence length of the repeat unit and number of repeats desired. At each timestep we symmetrize the noised sequence distribution accordingly (Figure 3D). Unconditional generation with this approach yielded largely beta solenoid structures which AF2 corroborated. We added helical caps to a subset of designs to promote stability and reduce aggregation^33, 34^. To encourage further exploration of the repeat protein universe we specified the secondary structure for a small percent (2%-10%) of residues, which yielded a wide range of all alpha, all beta, and mixed alpha-beta designs (Figure 3E). We generated 165-185 residue repeat proteins, filtered them (AF2 pLDDT>85 and RMSD to design < 2), and experimentally characterized 74 repeat proteins with helical caps and 86 repeat proteins without helical caps. Of these, 27 repeats with caps and 10 repeats without helical caps were soluble and monomeric by SEC, and 7/8 proteins evaluated using circular dichroism had the expected secondary structure (Figure 3E, Table S3^35^).

### 1.7 Guidance with sequence only classifiers

Designing proteins with a desired biological activity is a long standing goal of *de novo* protein design. An advantage of our approach is that diffusion can be directly guided by function classifiers that operate in sequence space. We first sought to guide the network with the DeepGOPlus Gene Ontology (GO) classifier^36^ to generate proteins with specific characteristics and functions. Although GO classification scores increased with guidance for nitrogen compound metabolic process (GO:0006807) and membrane (GO:0016020), we found the classifier had a high false positive rate often assigning high scores to native sequences outside the GO domain (Figure S10). In a separate approach, we trained a simple transformer encoder and single linear layer to discriminate unconditionally generated sequences from nanobody sequences and immunoglobulin (IG) folds aggregated from Integrated Nanobody Database for Immunoinformatics^37^ and Structural Classification of Proteins database^38, 39^. We generated 125 residue proteins, roughly the length of a nanobody, and found when classifier guidance or strand bias (1%) was used alone the classifier scores increased; when used in combination classifier scores increased (Figure 4A) along with the fraction of beta-strand containing proteins (Figure S11). 14% of designs made with the classifier alone were found to be beta sandwiches, which increased to 45% when applying a strand bias to 1% of the residues. Of the designs made with the combination of strand bias and classifier guidance 68.7% matched with tm-align > 0.5 to IG folds. AlphaFold models of sequences with high classifier scores matched the design models well (Figure 4B).

**Figure 4:**
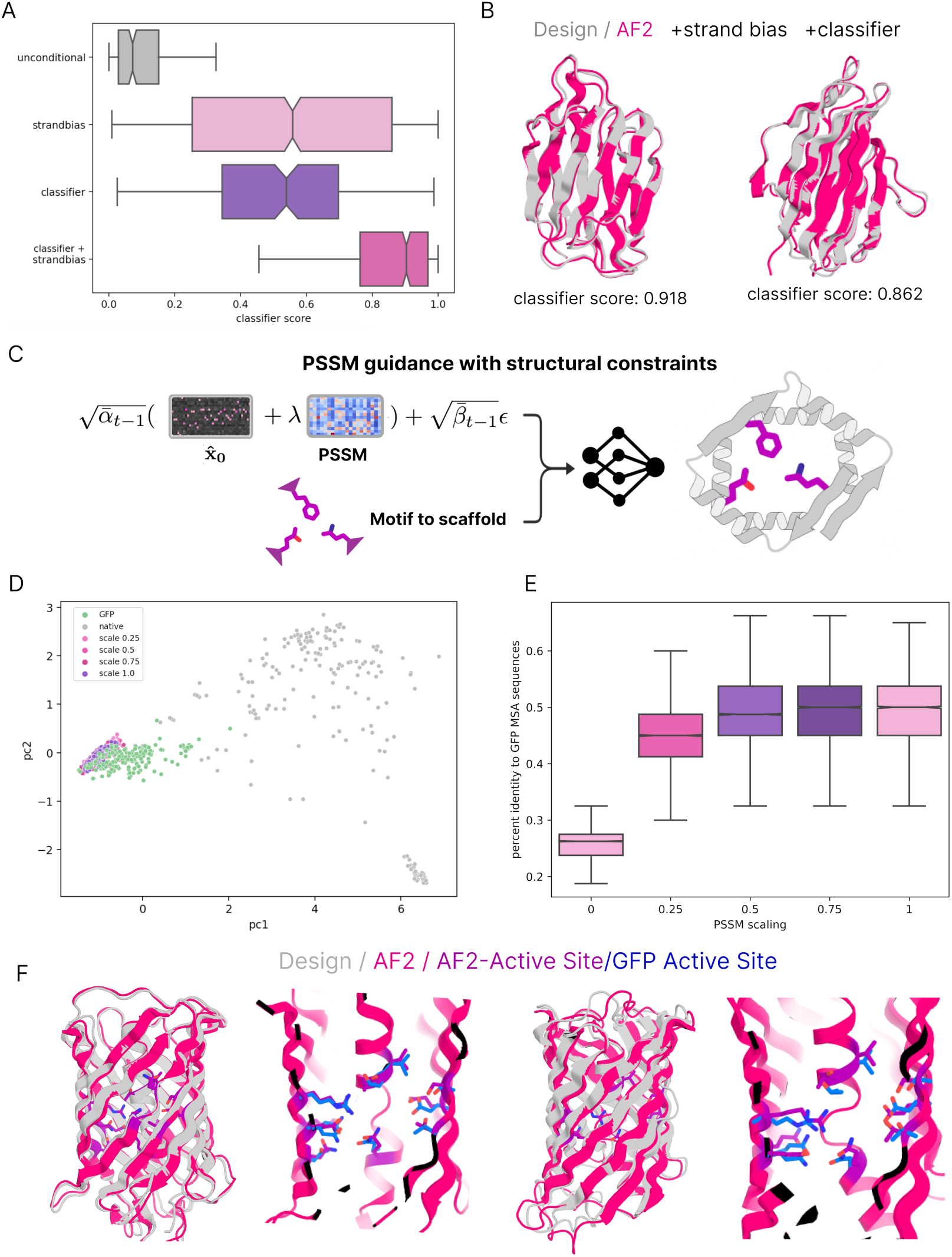
Fold and function guided protein generation. (A) immunoglobulin (IG) sequence based classifier score distributions from outputs generated unconditionally, with strand DSSP conditioning, with gradients from IG classifier, and with combination of IG classifier gradients and secondary structure conditioning. (B) Design model (grey) and AF2 model (pink) of proteins generated with classifier guidance and strand DSSP conditioning. (C) Schematic of PSSM guidance with scaling and motif scaffolding. (D) Guidance by position specific score matrix (PSSM) generates sequence-structure pairs sampled from a similar embedding space as native GFP designs. Designs and natives were embedded with ESM and first and second principle components derived from embeddings are plotted. (E) Sequence identity to native GFPs increase with increasing guidance scaling: Sequence similarity of PSSM designs at varying guide scales to native GFPs. (F) PSSM guided design examples: Design models (grey) and AF2 models (pink) of proteins generated with a PSSM guidance scale of 0 (left) or 8 (right) and 30% DSSP masking. Active site residues on AF2 predicted structures (purple) over-layed with wildtype GFP side chains in blue.

### 1.8 Guidance using protein family sequence information

Protein families often have specific residues important for function that are conserved throughout the family. Position specific scoring matrices (PSSMs) capture this information and have been previously used to generate active enzymes with corresponding sequence composition using consensus sequence design^40^, but these approaches do not consider sequence-structure coherence, while Rosetta PSSM guided flexible backbone structure based sequence design^41^ can require expensive MCMC calculations. We sought to use ProteinGenerator to design new members of protein sequence families, conditioning on both family multiple sequence alignments and key structural features associated with function. We generated a PSSM for the GFP fluorescent protein family^42^ (all sequences in uniprot having greater than 30% sequence identity to GFP)^43^. At each step in denoising, we used the PSSM to bias the sequence distribution towards that of the family, and the calculations were conditioned on the coordinates of the residues contacting the chromophore (this cannot be done using purely sequence based methods) (Figure 4C). To tune guidance the PSSM was scaled by a factor of 0.25, 0.5, 0.75, and 1.0, and we found that sequences clustered closer together became more similar to GFP family members as scaling increased (Figure 4D, E, S12). AlphaFold2 and ESM-Fold are unable to predict native GFP from a single sequence (Figure S13) but predict sequences generated by the model with high accuracy to the design (Figure 4F). Active site coordinates provided as conditioning are in close agreement between the AF2 models and the native model, and demonstrate the model’s ability to condition on both sequence and structural features (Figure 4F).

## 2 Discussion

By taking advantage of the ability of RoseTTAFold to jointly model protein sequences and structures, ProteinGenerator is able to directly sample in sequence space while ensuring that the 3D structure is coherent and satisfies any desired constraints. While RFdiffusion^8^, Chroma^9^, and other protein backbone diffusion models have demonstrated success in generating complex backbones with precise control over structural features, ProteinGenerator designs not only protein backbones but also sequences, enabling the design of proteins with any combination of desired sequence and structural attributes. We anticipate that our sequence space diffusion approach could also be employed with large language models that also generate structures, such as ESMfold^44^. A particularly attractive application is active learning-based protein design/engineering optimization: given a sequence-activity predictor, iterating between ProteinGenerator design of structurally coherent proteins predicted to have high activity, experimental testing, and updating the activity predictor could be a powerful path to achieving high activity.

## 3 Methods

### 3.1 Sequence representation

To apply the diffusion framework in sequence space, a continuous representation of the categorical sequence data is needed. To implement this we represented the sequence, **x_0_**, with dimensions Lx20 where L corresponds to the protein length with 20 possibilities for each amino acid type. This takes the form of a one-hot encoded vector that is centered at zero by multiplying the Lx20 tensor by 2 and subtracting 1. Each logit within the tensor is a real number, with higher values corresponding to a higher probability for that specific amino acid at that position. With this representation we noise **x_0_** to obtain **x_t_** with the below equation following Ho et al. formulation for a standard forward process sampling from Gaussian noise with mean at 0 and standard deviation of 1.

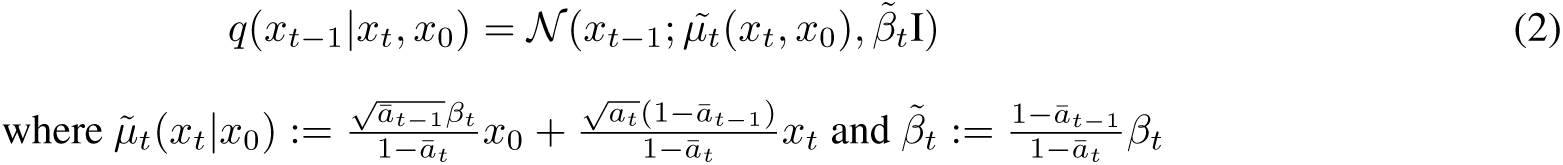

A critical part of the forward diffusion process is selecting the noising schedule. Determining the correct bin of a categorical distribution is trivial at low time steps by argmaxing the input sequence. Therefore more noise should be present at low timesteps to increase the difficulty of the task during training. The square root noise schedule^16^ satisfies this requirement and was employed in this study.

### 3.2 Training

To train the model we began by sampling t uniformly from [0,T], where t=0 is an un-noised sequence and t=T is pure Gaussian noise. We then noise **x_0_** to **x_t_** with equation (1) and tasked the model to predict the un-noised sequence **x_0_** and its corresponding structure **y**. The timestep feature was added to the sequence template passed to the model. We applied a categorical cross entropy loss to **x_0_** and structure losses to **y** (FAPE, bond angle, bond length, distogram, lddt). An additional KL loss^16^ was applied to the calculated **x_t-1_**. Self conditioning^14^ was implemented to allow the model to condition on the previous **x_0_**prediction and the back calculated **x_t-1_**during both training and inference. To self condition in practice the model was used with gradients turned off to first predict **x_0_** from **x_t+1_**, which was then passed in as a sequence template to the model. During training, RoseTTAFold was allowed 1 to 3 uniformly sampled “recycle” steps to refine structure predictions via multiple passes through the model^45^. Pseudo training and inference code is available in the supplementary information (Pseudocode S1, S2). In later training iterations secondary structure conditioning was provided to the model by concatenating a tensor representing DSSP features onto the sequence template. These features were provided 25% of the time and masked uniformly between 0% and 90% when provided.

Along with the standard diffusion task (40% of the time), the model was also challenged with structure prediction (seq2str) and fixed backbone sequence design (30% of the time each). Incorporating these additional tasks during training helped maintain the agreement of sequence-structure pairs diffused by the model. Training examples were conditioned on sequence or structure by either unmasking 1 to 4 spans of residues, each 4 to 8 amino acids in length to simulate motif scaffolding, or unmasking randomly selected residues for the model to scaffold as an active site scaffolding problem. Unmasked structure conditioning information was supplied to the input for RoseTTAFold as templates in the 1D sequence track as well as the 2D and 3D structural information tracks.

### 3.3 Inference

During inference starting from **x_t_** the model predicts **x_0_** and simultaneously decodes it to **y**. **x_0_** is then back calculated to **x_t-1_** with equation (1) and passed through the network with the previously predicted **x_0_** to apply self conditioning. Benchmarking against conditioning on **x_t_** as done in Ho et al^6^ with equation (2), shows this approach performs better (SF 1C), as seen in other categorical diffusion methods^15, 16^.

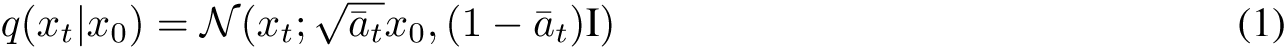

This is done for T steps, but T can be varied, and does not have to be what was used during training. The model finds solutions to some problems in as little as 10 steps (Figure 1C). Furthermore, clamping the model’s output logits from -3,3 gives better agreement with AF2 predictions (Fig S1B). **x_t-1_**is sampled from either a zero-mean Normal distribution or a Non-Bayesian Gaussian Mixture distribution with equal mixing probabilities. For the Non-Bayesian Gaussian Mixture models we defined a mixture with two Normals centered at [-1, 1] (GMM2) and a mixture with three Normals centered at [-1, 0, 1] (GMM3).

### 3.4 DSSP guidance

For constructing the DSSP features we calculated each training example’s DSSP based on the structure with helix, strand, loop, and masked labels^20^. During training, the calculated per-residue secondary structure features were appended to RoseTTAFold’s t1d features, and were one-hot encoded for 25% or 50% of the time and masked for 30% or 80% of the time. During inference, DSSP features are appended to the t1D features as necessary and masked when not.

### 3.5 Classifier guidance

For classifier guidance we utilized the DeepGOPlus model^36^ and trained a vanilla transformer model (2 Multihead Attention heads with each 2 layers, Embedding dimension: 64, Hidden Layer dimension: 64) on the INDI database for nanobodies^37^. Classifier guidance was implemented as described by Dhariwal and Nichol, 2021^18^.

### 3.6 Multistate guidance

Parent and child protein pairings were generated in 25 steps using DSSP features “XH-HHHHHHHHHHHHLLLHHHHHHHHHHHHHLLLEEEEEELLLEEEEEXXXXXXXEEEEEL-LLEEEEEELLLHHHHHHHHHHHHHLLLHHHHHHHHHHHHHX” for parent, “XHHHHHHHHHHHHH-LLLHHHHHHHHHHHHHLLLHHHHHHHHHHHHHHHHHHHX” for child A, and “XHHHHHHHHHHHHHH-LLLHHHHHHHHHHHHHLLLHHHHHHHHHHHHHX” for child B. A mixing coefficient of 0.25 was used to combine parent and child sequences together at each step. Multistate design pseudocode implementation is available in the supplements (Pseudocode S3).

### 3.7 Single Sequence Prediction with AlphaFold2

All designs used in ESMFold benchmarks were modeled by using the curl command to predict single sequence structures. All designs used in AlphaFold2 (AF2) benchmarks and ordered for experimental characterization were predicted in single-sequence structure prediction mode with model 4. Pairwise backbone RMSDs between the design model and AF2 model were calculated for each design.

### 3.8 Sequence Identity calculations

Blast alignment was used to examine sequence alignment and similarity using query coverage >90% and target coverage >50%. Alignment to natives was done against Uniref-90.

### 3.9 Unconditional Protein Generation

Unconditionally generated proteins were assessed against a set of thousand native proteins with a length deviating up to five residues randomly sampled from the RCSB database. For experimental verification proteins ranging from 70-80 amino acids in length with no conditioning information were generated in 25 steps. Designs were filtered by AF2 pLDDT > 90 and AF2 RMSD to design < 2 Å for ordering final constructs. Additionally, proteins with high model confidence but moderate AF2 confidence were ordered by filtering on design pLDDT > 90, AF2 pLDDT < 80, and AF2 RMSD to design < 5 Å.

### 3.10 Compositionally Biased Protein Generation

Proteins ranging from 70-80 amino acids in length with an amino acid compositional potential were generated in 25 steps. Designs were filtered by AF2 pLDDT > 90, AF2 RMSD to design < 2 Å, and SAP score^46^ < 30. The top 10 to 22 designs were ordered for each upweighted amino acid type (tryptophan, cysteine, valine, histidine, and methionine). Pseudocode for the implementation of the amino acid compositional potential is provided in the supplements (Pseudocode S4).

### 3.11 Charge Biased Protein Generation

Proteins of 50 amino acids in length with charge potentials applied were generated in 25 steps with charge conditioning information. The ground truth charge for each protein was calculated at pH 7.4 by using the Henderson-Hasselbach equation. Pseudocode for the implementation of the charge potential is provided in the supplements (Pseudocode S5A, S5B).

### 3.12 Hydrophobic Biased Protein Generation

Proteins of 50 amino acids in length with hydrophobic potentials applied were generated in 25 steps with hydrophobicity conditioning information. The ground truth hydropathy index for each design was calculated by summing the hydropathy index for each residue and dividing by the sequence length^47^. Pseudocode for the implementation of the hydrophobic potential is provided in the supplements (Pseudocode S6A, S6B).

### 3.13 PSSM guidance

We formulate guidance with a PSSM by simply adding a precalculated PSSM to the output **x_0_** prediction. This can be further scaled with a scaling factor, λ, to either promote stronger agreement with the PSSM, or lower, to promote more diversity. PSSMs were calculated from MSAs generated by mmseqs2^48^ with a 30-90% sequence identity cutoff to the query GFP sequence. Designs were filtered on AF2 pLDDT > 80 and AF2 RMSD to design < 2 Å.

### 3.14 Repeat protein generation

Repeat proteins ranging from 125-150 amino acids in length were generated in 50 steps with and without DSSP conditioning information. Designed proteins contained 5 repeat units using one of the one of the following DSSP strings, where X represents mask, E represents strand, and H represents helix: "XXXXEEEEEXXXXXXXXXXXXXXXHHHHHXXXX", "XXXXEEEEEXXXXXHHHHHXXXXXEEEEEXXXX", "XXXXHHHHHXXXXXEEEEEXXXXXHHHHHXXXX", "XXXXHHHHHXXLXXHHHHHXXLXXHHHHHXXXX", "XXXXEEEEEXXHXXEEEEEXXHXXEEEEEXXXX", "XXXXXXXXXXXXXXXXXXXXXXXXXXXXXXXXX", "XXXXEXXXXXXXXXEXXXXXXXXXEXXXXXXXX".

Designs were filtered on AF2 pLDDT > 80 and AF2 RMSD to design < 2 Å.

### 3.15 Scaffolding Bioactive peptides

Proteins of 155 amino acids in length were generated in 25 steps with spans of helical DSSP conditioning to encourage the model to generate helical bundles and cleavage loops. Additional DSSP features were provided to scaffold the furin protease cleavage site “GRRKR”. The sequence for melittin was provided without DSSP conditioning as N-terminal 26 amino acids “GIGAVLKVLTTGLPALISWIKRKRQQ”. Designs were filtered by AF2 pLDDT > 85, AF2 RMSD to design < 2 Å, and SAP scores < 40.

### 3.16 Plasmid construction

Protein designs were cloned into plasmids as in Watson et al^8^. Briefly, designs were ordered as synthetic genes (eBlocks, Integrated DNA Technologies) with BsaI overhangs compatible with a ccdB-encoding expression vector LM0627^13^. Genes cloned into LM0627 result in the following sequence: MSG-design-GSGSHHWGST**HHHHHH**, (SNAC cleavage tag and **6XHis affinity tag** are indicated). We used the NEBridge® Golden Gate Assembly Kit (New England Biolabs) with a total reaction volume of 5 µL and a ratio of 1:2 by mass of LM0627 plasmid DNA to design. We then incubated the reaction mixture at 37 C for 30 minutes, halted the reaction by incubating the reaction mixture at 60 C for 5 minutes, and transformed 1 µL of the reaction mixture into 6 µL of BL21 competent cells (New England Biolabs). After heat shock and recovery in SOC media, transformed BL21 cells were grown overnight in 1.0 mL of LB from which glycerol stock were created and small-scale expression cultures were inoculated.

### 3.17 1 mL-scale protein purification

Initially, proteins were expressed with small-scale expression screens as previously reported^13^ with small adaptations. Briefly, designs were inoculated with 100 uL of overnight growths and 900 uL of auto-induction media (sterile-filtered TBII media supplemented with 50 µg/mL kanamycin, 2 mM MgSO4, 1X 5052) in deep-well 96-well plates. 16 hours post-inoculation, cells were harvested and lysed in lysis buffer (50 mM Tris-HCl (pH 8), 0.5 M NaCl, 30 mM imidazole supplemented with 1X BugBuster, 1 mM PMSF, 0.1 mg/mL lysozyme, 0.1 mg/mL DNase). Clarified lysates were added to a 50 µL bed of Ni-NTA agarose resin in a 96-well fritted plate equilibrated with wash buffer (50 mM Tris-HCl (pH 8), 0.5 M NaCl, 30 mM Imidazole). After sample application and flow through, the resin was washed three times with wash buffer, and samples were eluted in 200 µL of elution buffer (50 mM Tris-HCl (pH 8), 0.3 M NaCl, 0.5 M imidazole, 5 mM EDTA (pH 8)). All eluates were sterile filtered with a 96-well 0.22µm filter plate (Agilent 203940-100) prior to size exclusion chromatography (SEC). Protein designs were then screened via SEC using an AKTA FPLC outfitted with an autosampler capable of running samples from a 96-well source plate. Samples were run on a SuperdexS75 Increase 5/150 GL column (Cytiva 29148722; 3,000 to 70,000 Da separation range) in a running buffer (20 mM Tris pH 8, 150 mM NaCl). To improve peak resolution, the SEC column was connected directly in line from the autosampler to the UV detector. 0.25 mL fractions were collected from each run. Absorption spectra were collected by the AKTA U9-M at 230 nm and 280 nm.

### 3.18 50 mL-scale protein purification

Proteins selected for further downstream characterization were expressed in 50 mL of auto-induction media. 16 hours post-inoculation, cells were harvested and lysed in lysis buffer (50 mM Tris-HCl (pH 8), 0.5 M NaCl, 30 mM imidazole, 1 mM PMSF, 0.1 mg/mL lysozyme, 0.1 mg/mL DNase) through sonication. Clarified lysates were added to a 2 mL bed of Ni-NTA agarose resin in a 20 mL column (Bio-Rad 7321010) equilibrated with wash buffer (50 mM Tris-HCl (pH 8), 0.5 M NaCl, 30 mM Imidazole). After sample application and flow through, the resin was washed 3 times with 10 mL wash buffer, and samples were eluted in 2 mL elution buffer (50mM Tris-HCl (pH 8), 0.5M NaCl, 200mM Imidazole). All eluates were sterile filtered with a 3 mL 0.22uM filter plate prior to SEC. Protein designs were then screened via SEC using an AKTA FPLC outfitted with an autosampler capable of running samples from a 96-well source plate. Samples were run on a SuperdexS75 Increase 10/300 GL column (Cytiva 29148721; 3,000 to 70,000 Da separation range) in a running buffer (20 mM Tris pH 8, 150 mM NaCl). 1 mL fractions were collected from each run. Absorption spectra were collected by the AKTA U9-M at 230 nm and 280 nm.

### 3.19 Cysteine bias protein expression

Proteins guided towards high cysteine content were transformed into and expressed in Rosetta-gami B(DE3) Competent Cells (Novagen 71137). The 1 mL and 50 mL scale protein purification protocols were otherwise followed.

### 3.20 Circular Dichroism

Circular dichroism (CD) spectra were collected on a Jasco J-1500 CD Spectrometer with 1 nm bandwidth, 50 nm permanent scan rate, and data integration time of 4 seconds per read. Sample cuvettes stored in 2% Hellmanex (Hellma 9-307-011-4-507) were washed with deionized water, 2% Hellmanex, deionized water, then 20% ethanol, after which 300 uL SEC-purified protein was added for CD spectra measurements. Thermal melts were performed at 25°C and 95°C.

### 3.21 Mass Spectrometry

To identify the molecular mass of each protein, intact mass spectra were obtained via reverse-phase LC/MS on an Agilent G6230B TOF on an AdvanceBio RP-Desalting column, and subsequently deconvoluted by way of Bioconfirm using a total entropy algorithm. Disulfide formation was determined by injecting protein at 1.5 mg/mL in the presence and absence of 50 mM TCEP-HCl (Millipore Sigma 646547-10X1ML) and detecting the mass shift.

## Acknowledgements

We would like to acknowledge Lisa Li, Saana Mansoor, Dimitry Zorine, Ian Humphreys, Harley Pyles, Brian Trippe, DéJenaé Ray, Abbas Idris, Xiaochuang Han, Meerit Said, Florence Dou, Linna Ann, Kejia Wu, Derrick Hicks, Hao Nguyen, Elias Kinfu, Adam Chazin-Gray, Quoc Tran, Marlo Zorman, Namrata Anand, and Naveen Jasti for helpful discussions and support. Chris Norn for PSSM scripts. Sergey Ovchinnikov for DSSP scripts. David Chmielewski for help with experimental procedures. Nate Ennist for help with CD. Doug Tischer for developing “contig” class for processing user inputs when running inference. Ivan Anishchenko for scripts to run TM align, sequence similarity, and multidimensional scaling plots. Jue Wang and Justas Dauparas for benchmarking scripts. Minkyung Baek and Frank DiMaio for training scripts and RoseTTAFold code base. Joe Watson, David Juergens, and Nate Bennett for helpful scripts and conversations. Ian Haydon, Lance Stewart, Luki Goldschimdt, Adam Sadowski, Kandise Van Wormer, and Lauren Carter for general operations.

This work was supported by the Defense Threat Reduction Agency Grant HDTRA1-19-1-0003 (X.L.), by funding from the DARPA program Harnessing Enzymatic Activity for Lifesaving Remedies (HEALR) under award HR0011-21-2-0012 (X.L.), the Juvenile Diabetes Research Foundation International (JDRF) grant # 2-SRA-2018-605-Q-R (X.L.), AMGEN (S.L.), the Helmsley Charitable Trust Type 1 Diabetes (T1D) Program Grant # 2019PG-T1D026 (X.L.), the Bill and Melinda Gates Foundation Grant #OPP1156262 (X.L.), the Audacious Project at the Institute for Protein Design (J.S.), the Howard Hughes Medical Institute (J.S.).

## Code Availability

The code for this projects is available here (with the exception of training scripts): https://github.com/RosettaCommons/protein_generator. For greater accessibility, thank you to Simon Dürr and HuggingFace who supplied a GPU grant to run the model interactively in your browser: https://huggingface.co/spaces/smerle/PROTEIN_GENERATOR.

## Supplements

**Figure S1:**
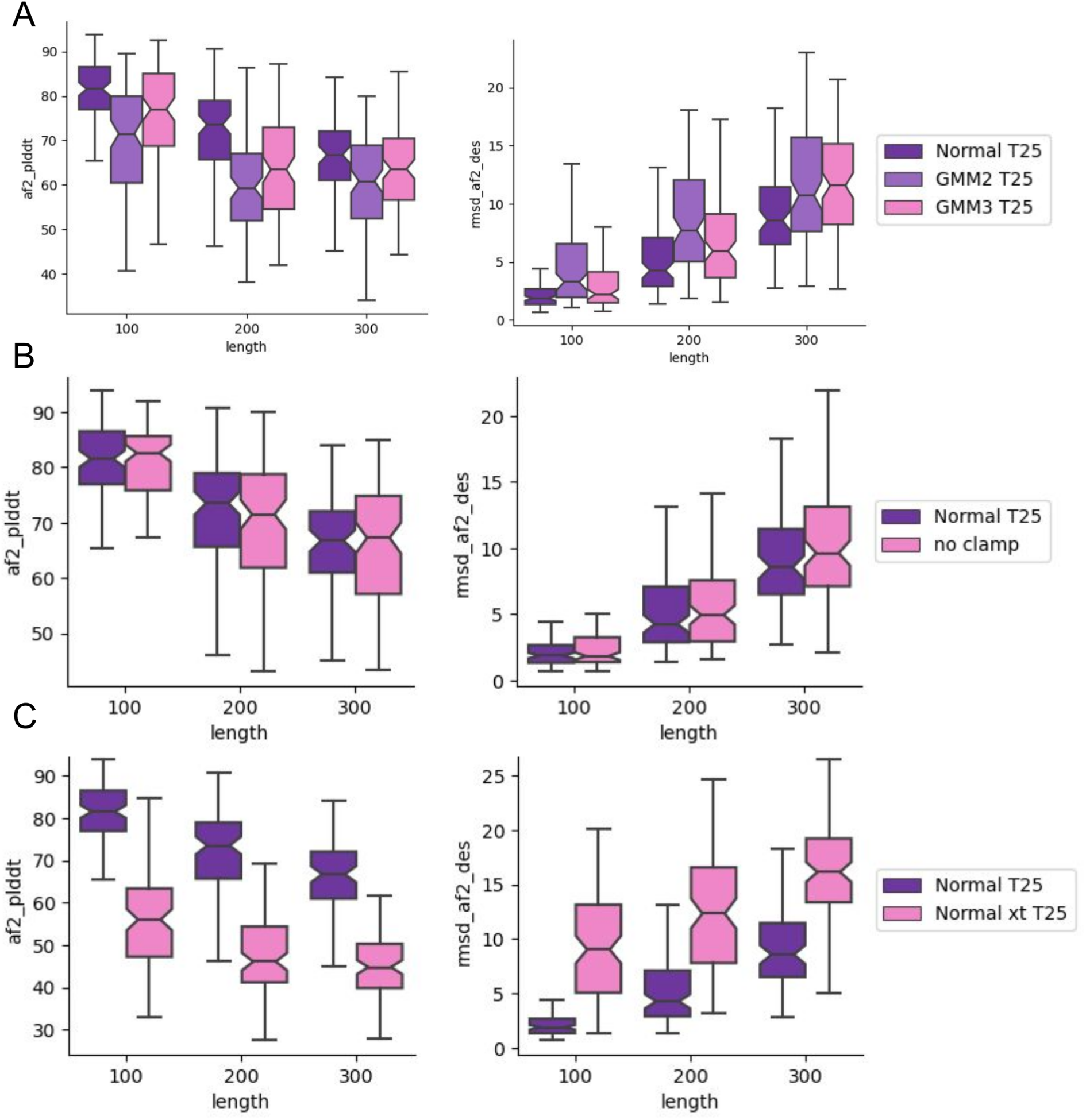
Inference benchmarks. (A) Boxplot of AF2 pLDDT of sequences from model clustered by length on left, right RMSD of AF2 model to design. (B) Boxplot AF2 pLDDT with clamp (-3,3) applied post sampling **x_t-1_** and no clamp on left, AF2 RMSD to design. (C) Comparison of with and without conditioning on **x_t_** when sampling **x_t-1_**, AF2 pLDDT right, AF2 RMSD to design left.

**Figure S2:**
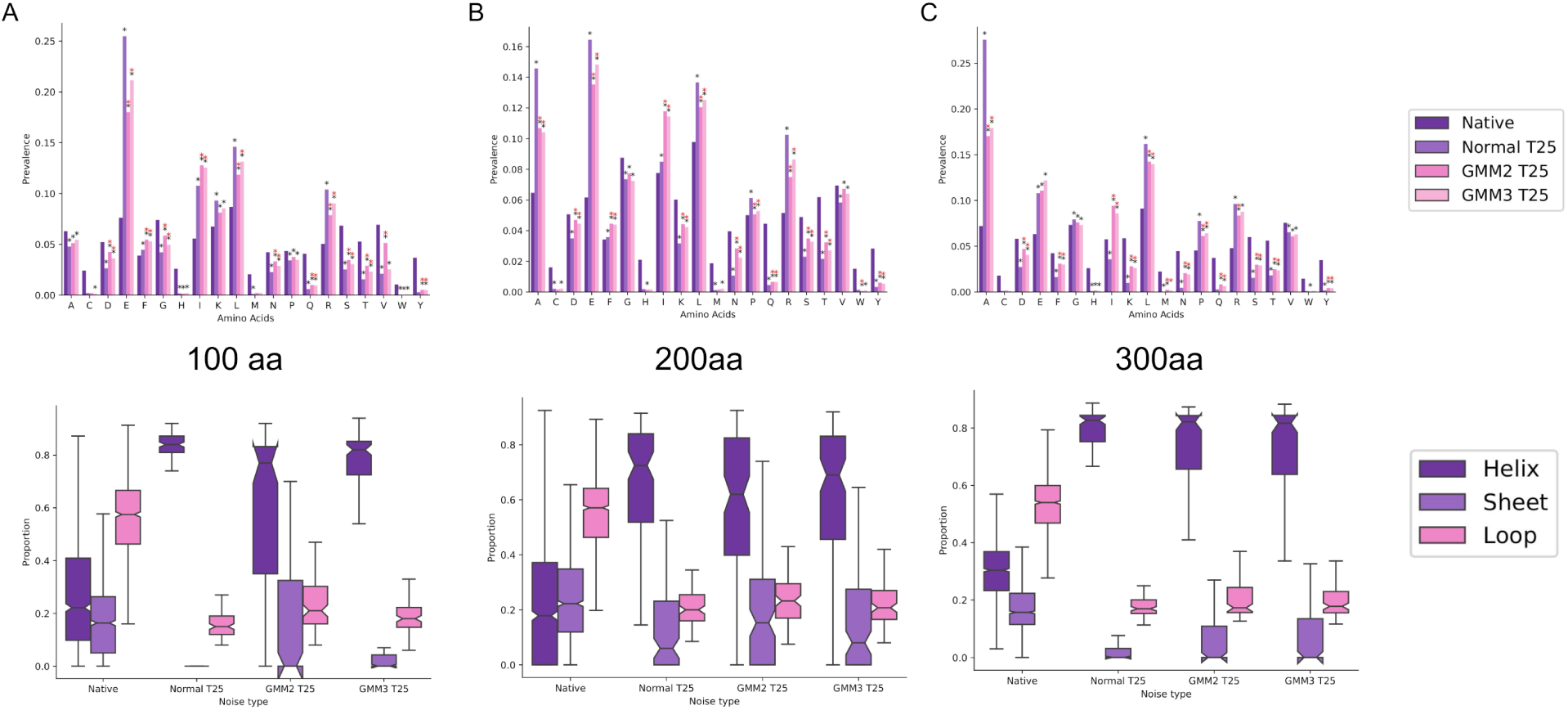
Amino acid distributions and secondary structure propensities for (A) 100AA, (B) 200AA, and (C) 300AA length proteins when sampling from normally distributed noise, GMM2, or GMM3. Significant amino prevalence changes between native and unconditional designs as well as between unconditional designs sampled from normal noise and sampled from GMM2 or GMM3 noise are displayed via black asterisk and a red asterisk.

**Figure S3:**
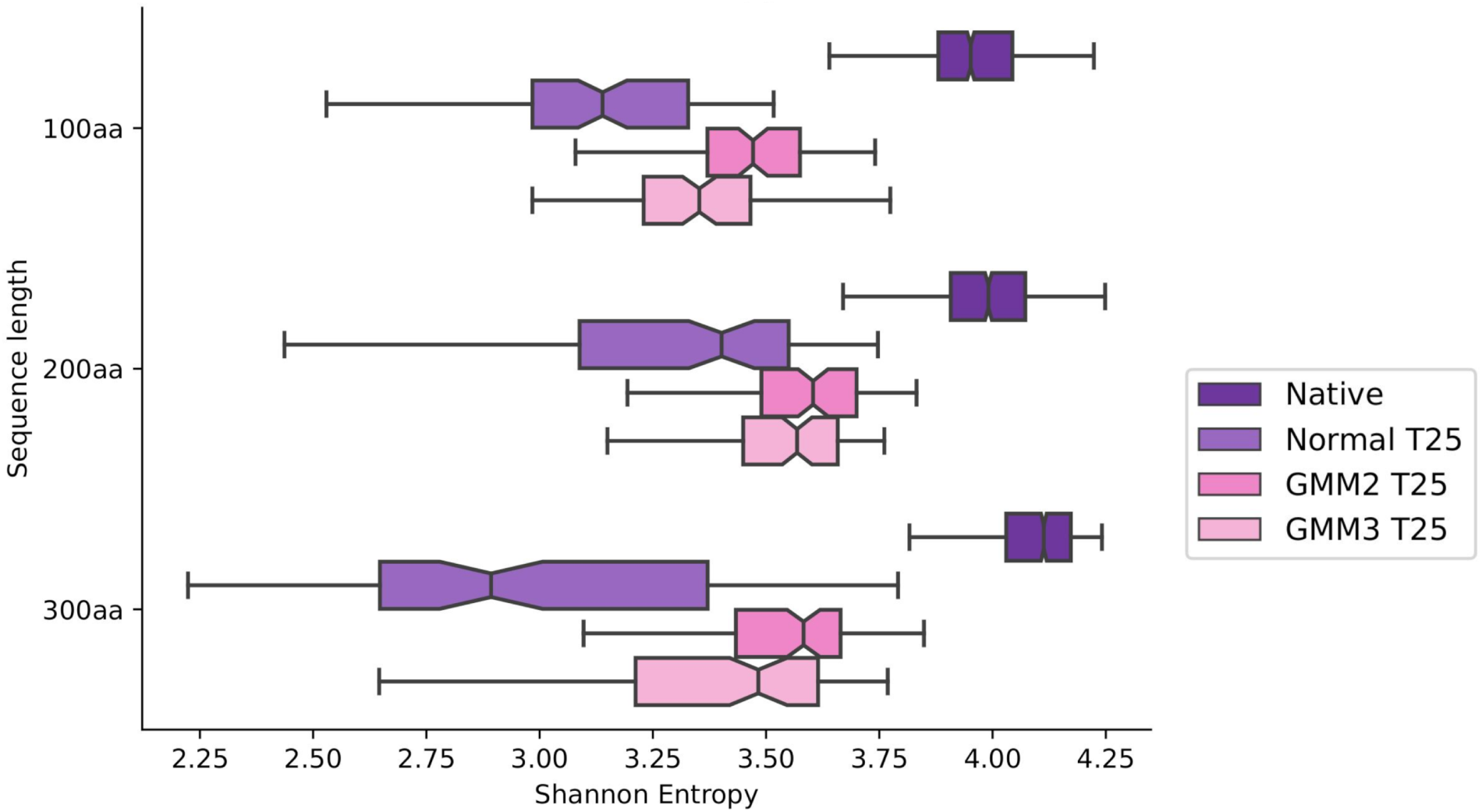
Sequence entropy of native proteins, normally sampled sequences, GMM2, and GMM3 for 100AA, 200AA, and 300AA proteins.

**Figure S4:**
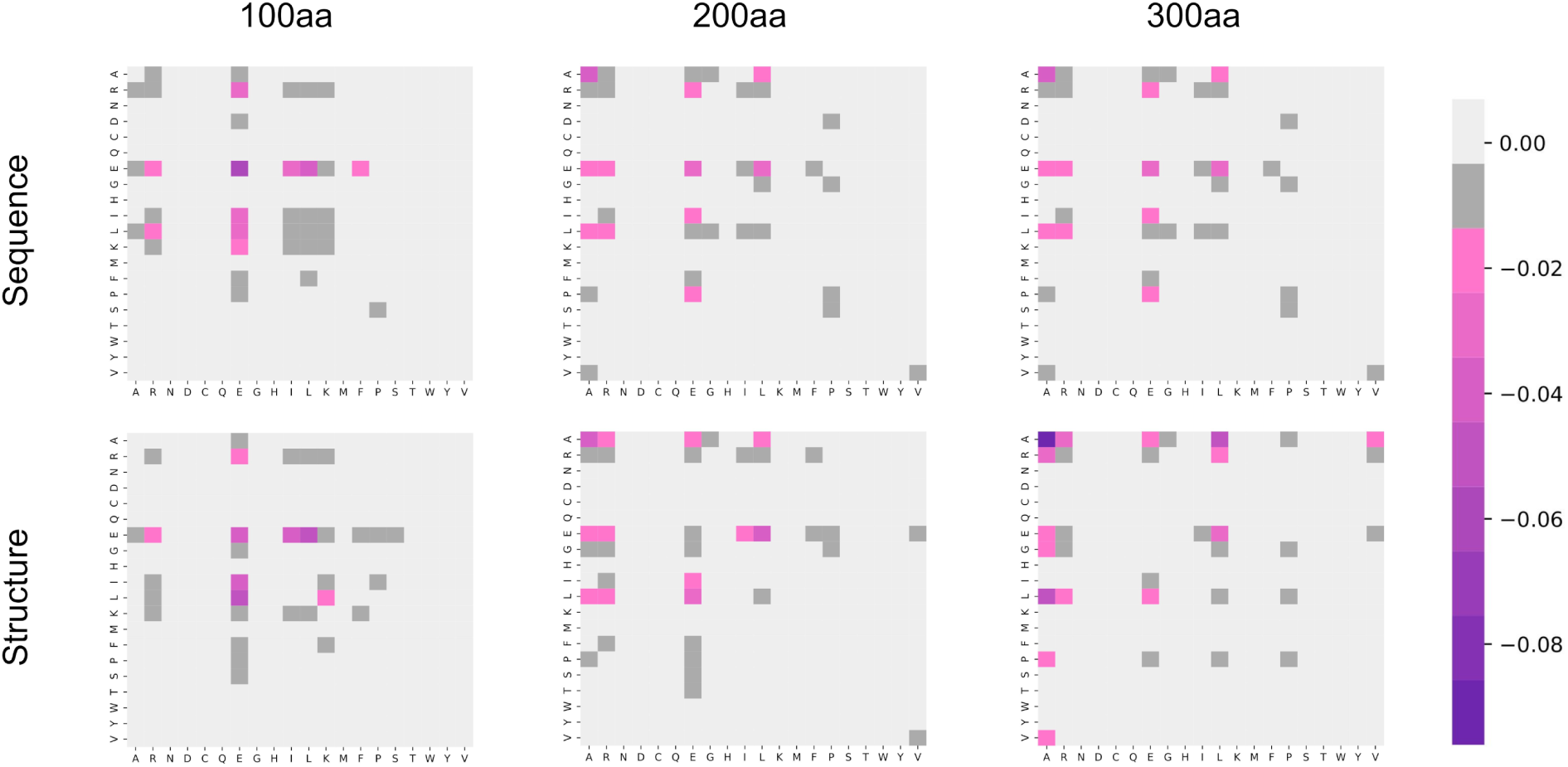
Sampling from different noise distributions generates proteins with different sequence and structural neighbors. Frequencies of amino acid neighbors in generated sequences (top row) and nearest structure neighbors (bottom row). Values are calculated as the difference between native sequences and unconditionally generated sequences for 100AA (left), 200AA (middle) and 300AA (right). Unconditional designs of 200AA and 300AA are characterized by more frequent alanine-leucine, alanine-glutamic acid, and alanine-alanine sequence and structure contacts. In structure space unconditional proteins exhibit lower frequencies of glutamic acid-glutamic acid and glutamic acid-proline neighbors.

**Figure S5:**
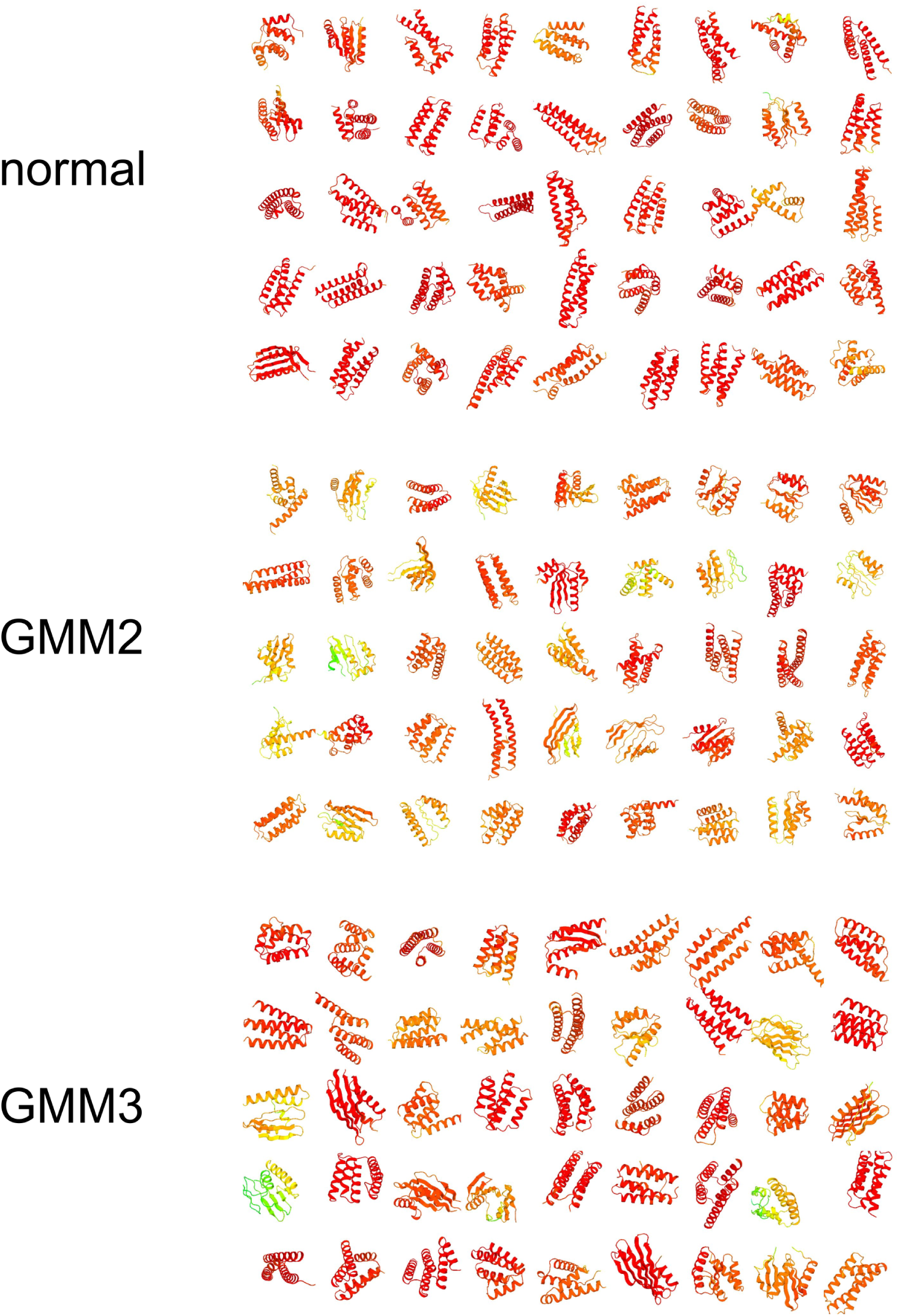
Sampling from different noise distributions generates proteins with more diverse secondary structure. Representative 100AA unfiltered and unconditionally generated proteins from normal distribution, GMM2, and GMM3. Colored by model pLDDT (red → high confidence).

**Figure S6:**
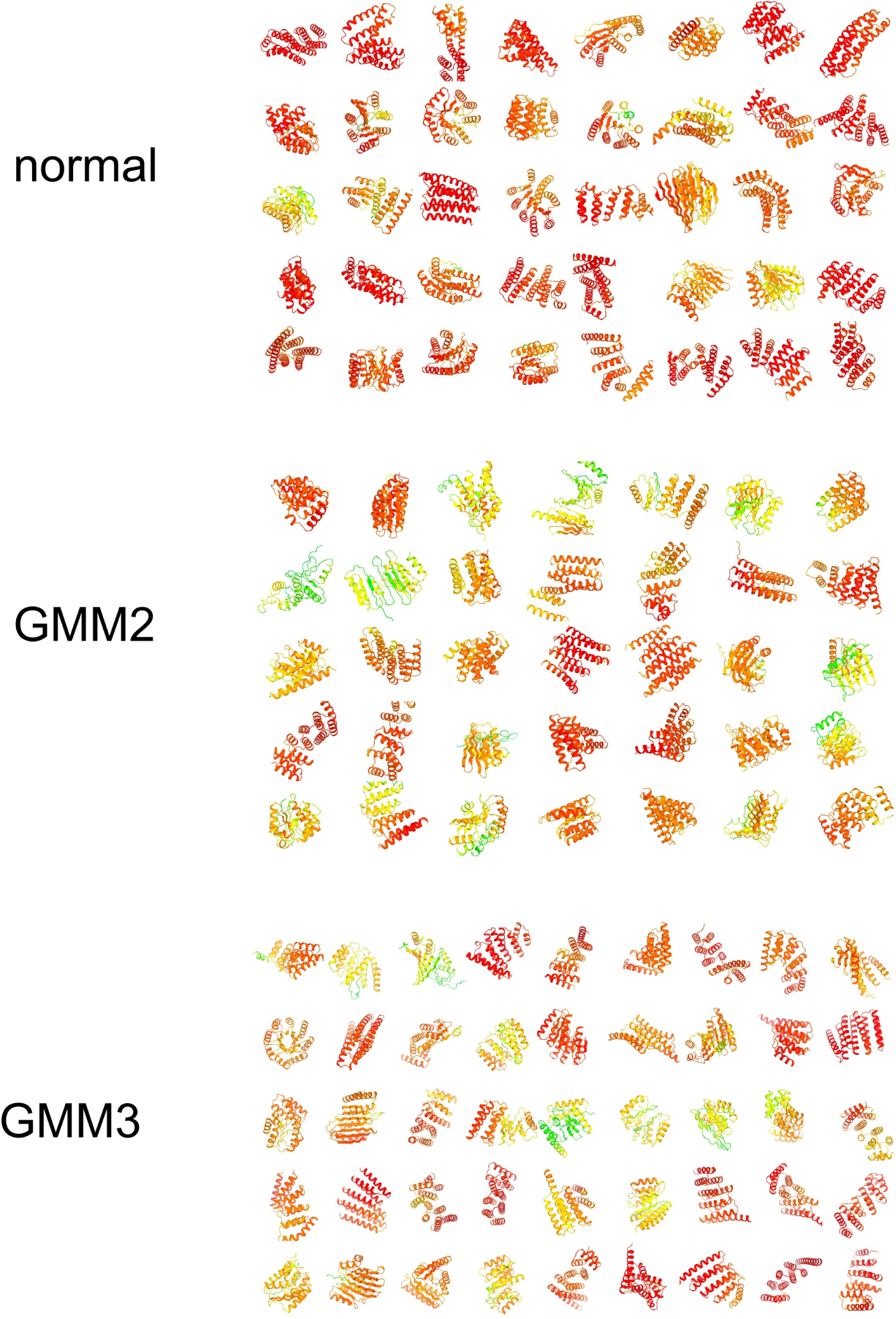
Sampling from different noise distributions generates proteins with more diverse secondary structure. Representative 200AA unfiltered and unconditionally generated proteins from normal distribution, GMM2, and GMM3. Colored by model pLDDT (red → high confidence).

**Figure S7:**
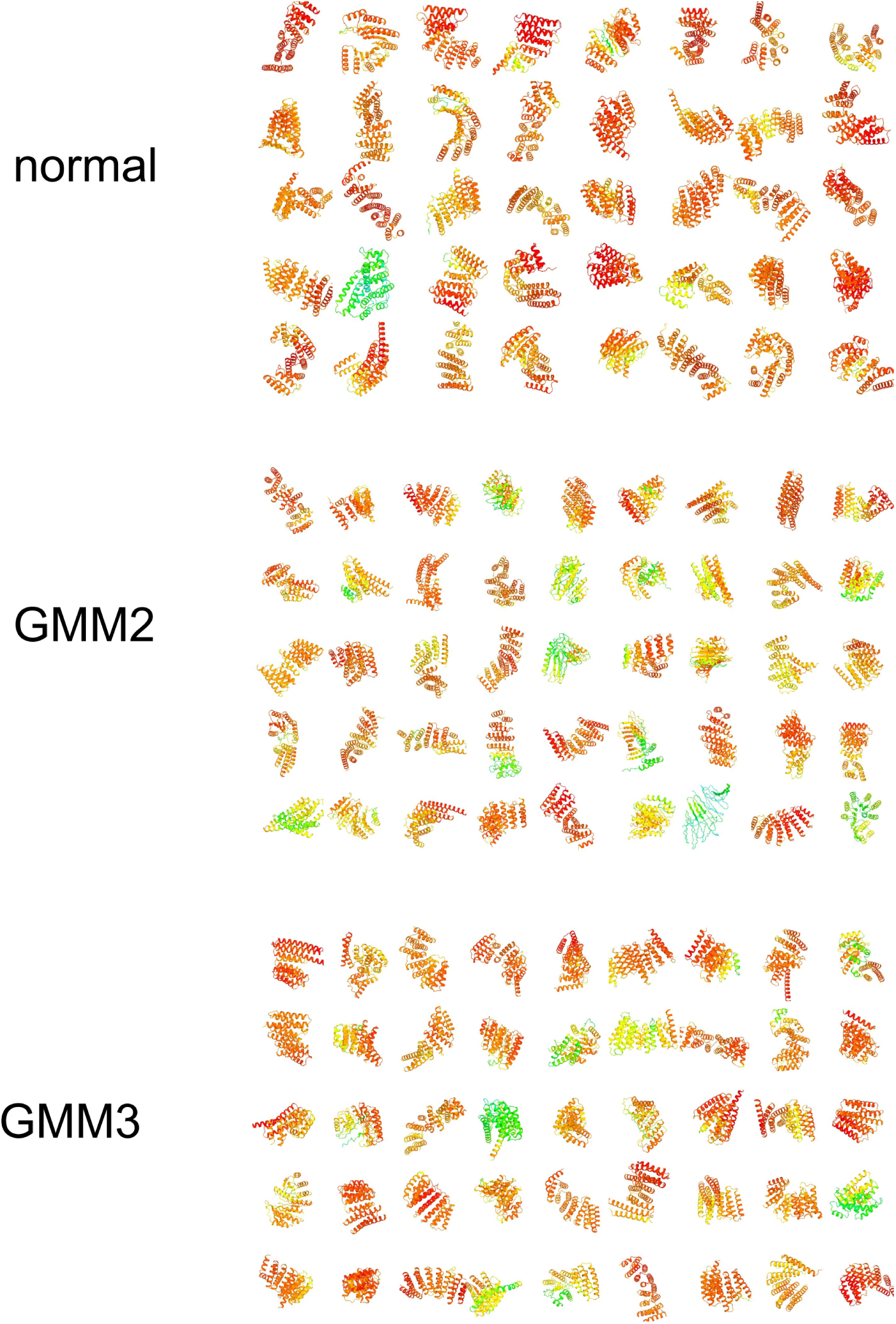
Sampling from different noise distributions generates proteins with more diverse secondary structure. Representative 300AA unfiltered and unconditionally generated proteins from normal distribution, GMM2, and GMM3. Colored by model pLDDT (red → high confidence).

**Figure S8:**
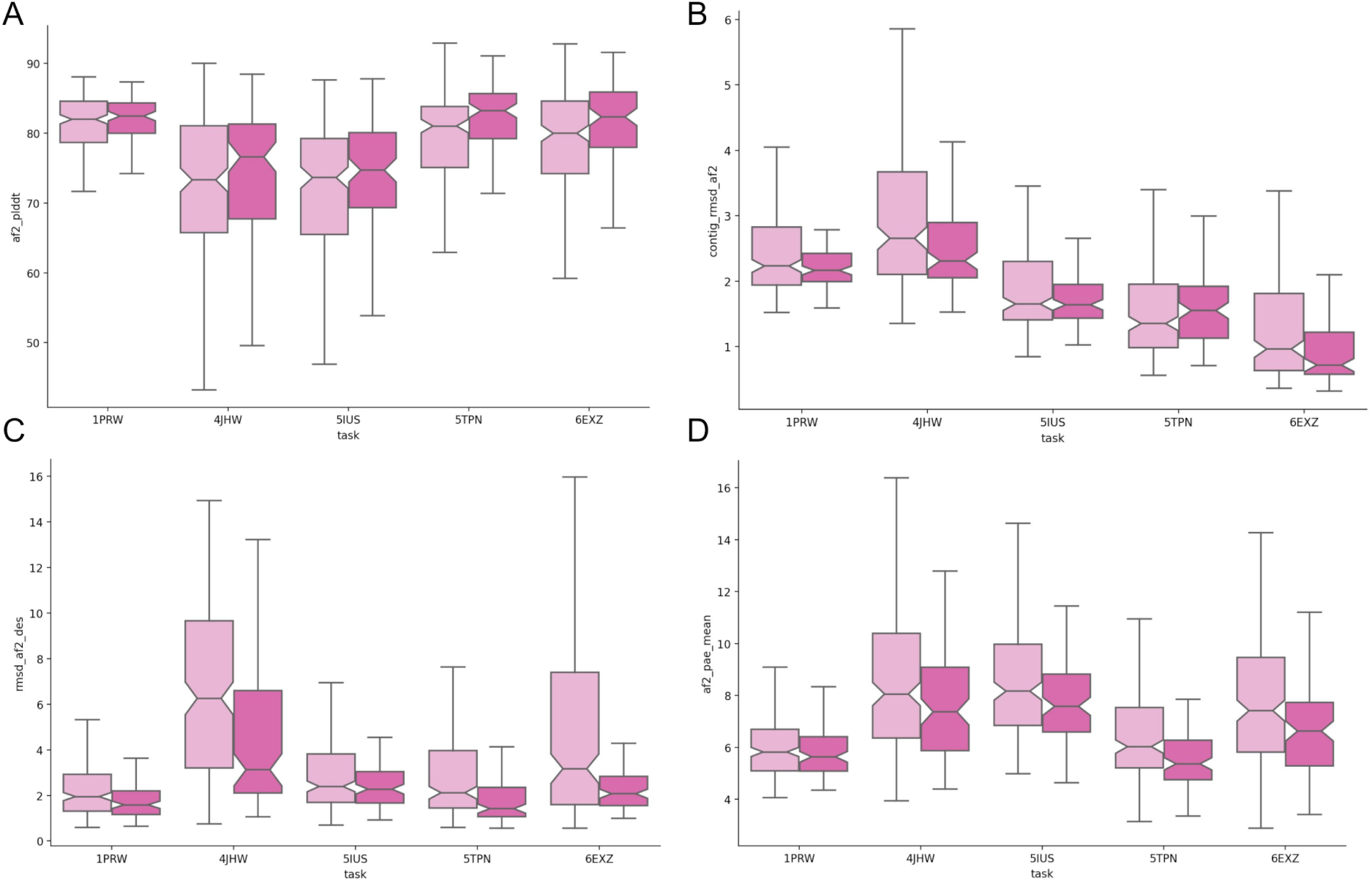
AF2 metrics for scaffolding of structure-sequence motifs in the PDB IDs listed in 25 (light pink) and 100 (dark pink) time steps. (A) AF2 pLDDT for designs, (B) RMSD of motif predicted by AF2 to design, (C) RMSD of AF2 to design for whole structure, (D) predicted alignmed error (pAE) of designs from AF2. The following contig arguments were used to run motif scaffolding benchmark: 1PRW- contigs 8-20,A21-31,16-25,A56-67,8-20, 6EXZ - contigs 0-95,A28-42,0-95, 5TPN - contigs 10-40,A163-181,10-40, 5IUS - contigs 0-30,A119-140,15-40,A63-82,0-30, 4JHW - contigs 0-25,F196-212,15-30,F63-69,10-25

**Figure S9:**
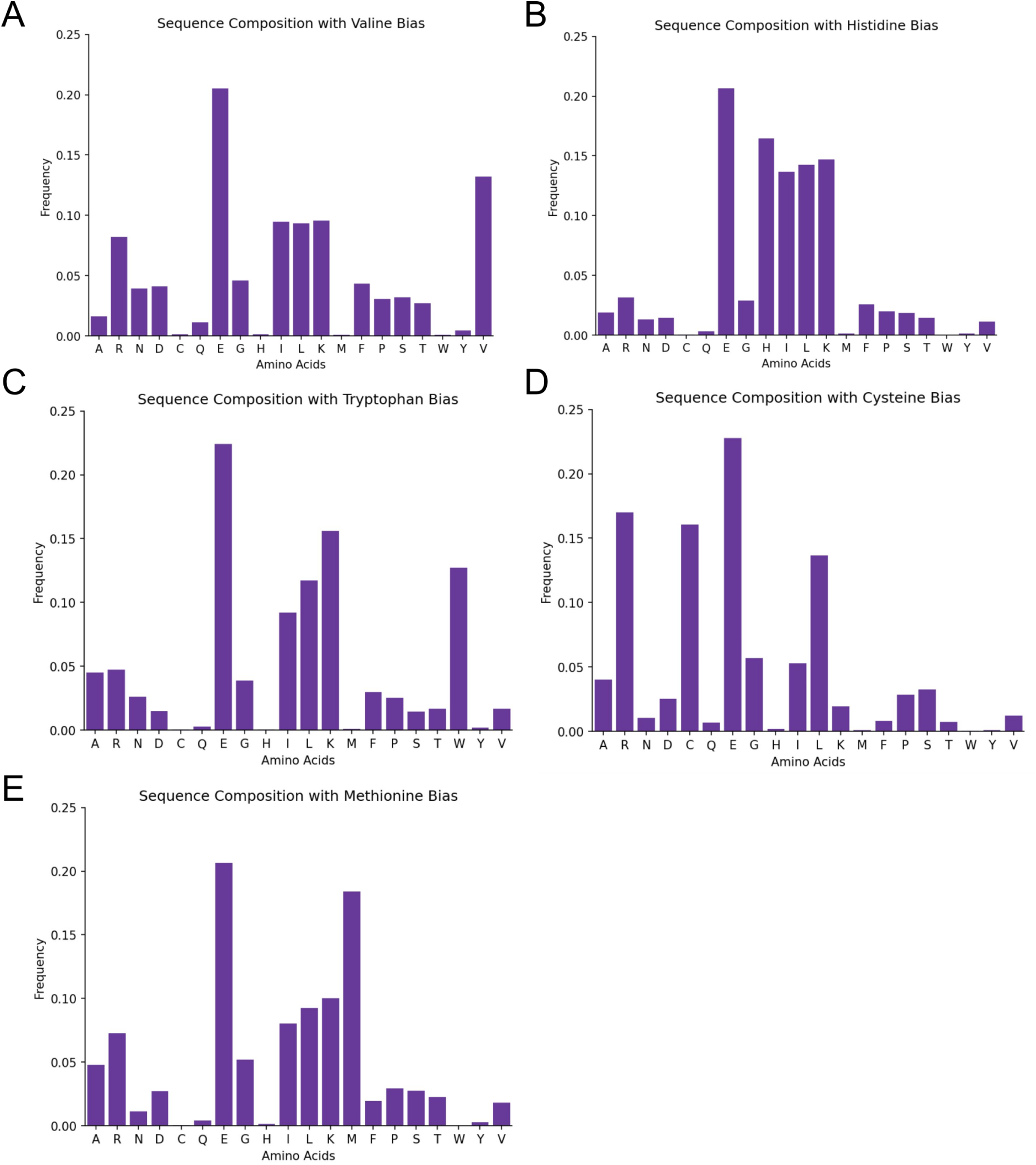
Biasing for specific amino acids results in increased frequency of the specified residue in generated proteins. Amino acid distributions of proteins generated with amino acid compositional bias for (A) valine, (B) histidine, (C) tryptophan, (D) cysteine, and (E) methionine.

**Figure S10:**
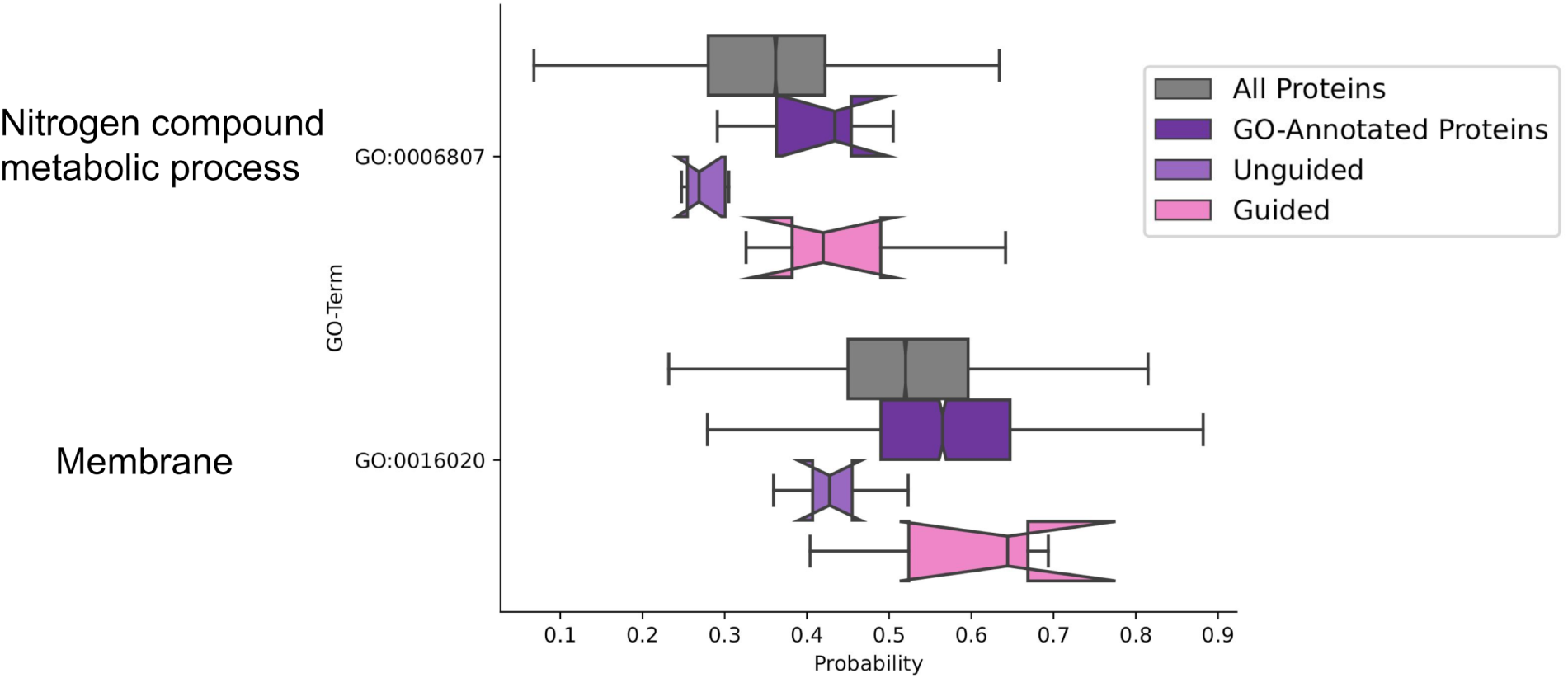
GO-guidance. The network has been guided with the DeepGOPlus Gene Ontology (GO) classifier to generate proteins with specific characteristics and functions. Exemplary, the classifier GO probability scores for all UniProt proteins, all proteins annotated with the chosen GO term, unconditionally unguided proteins generated with our model and guided proteins generated with our models for the GO terms nitrogen compound metabolic process (GO:0006807) and membrane (GO:0016020) are shown. The classifier has a high false positive rate due to a high mean probability as well as for all UniProt proteins including proteins not annotated with this specific GO term. For both GO terms a shift in the probabilities can be shown for guided proteins in comparison to unguided proteins.

**Figure S11:**
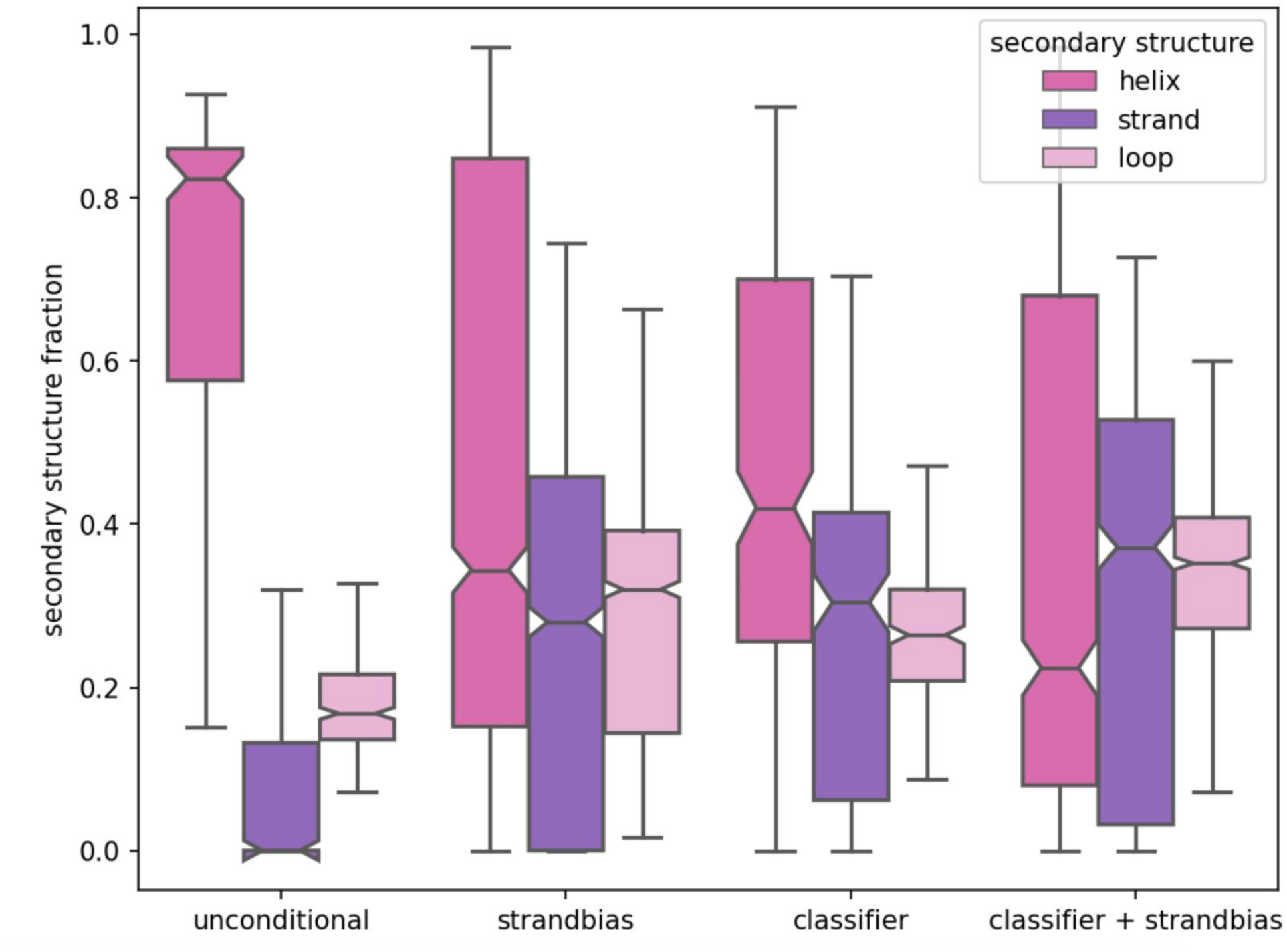
Secondary structure composition comparison when generation unconditional designs, designs with strand bias, designs with classifier guidance, and combination of classifier guidance and strand bias.

**Figure S12:**
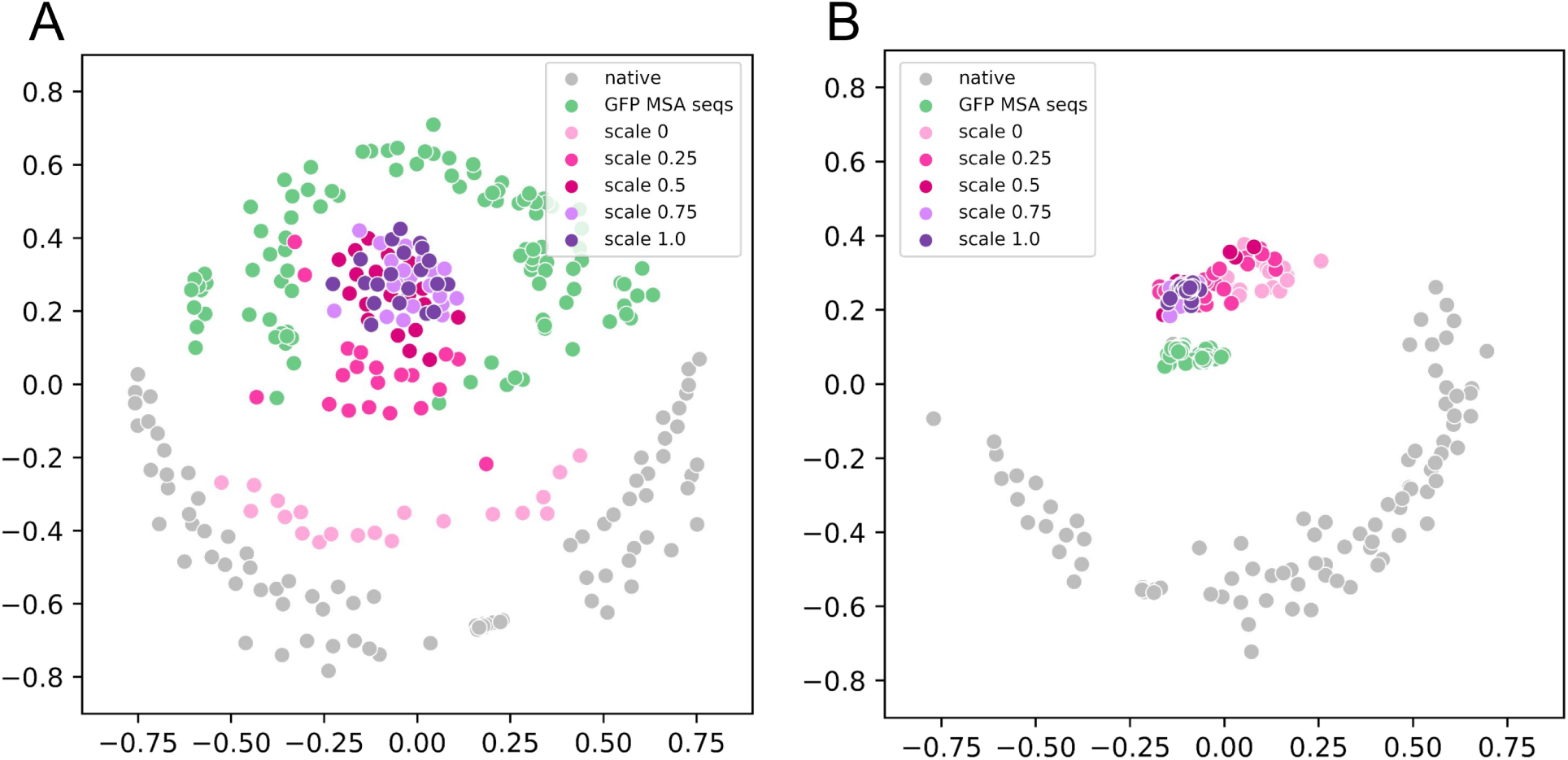
Multidimensional scaling plots of proteins generated with increasing GFP PSSM guidance scales. (A) Higher PSSM scaling increases sequence clustering to native GFPs. Distance metric is percent sequence identity. Green dots are native GFP sequences derived from a GFP MSA with sequence identity cutoffs 30-90% to the query sequence. Grey are randomly sampled native sequences from Uniprot90 (B) Low PSSM scaling results in increased structural diversity and samples of more diverse beta barrels. Higher PSSM scaling reduces structural diversity and clusters closer to native GFPs. Distance metric is TM score. Green dots are structures derived from the same MSA as (A) and grey dots are structures derived from the same set as (A).

**Figure S13:**
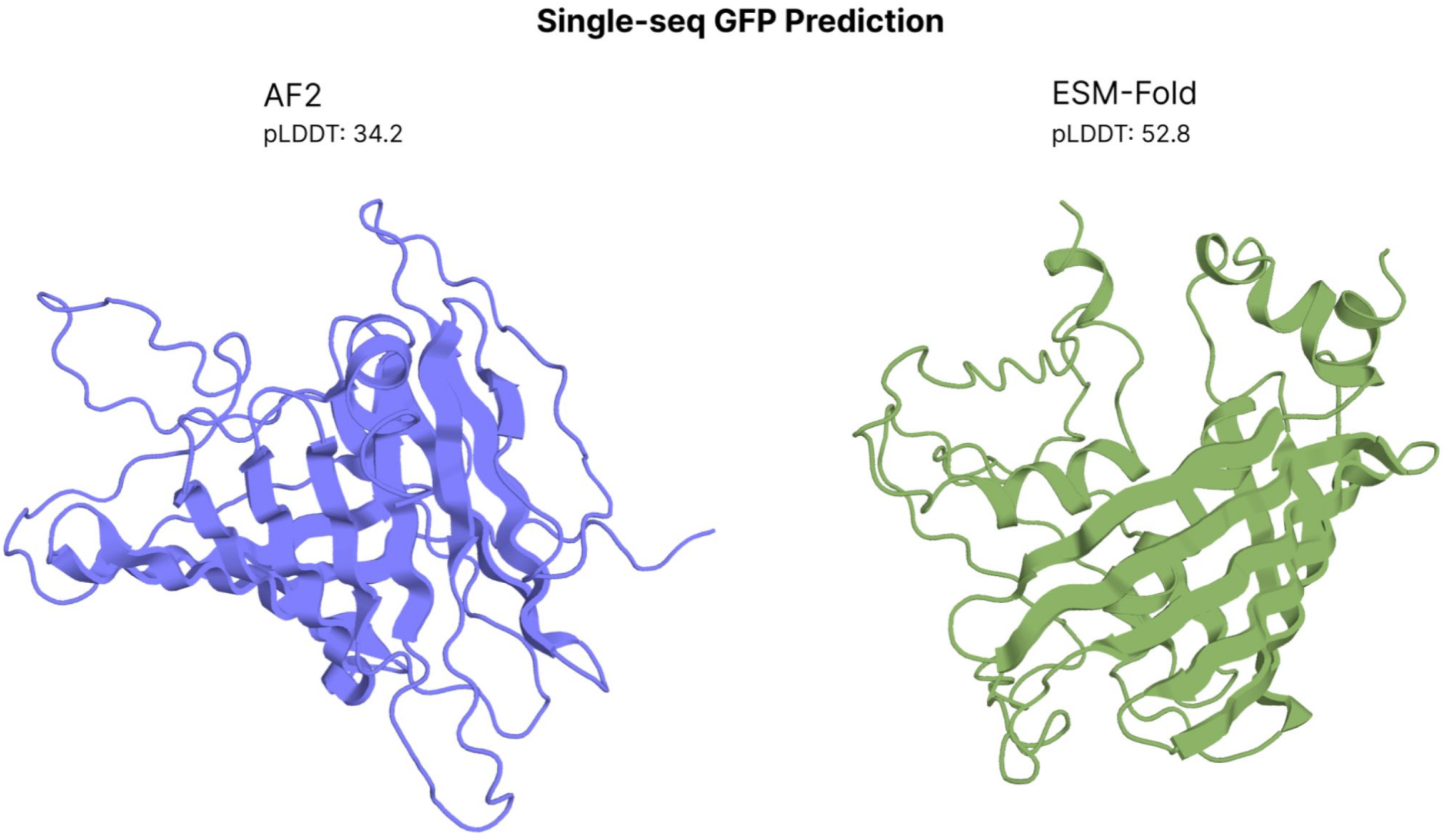
Single-sequence structure predictions of Green Fluorescent Protein (GFP), (PDB 1EMA). AlphaFold2 (left) and ESM-Fold (right) predictions fail to recover the tertiary structure of GFP when run in single-sequence mode. Both models return structures with low confidence (pLDDT). Models were run with 6 recycles.

**Figure S14:**
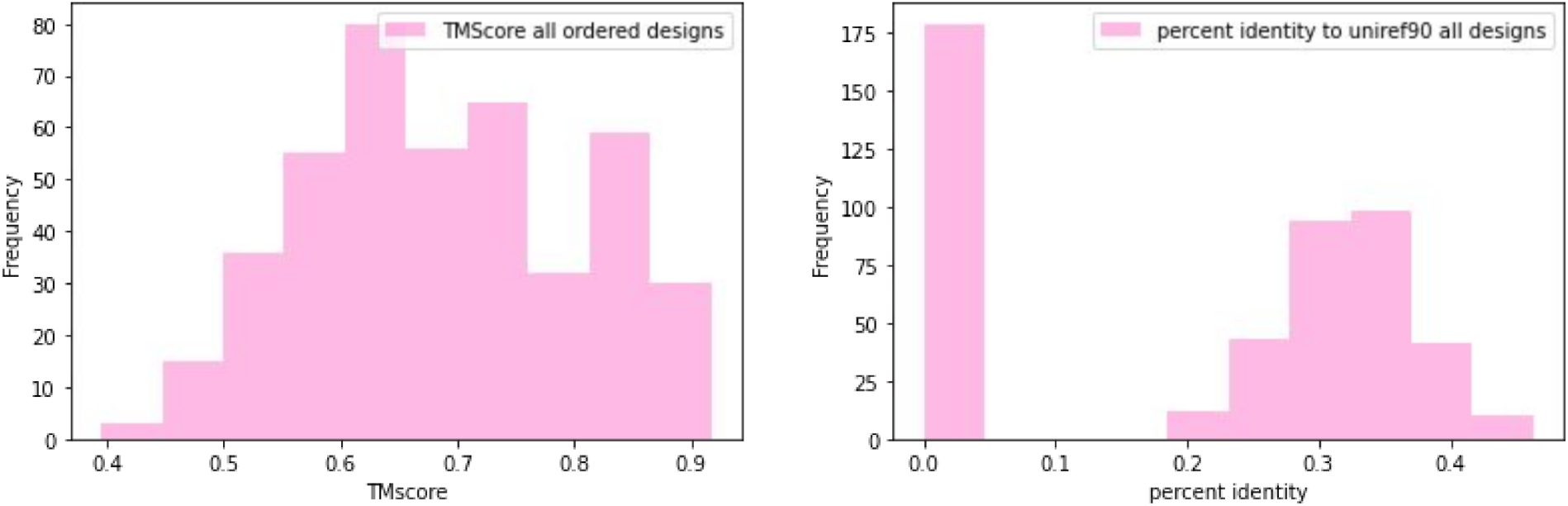
TMScore (left) and sequence identity (right) distributions of all ordered designs against PDB.

**Pseudocode S1:**
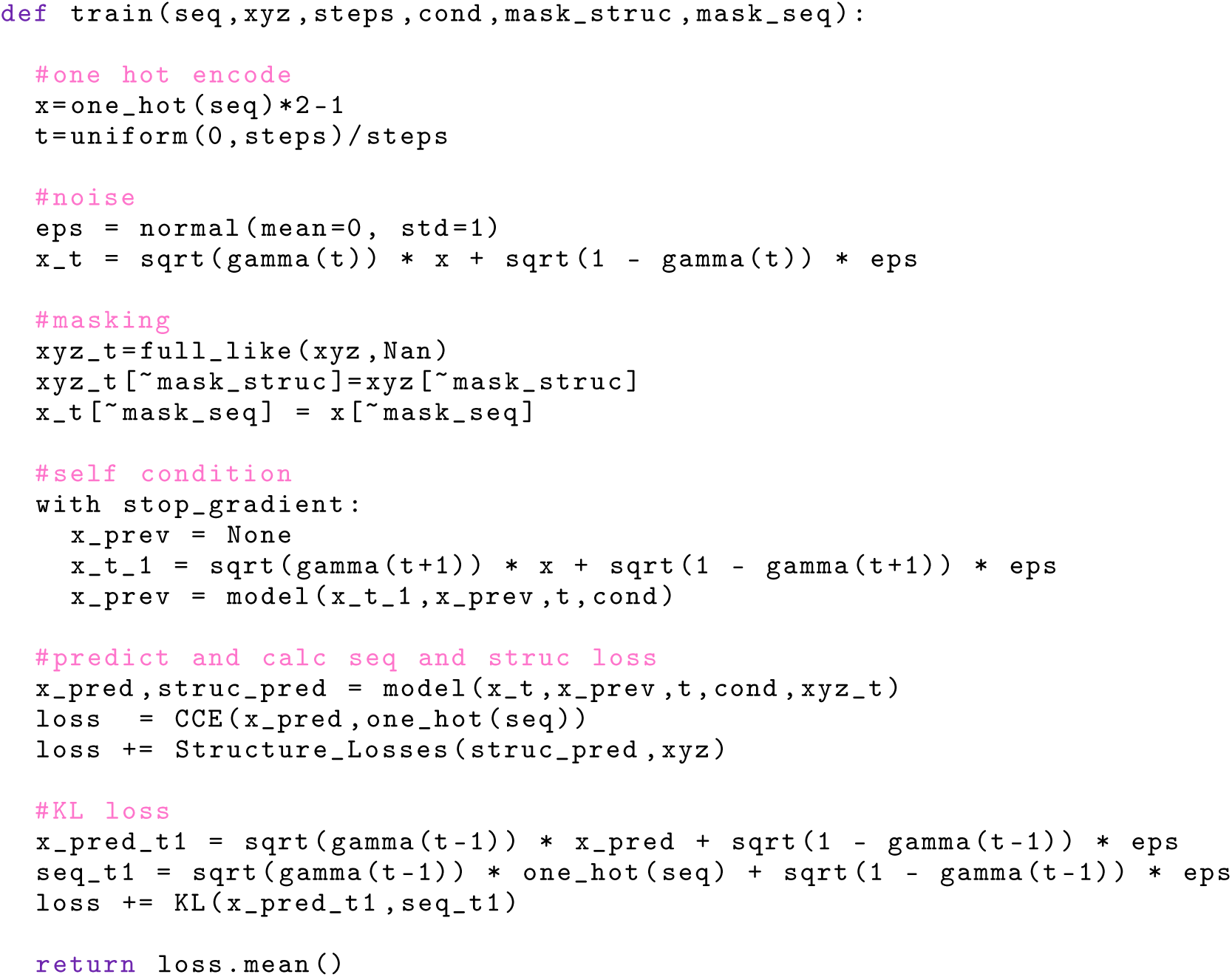
Training.

**Pseudocode S2:**
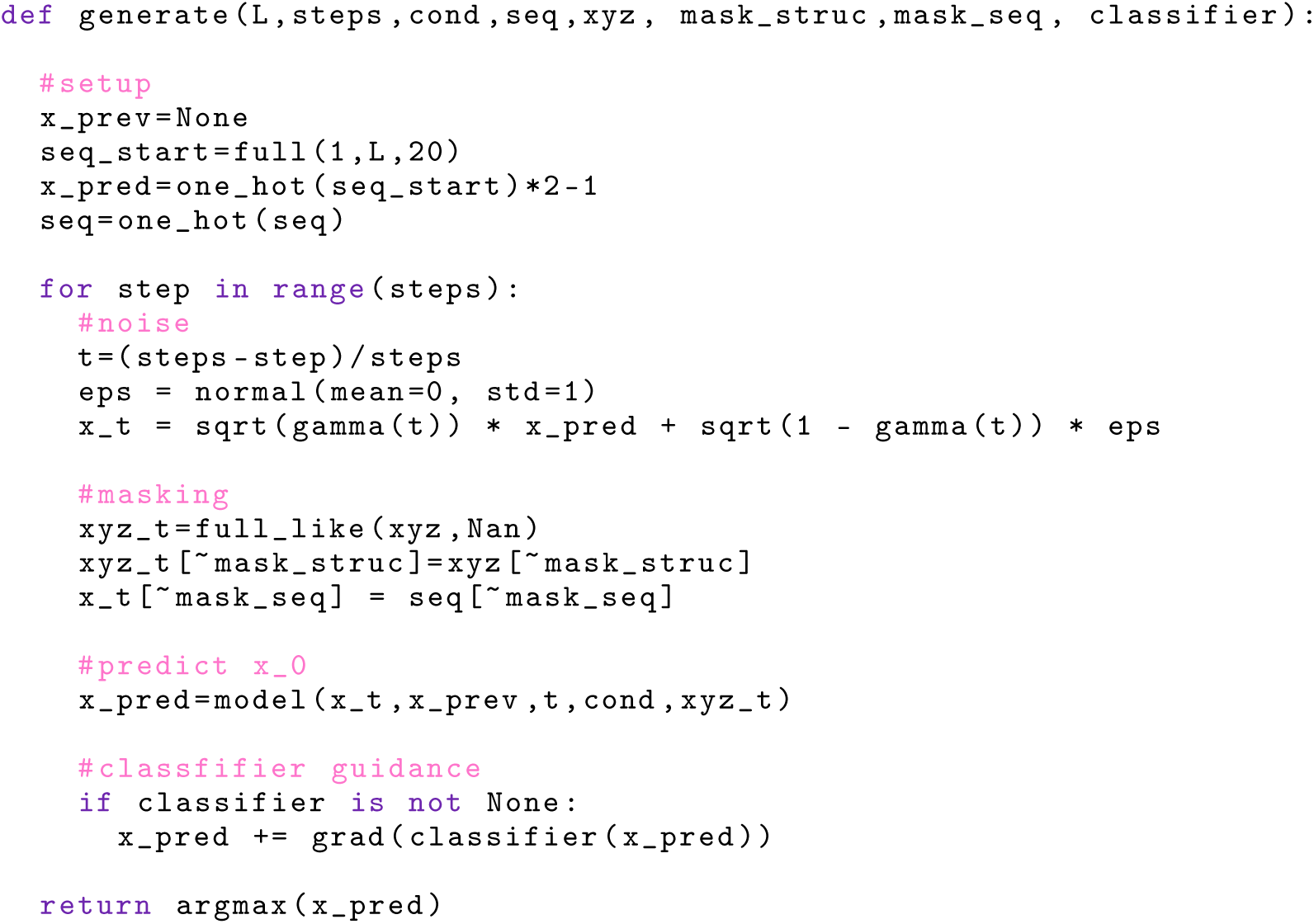
Inference.

**Pseudocode S3:**
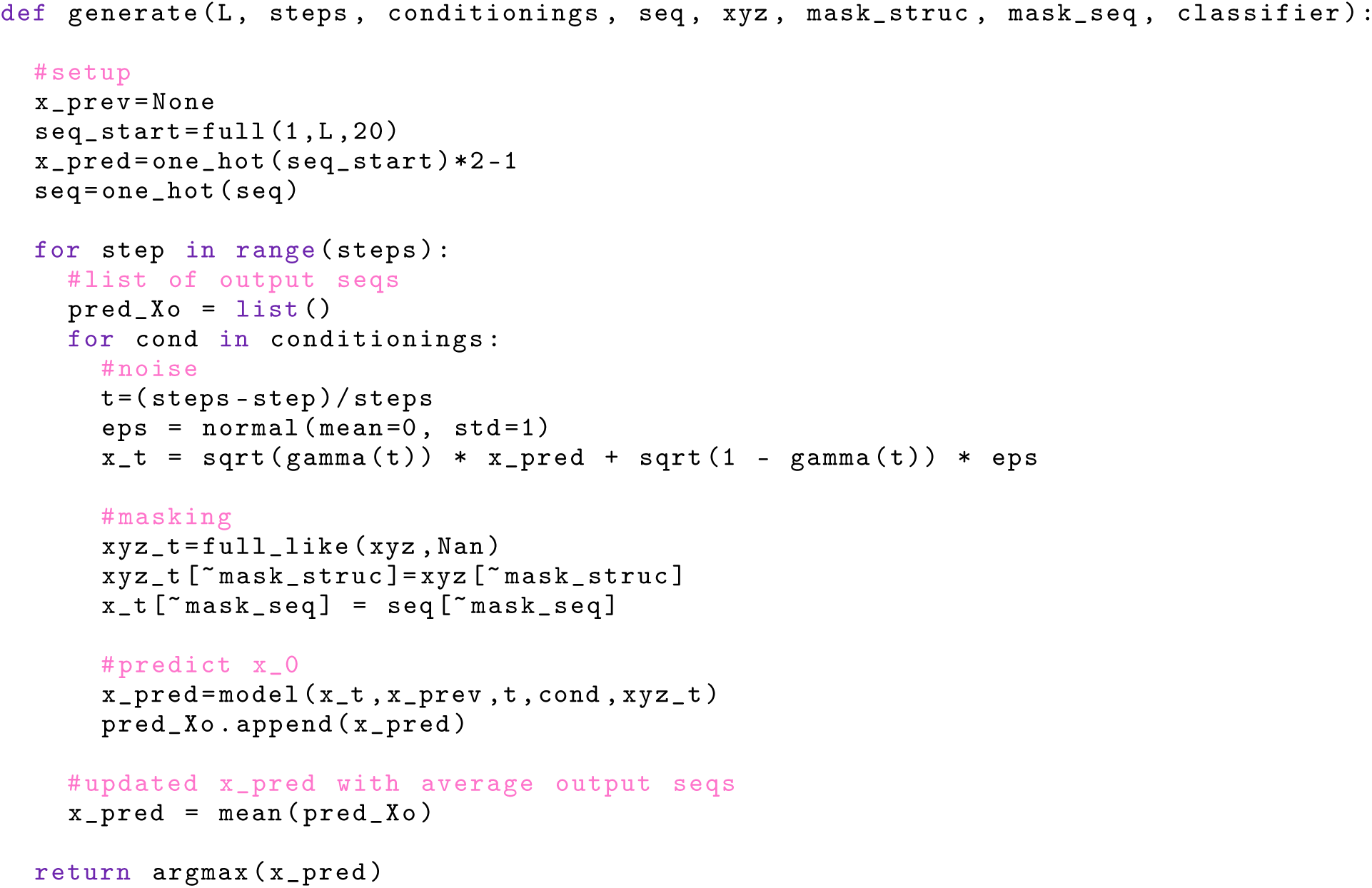
Multistate design.

**Pseudocode S4:**
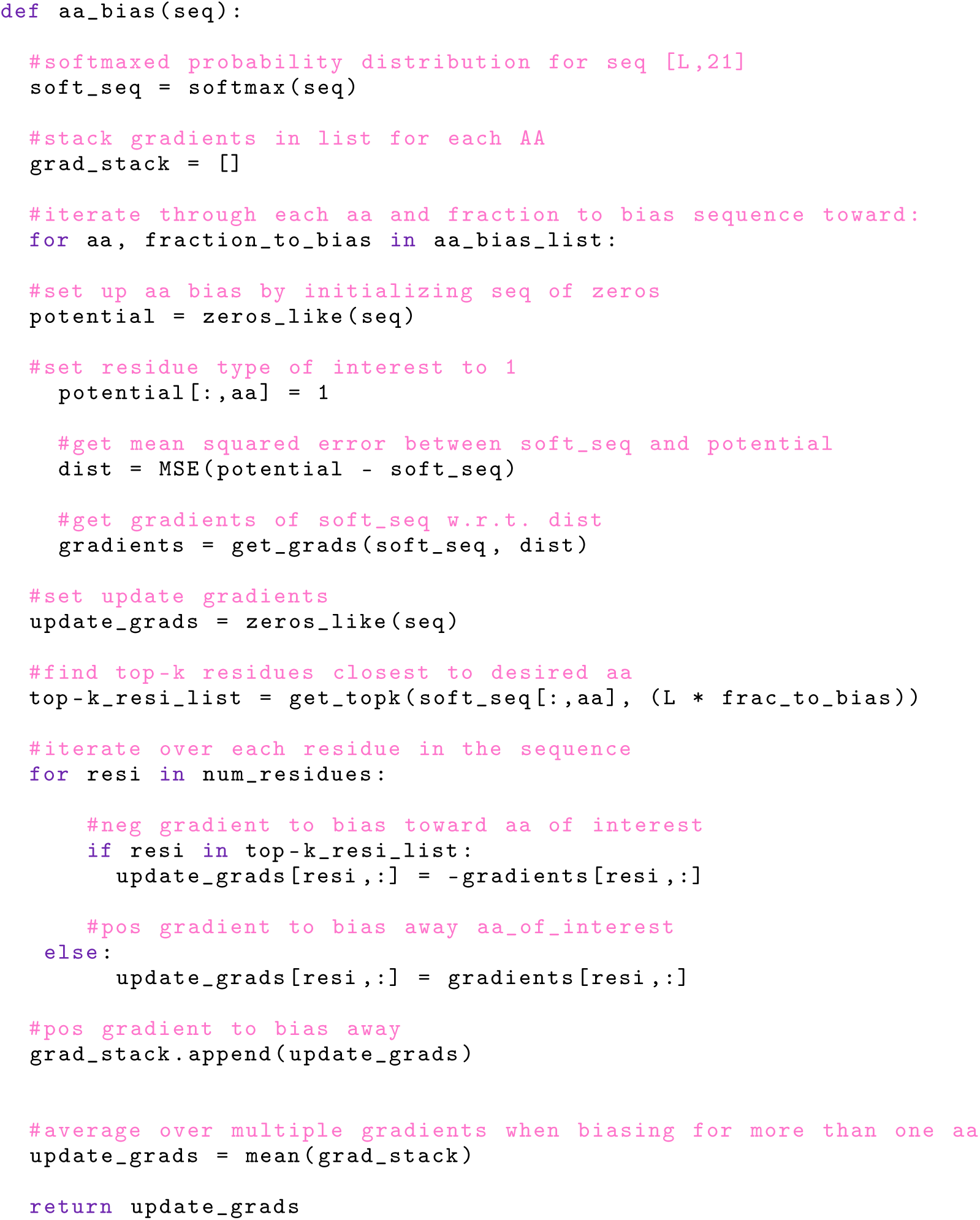
Amino acid composition potential.

**Pseudocode S5A:**
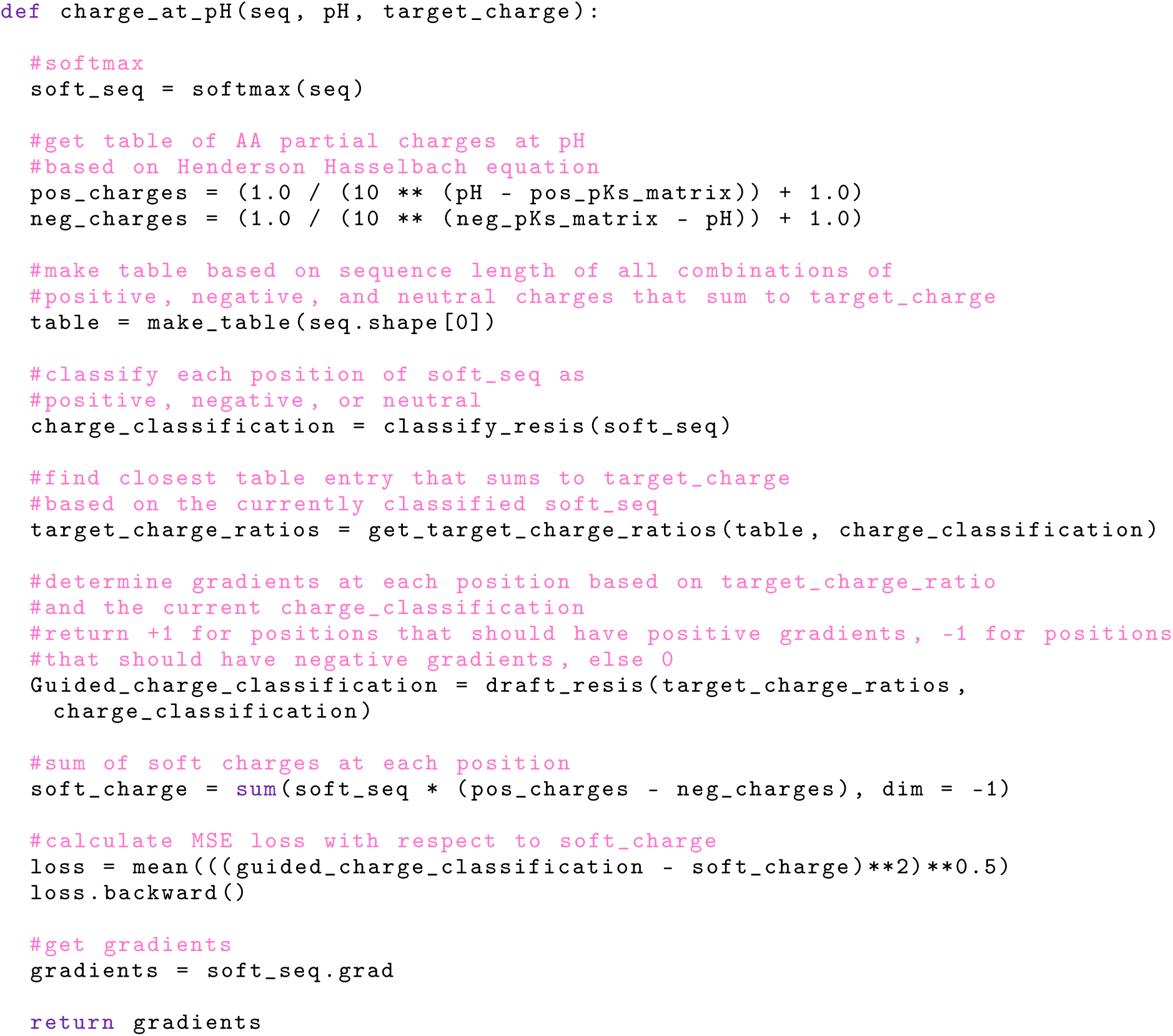
Net charge potential.

**Pseudocode S5B:**
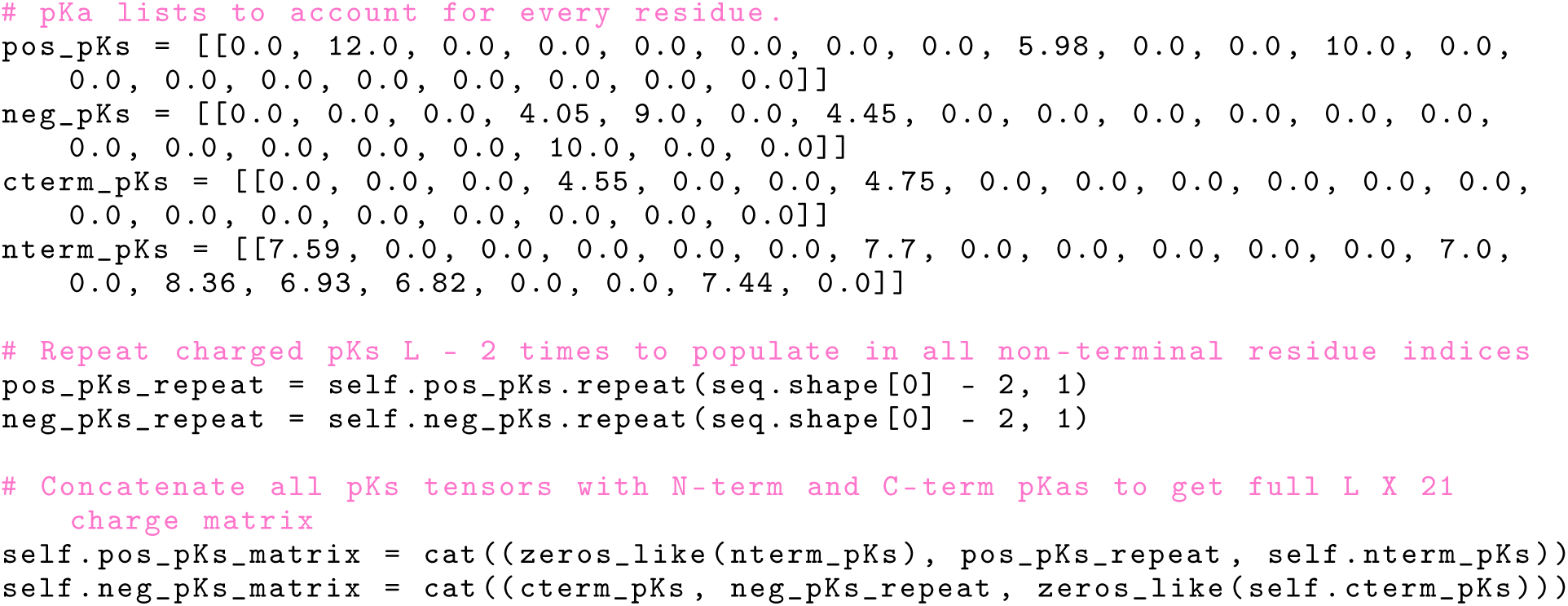
Net charge potential data structures and tables.

**Pseudocode S6A:**
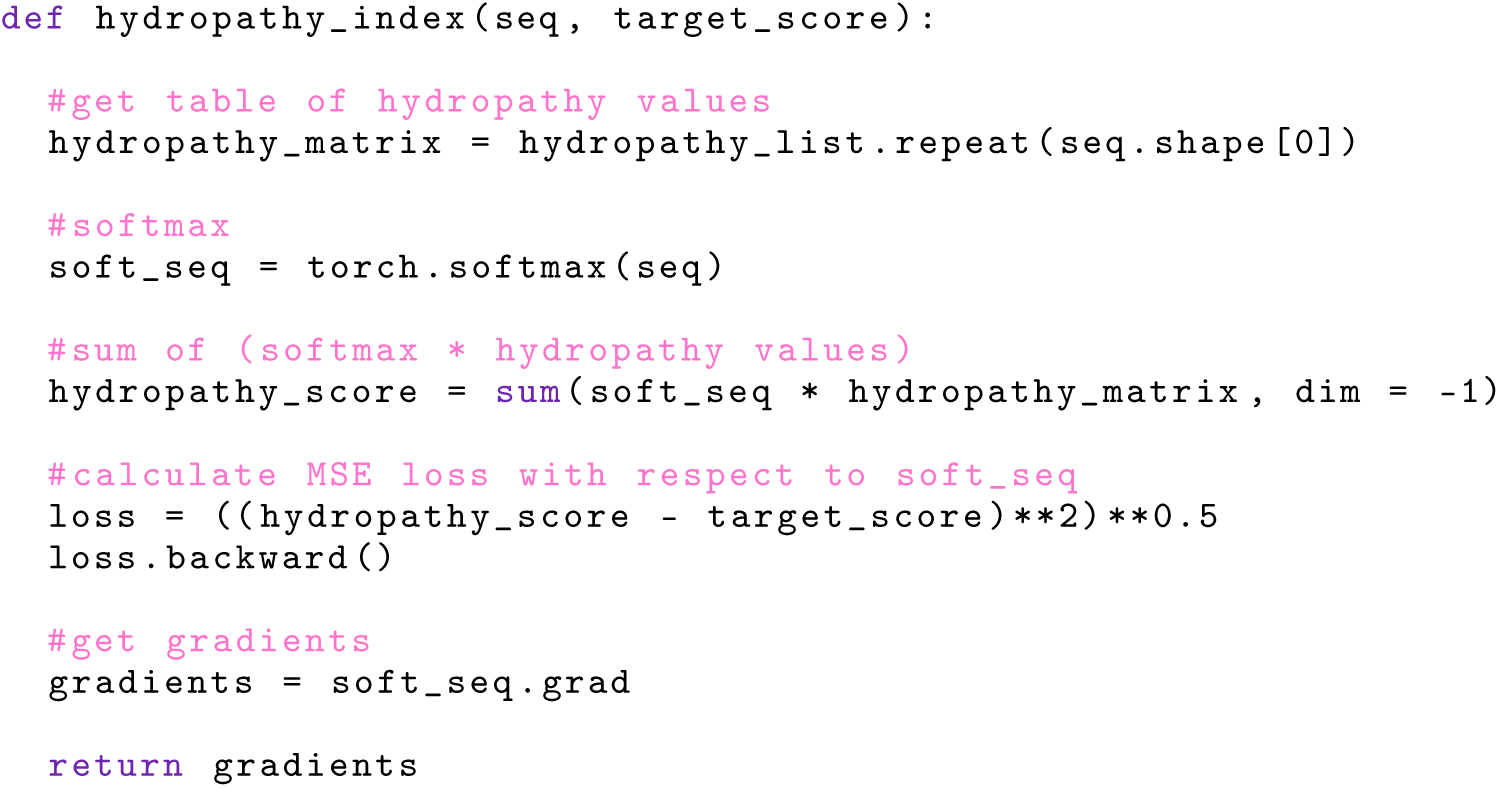
Hydrophobicity potential.

**Pseudocode S6B:**
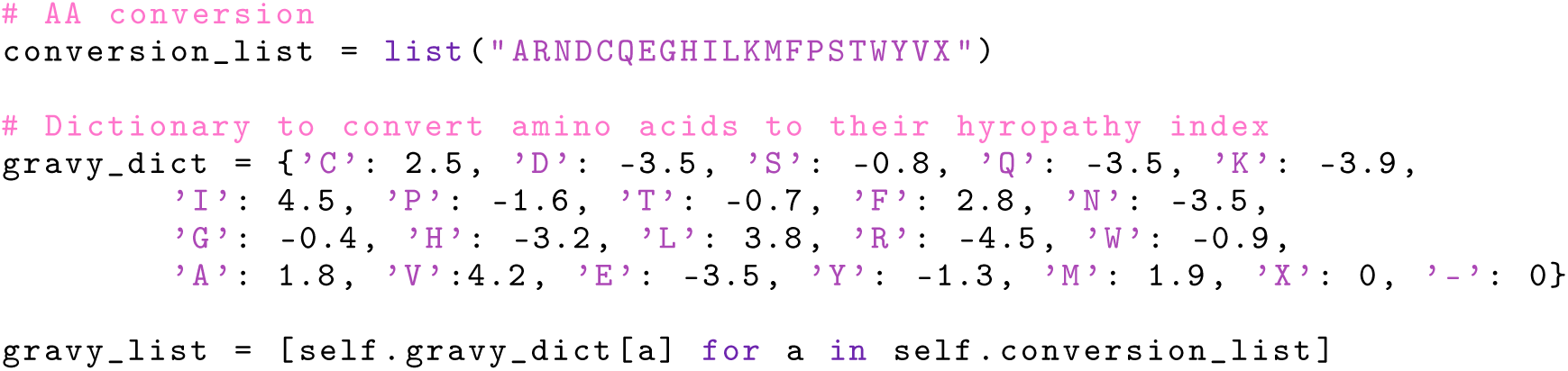
Hydrophobicity potential data structures and tables.

**Table S1:** Observed and predicted mass for designs used in mass spectrometry experiments.

| **Design** | **Observed**<br>**Mass (Da)** | **Predicted**<br>**Mass (Da)** | **Difference** |
| --- | --- | --- | --- |
| TrpA5 | 11955 | 12086.18 | -131.18 |
| TrpA9 | 11592 | 11722.61 | -130.61 |
| TrpB9 | 11312 | 11765.34 | -453.34 |
| ValF3 | 10470 | 10600.58 | -130.58 |
| ValF12 | 11203 | 11333.68 | -130.68 |
| ValF11 | 10850 | 10981.23 | -131.23 |
| UncC7 | 11690 | 11820.99 | -130.99 |
| UncC4 | 10851 | 10981.47 | -130.47 |
| UncD5 | 11191 | 11321.68 | -130.67 |
| RCC12 | 23072 | 23202.83 | -130.83 |
| RCE4 | 22880 | 23010.88 | -130.88 |
| RCE8 | 22575 | 22705.43 | -130.43 |
| RCF12 | 22997 | 22996.3 | 0.7 |
| NCG1 | 21162 | 21292.13 | -130.13 |
| NCA1 | 20779 | 20910.78 | -131.78 |
| CysG4 | 8427 | 8561.43 | -134.43 |
| CysH6 | 8675 | 8813.75 | -138.75 |
| CysH11 | 8633 | 8642.57 | -9.57 |
| TEV A1 | 14905 | 15036 | -131 |
| TEV A2 | 14713 | 14844 | -131 |
| TEV A3 | 14005 | 14136 | -131 |
| TEV A4 | 14044 | 14175 | -131 |
| TEV A5 | 14265 | 14396 | -131 |
| TEV A6 | 14178 | 14309 | -131 |
| TEV A7 | 14714 | 14845 | -131 |
| FUR A9 | 22894 | 23025 | -131 |
| FUR A10 | 23556 | 23687 | -131 |
| TEV B1 | 14864 | 14995 | -131 |
| TEV B2 | 14312 | 14443 | -131 |
| TEV B3 | 14561 | 14692 | -131 |
| TEV B6 | 14547 | 14661 | -114 |
| FUR B10 | 19733 | 19916 | -183 |
| FUR C2 | 18934 | 19065 | -131 |
| FUR D2 | 18853 | 18984 | -131 |
| FUR D3 | 18869 | 19000 | -131 |

**Table S2:** Mass spec data of experimentally validated cysteine-rich designs. The mass of each design is reported in the presence and absence of the reducing agent TCEP. The mass difference between reduced and non-reduced designs is used to calculate the n<u>umber of disulfides formed and compared to the number of designed dis</u>ulfides.

| Design ID | Mass with MET loss (Da) |  |  | # Disulfides formed |
| --- | --- | --- | --- | --- |
|  | + TCEP | - TCEP | Difference |  |
| cys_G4 | 8431 | 8425 | 6 | 3/3 |
| cys_G6 | 8778 | 8770 | 8 | 4/4 |
| cys_G10 | 8695 | 8687 | 8 | 4/4 |
| cys_G11 | 8368 | 8363 | 5 | 3/3 |
| cys_H6 | 8683 | 8675 | 8 | 4/5 |

**Table S3:** Secondary structure prediction of CD data (200-250 nm) of designs RC_E8 Fig 3E middle top, and RC_F11 Fig 3E middle bottom with BeStS<u>el server indicating high percentage of beta cont</u>ent.

|  | CD RC_E8 | CD RC_F11 |
| --- | --- | --- |
| **Helix** | 1.0 | 2.7 |
| regular | 0.2 | 1.2 |
| distorted | 0.8 | 1.5 |
| **Antiparallel** | 40.7 | 26.8 |
| left-twisted | 0.0 | 0.0 |
| relaxed | 24.3 | 12.2 |
| right-twisted | 16.4 | 14.6 |
| **Parallel** | 0.0 | 0.0 |
| **Turn** | 14.0 | 17.9 |
| **Others** | 44.3 | 52.7 |

**Table S4:**
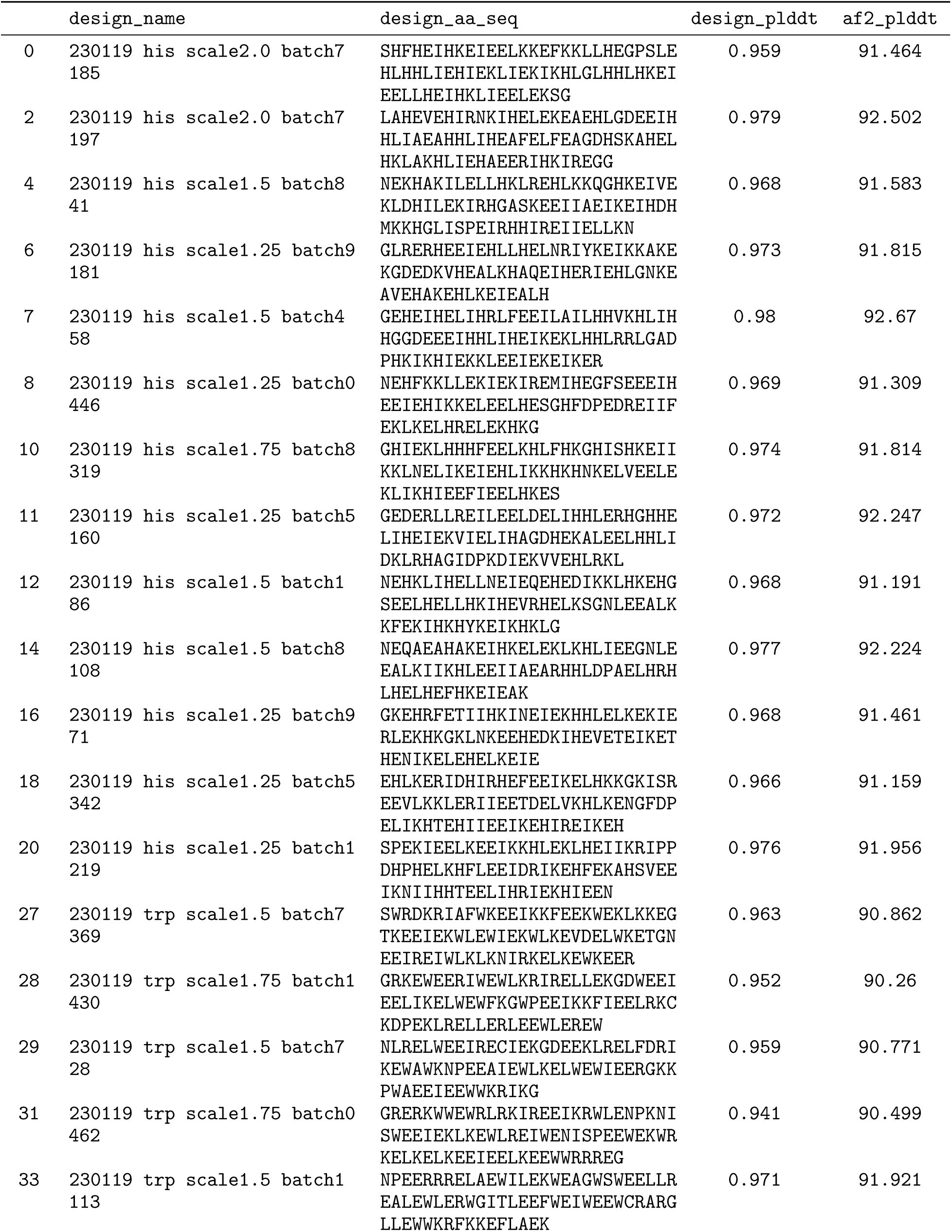
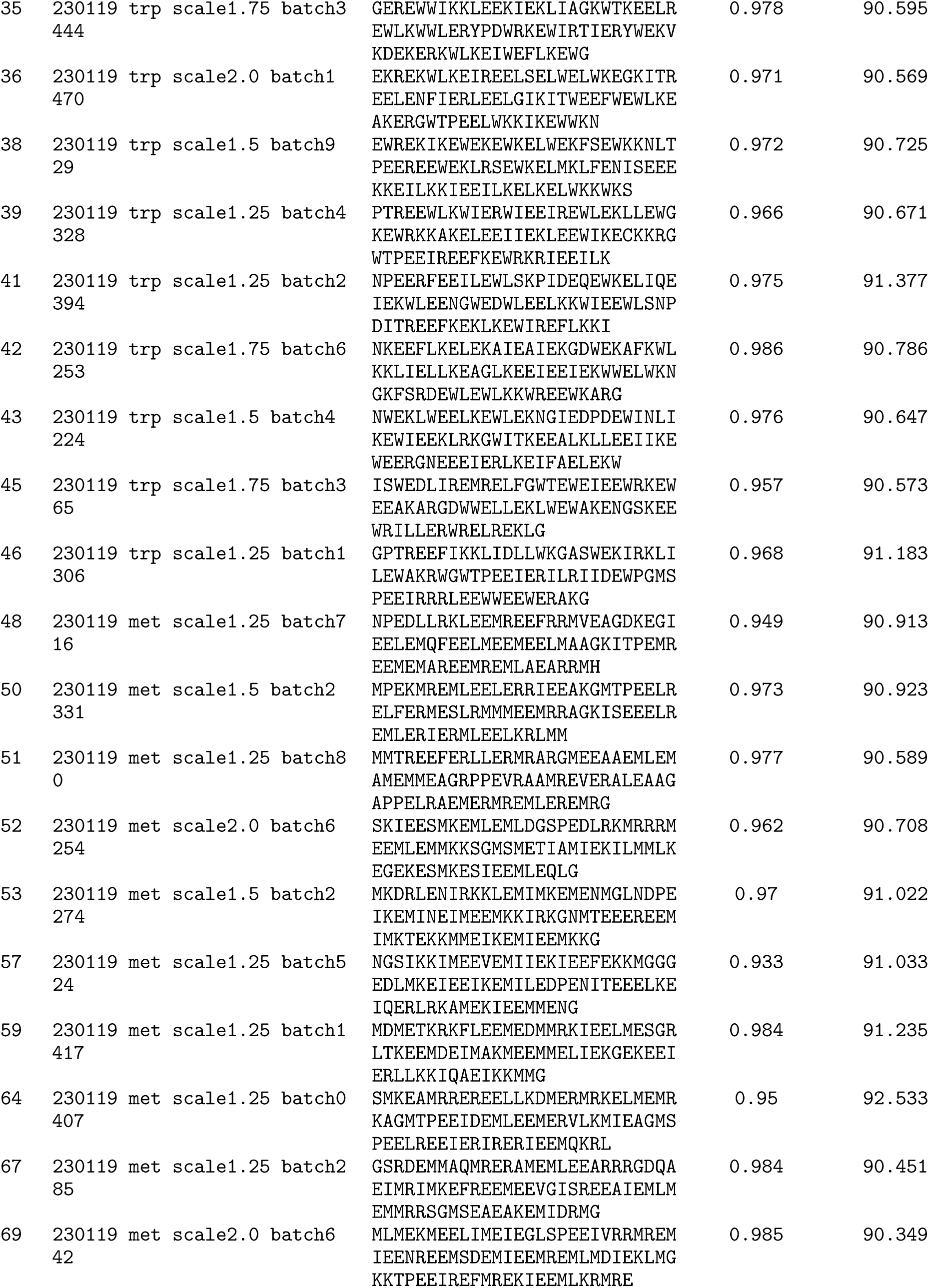
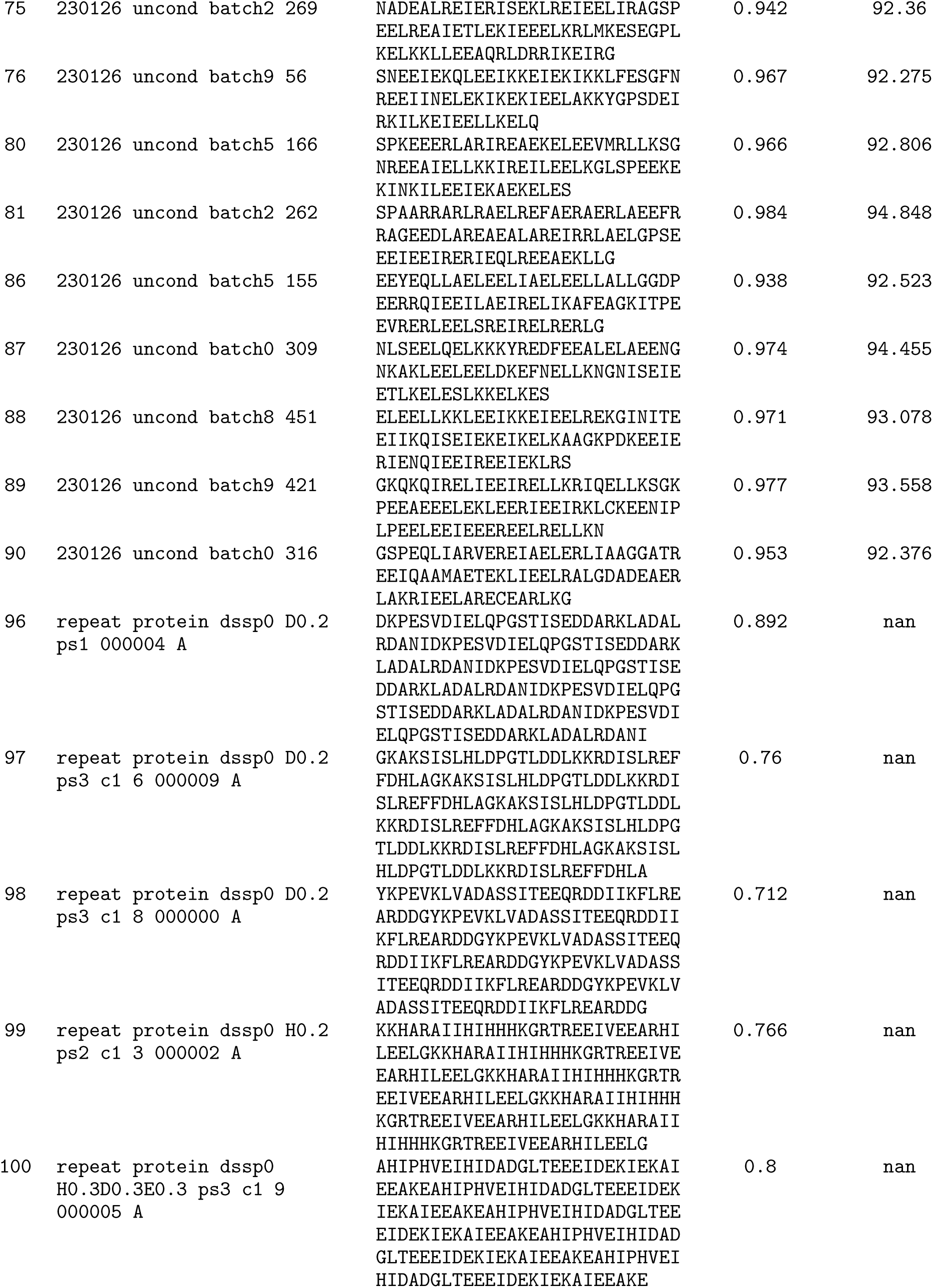
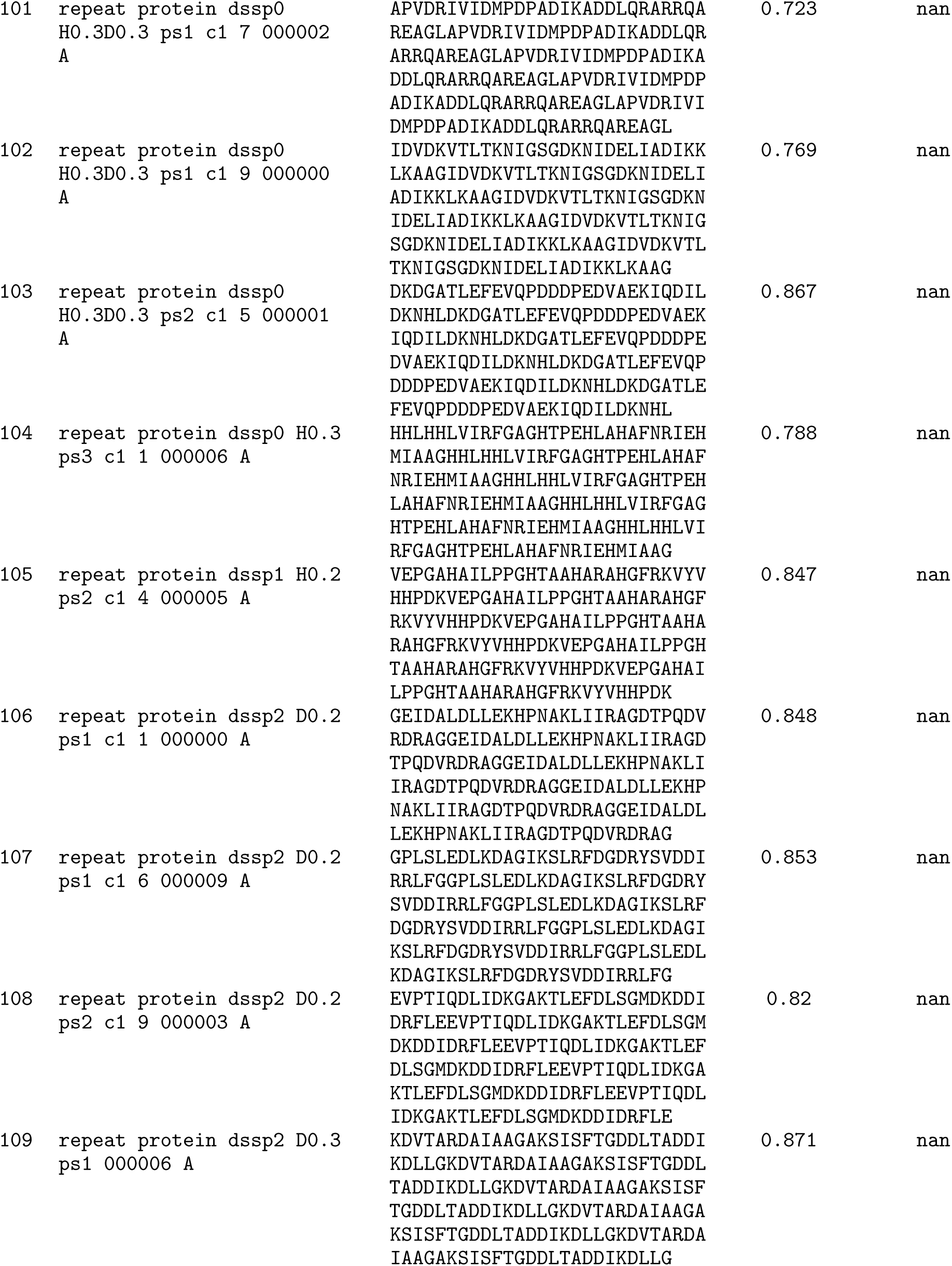
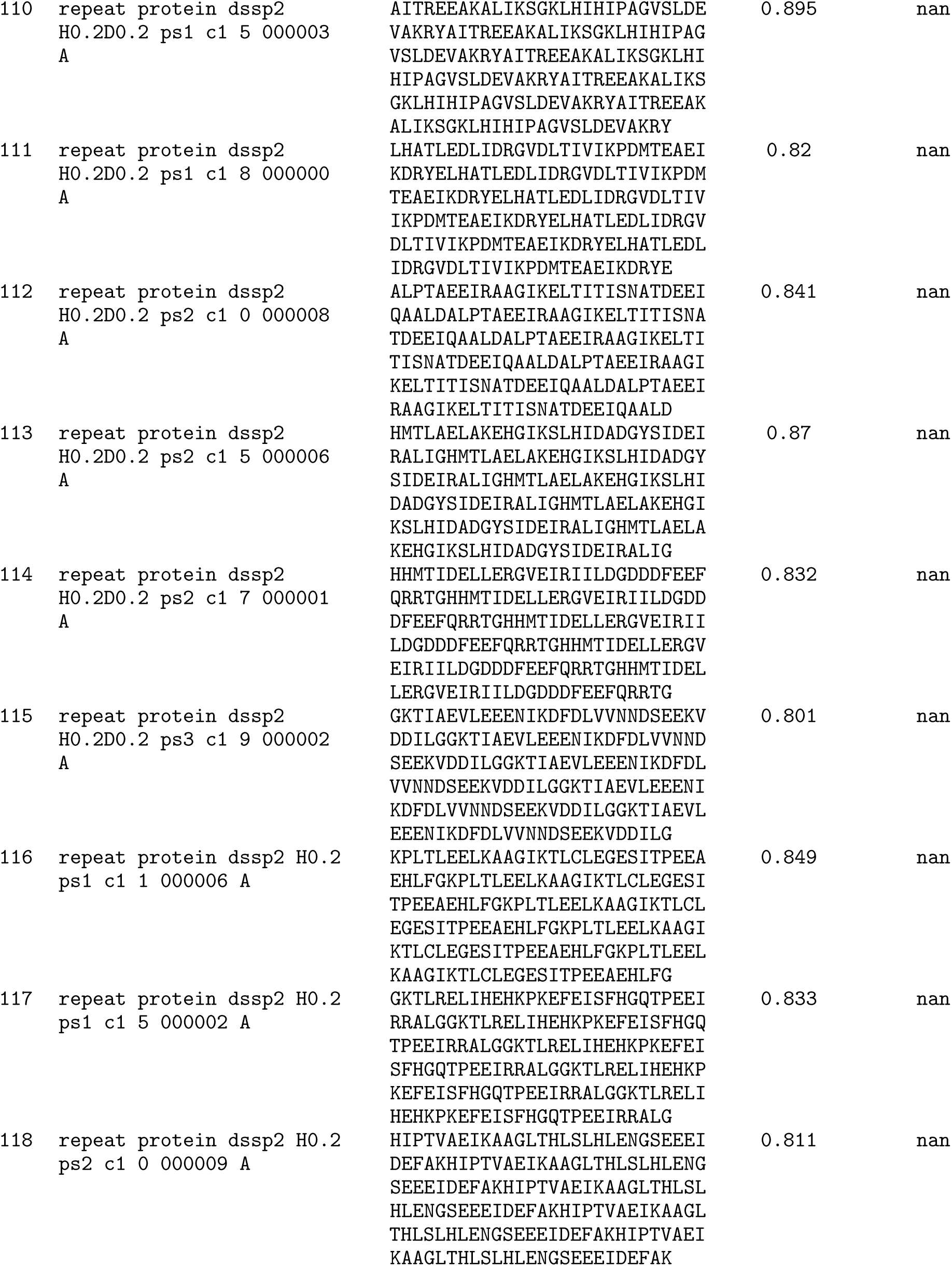
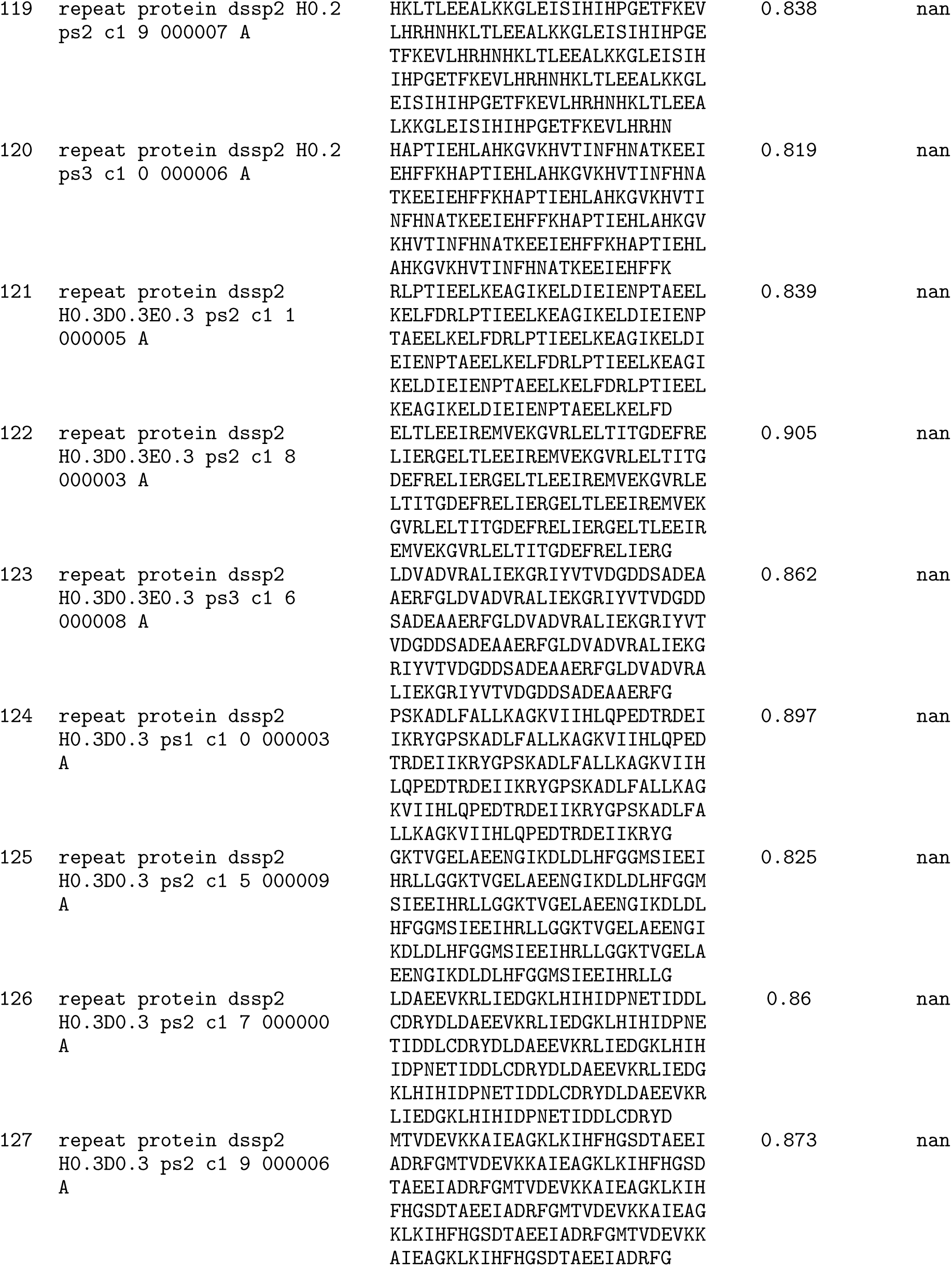
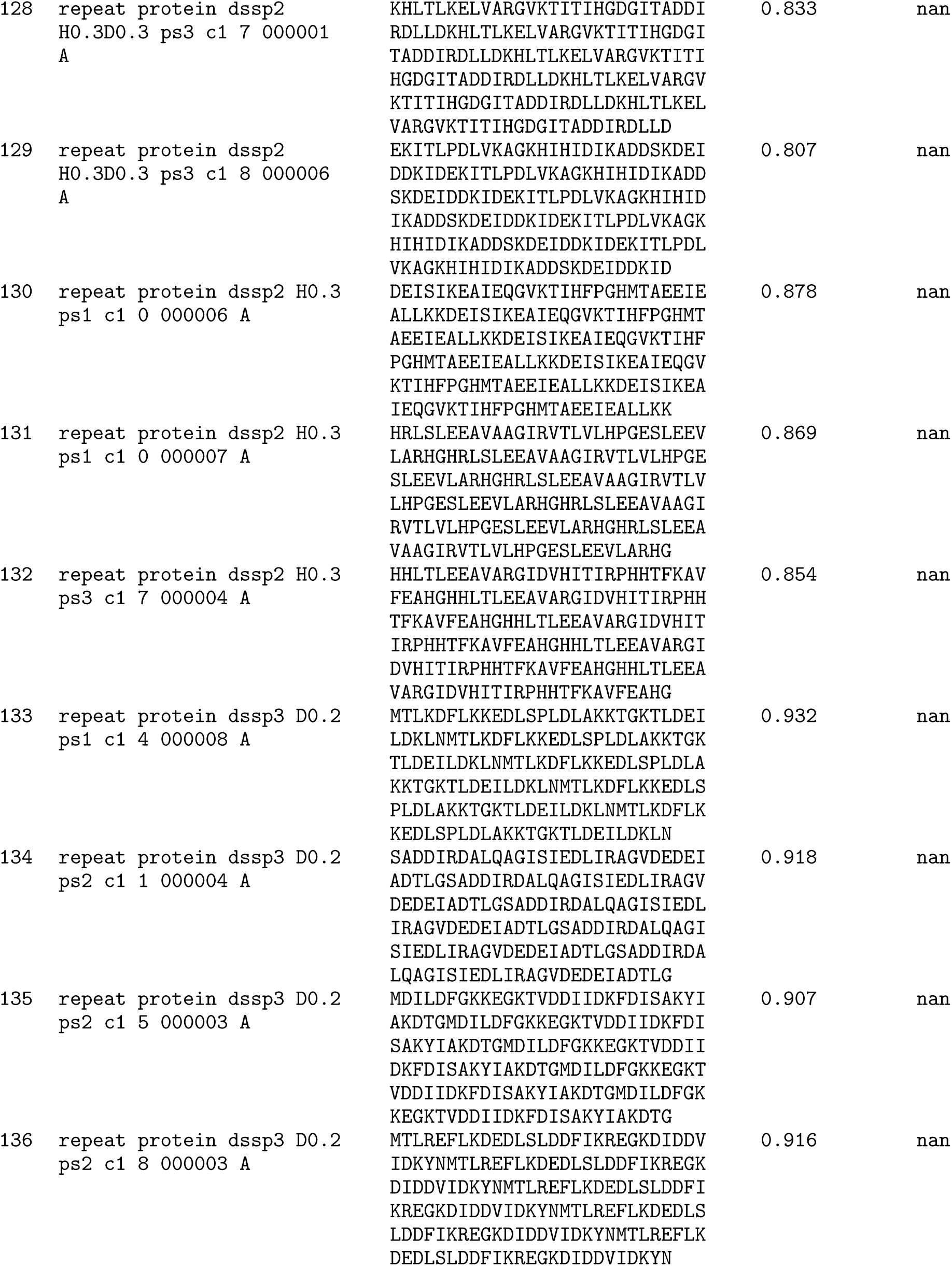
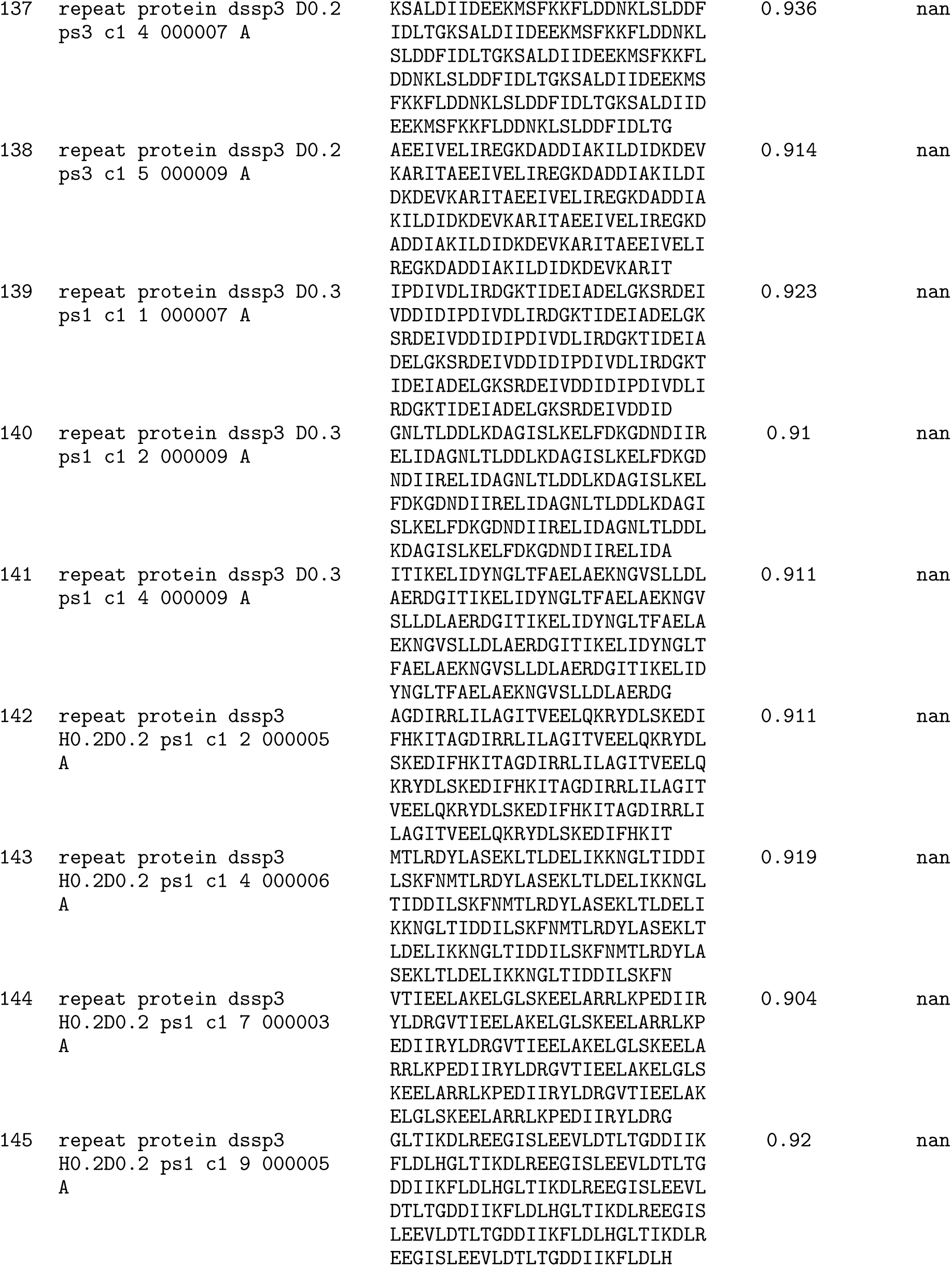
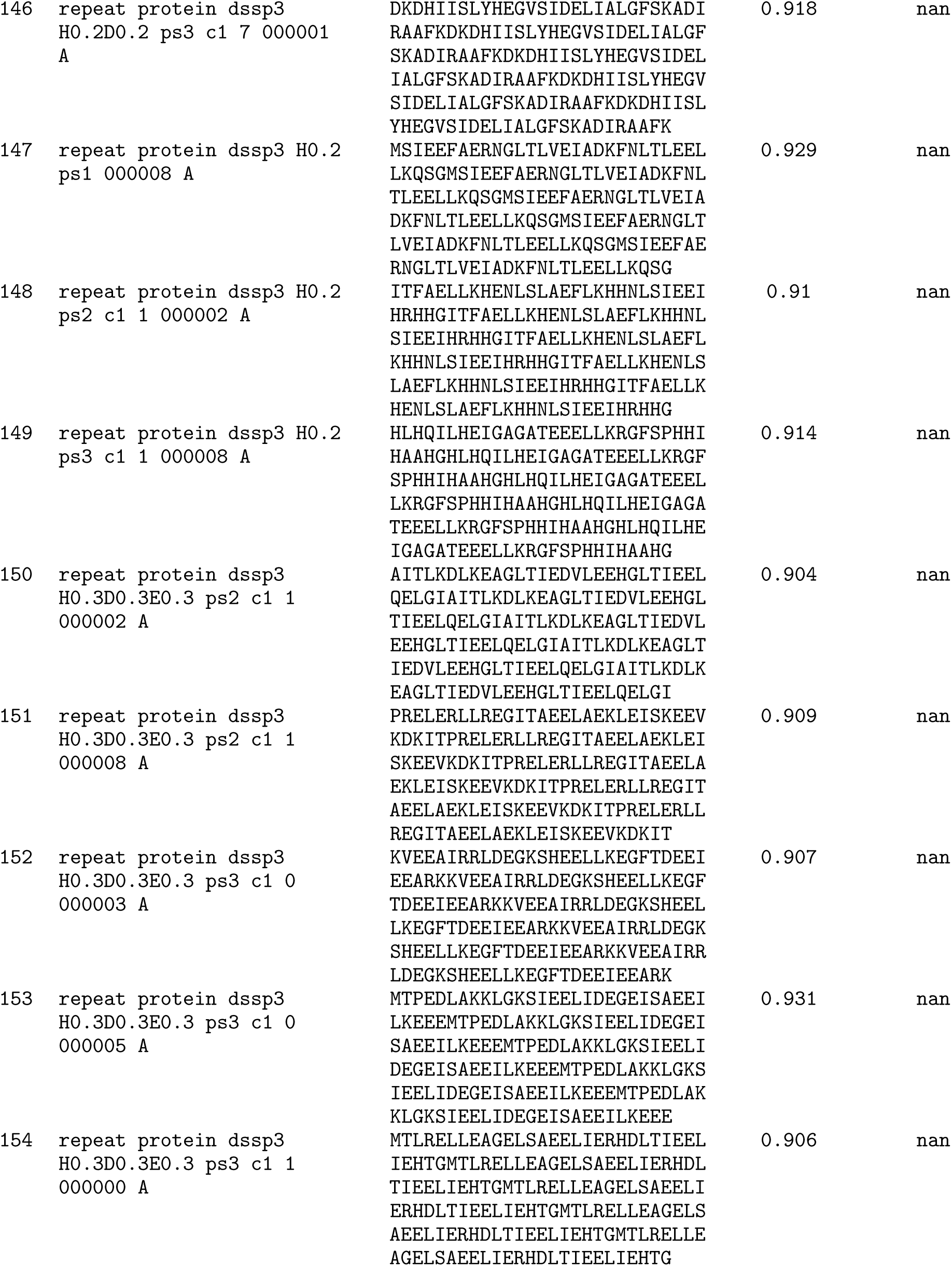
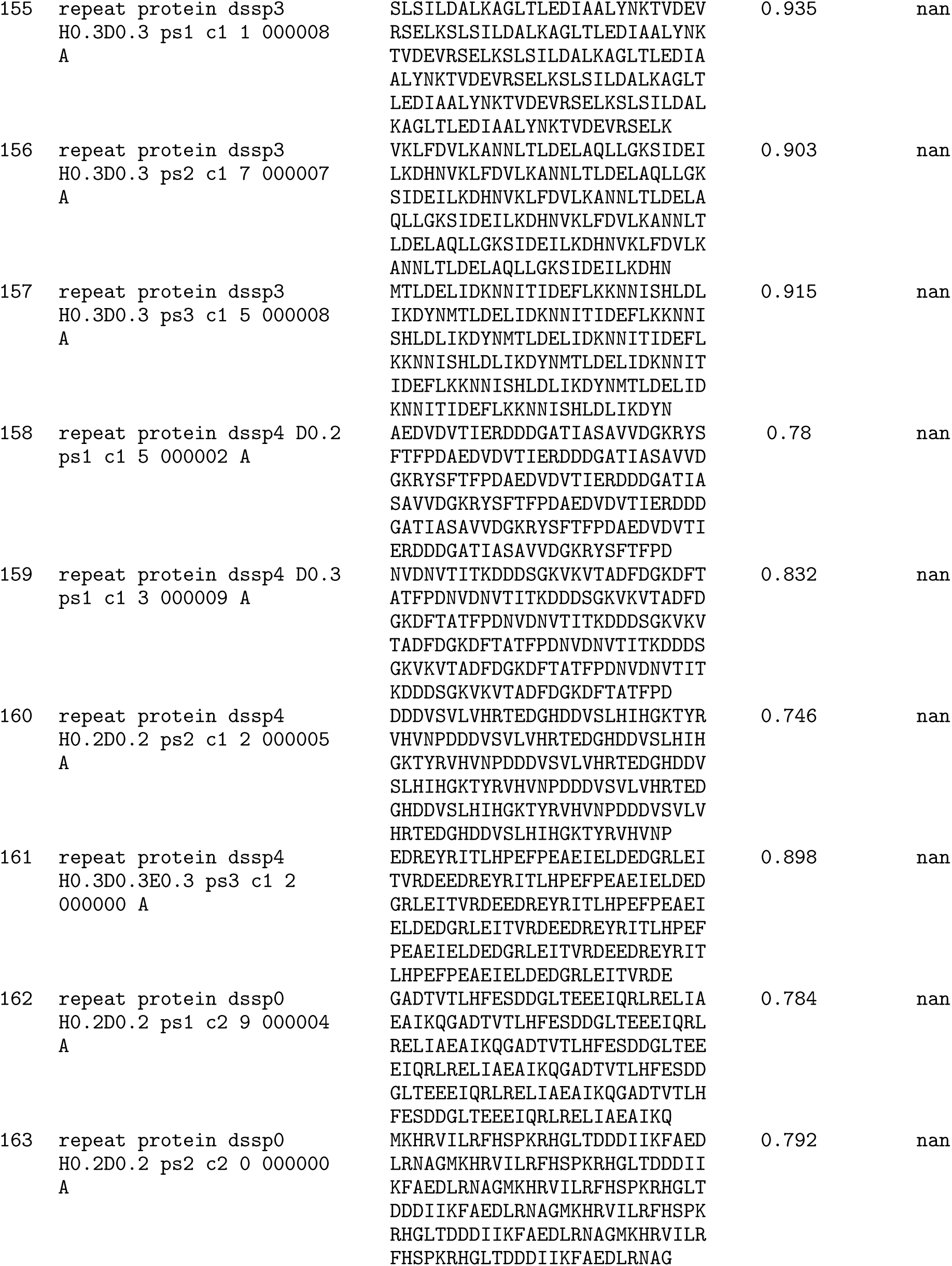
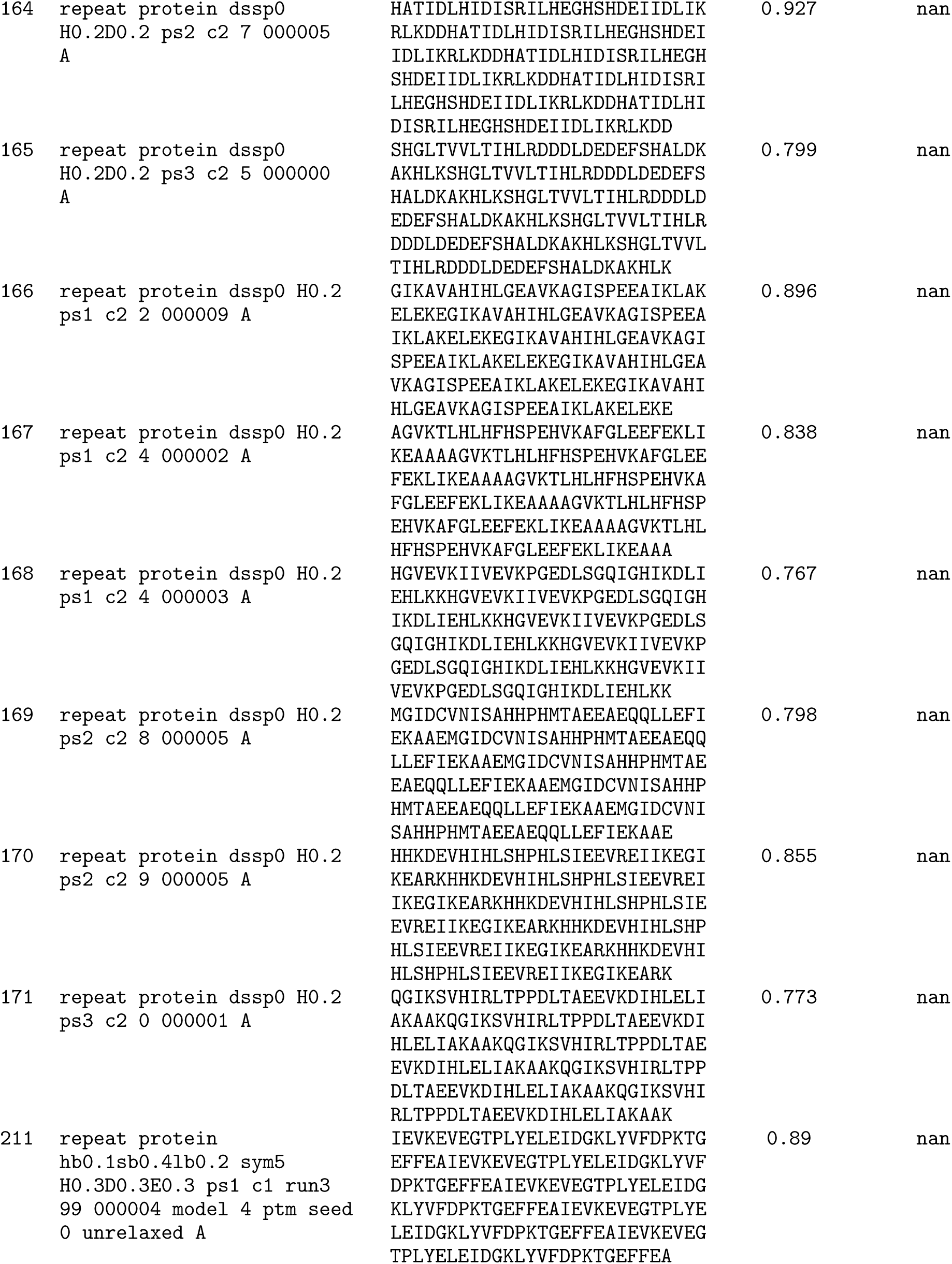
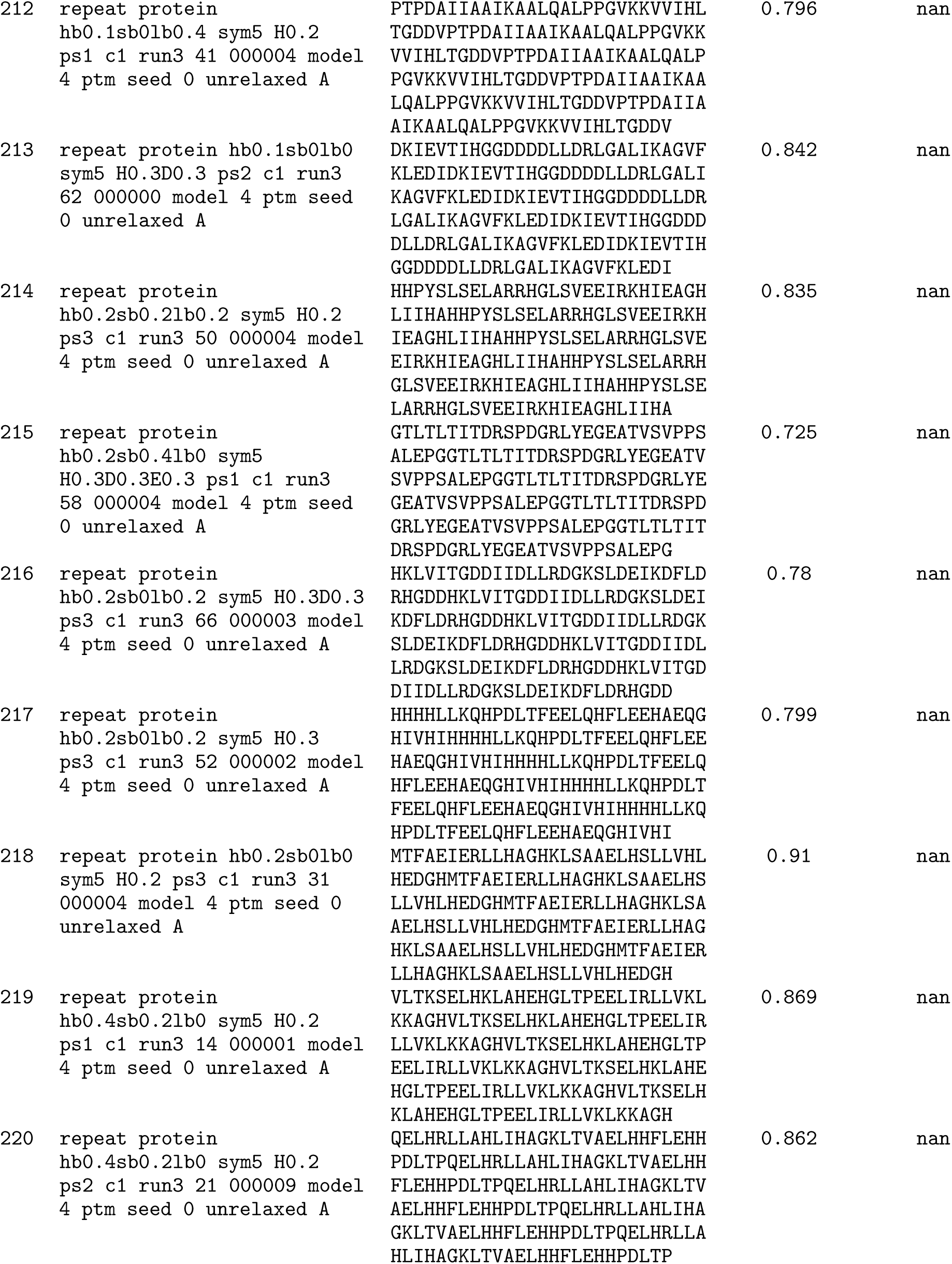
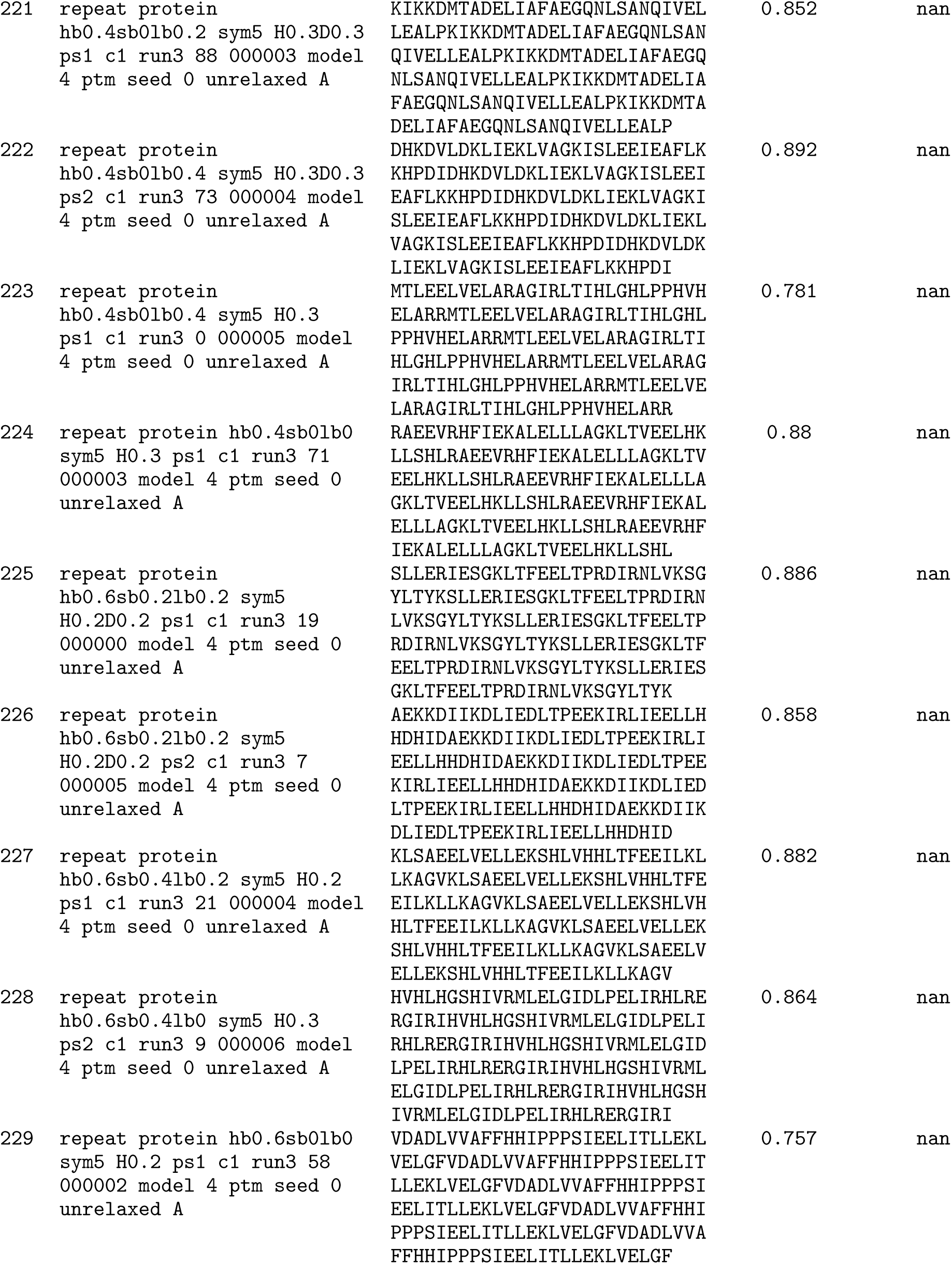
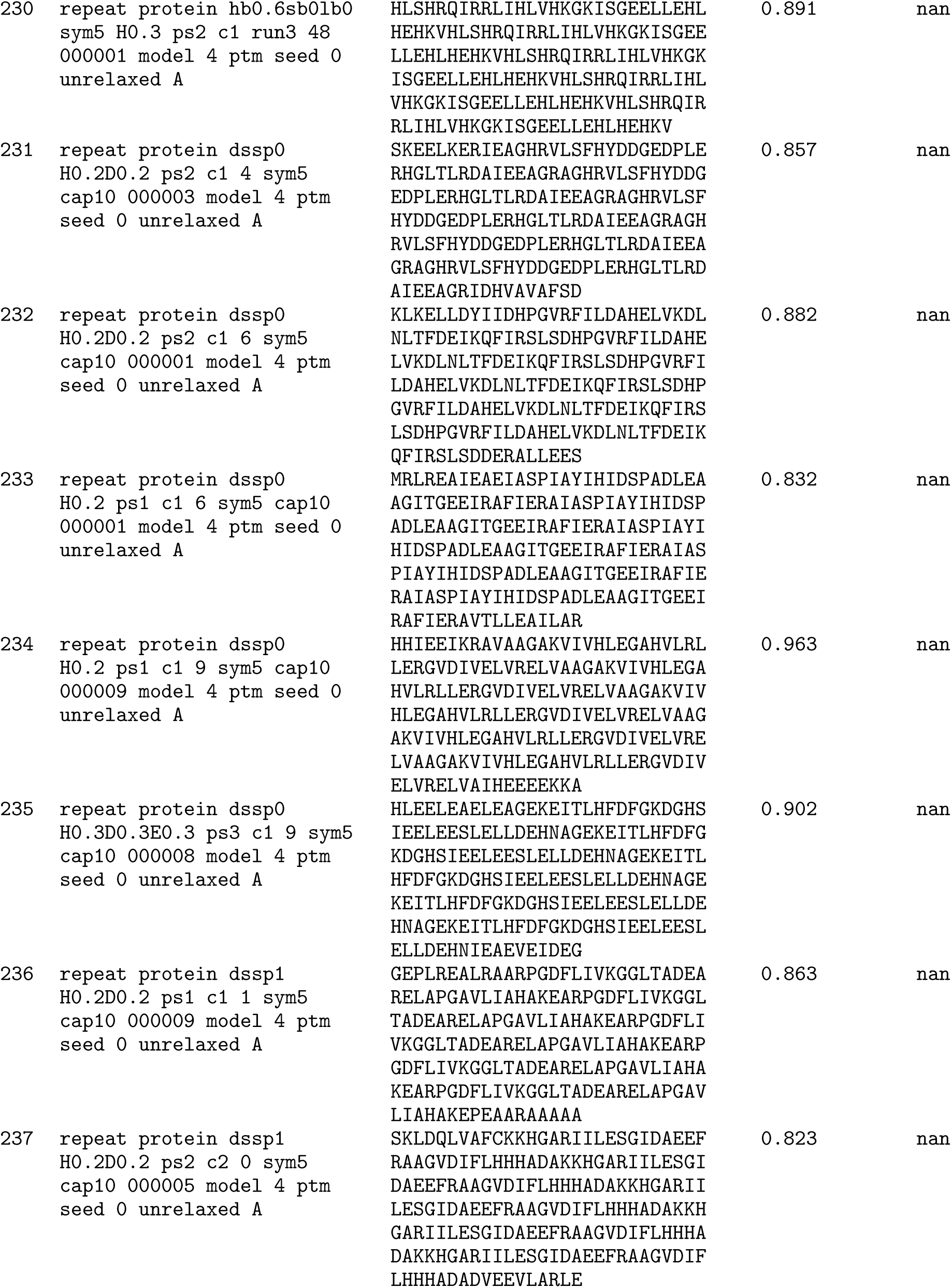
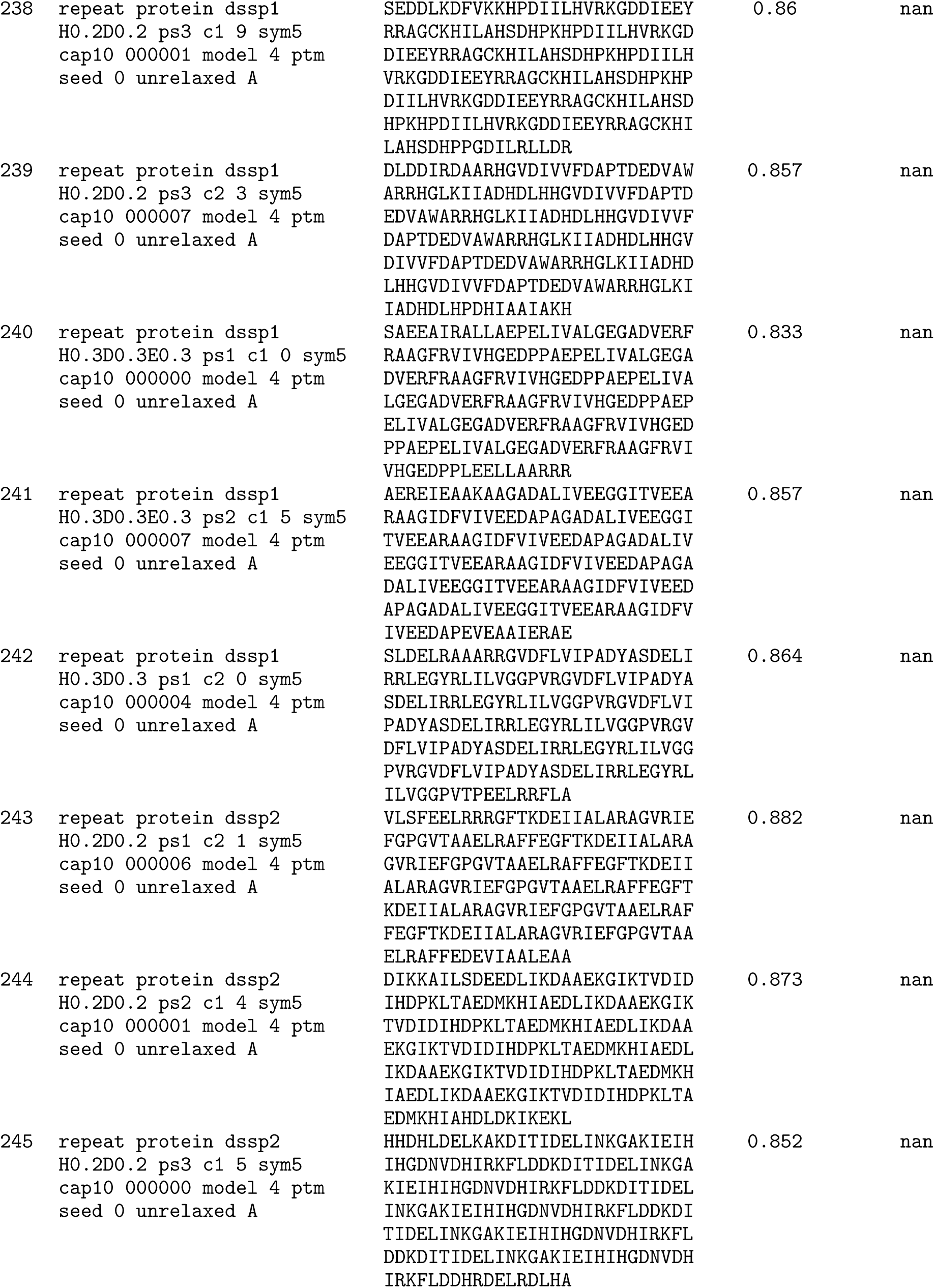
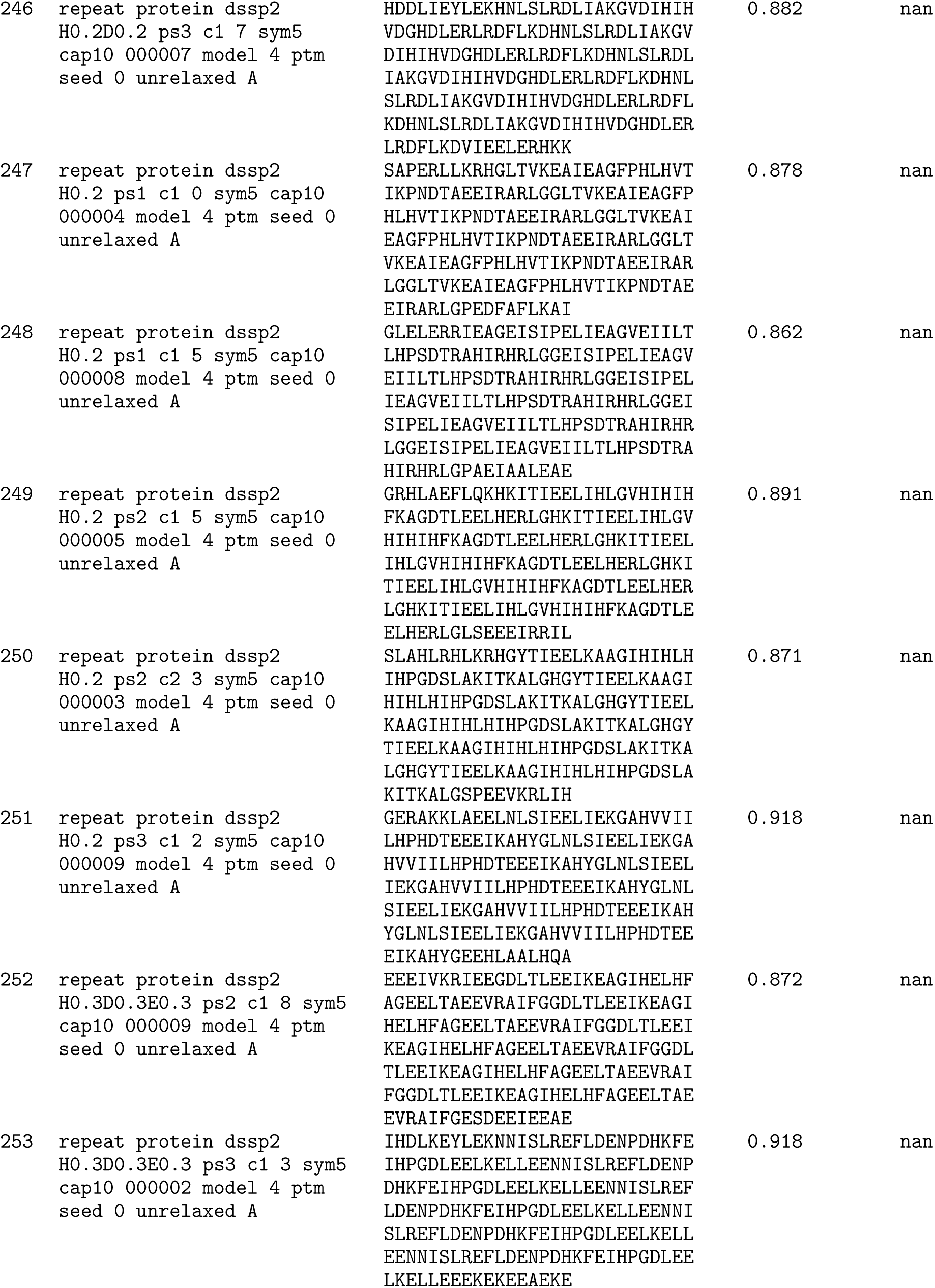
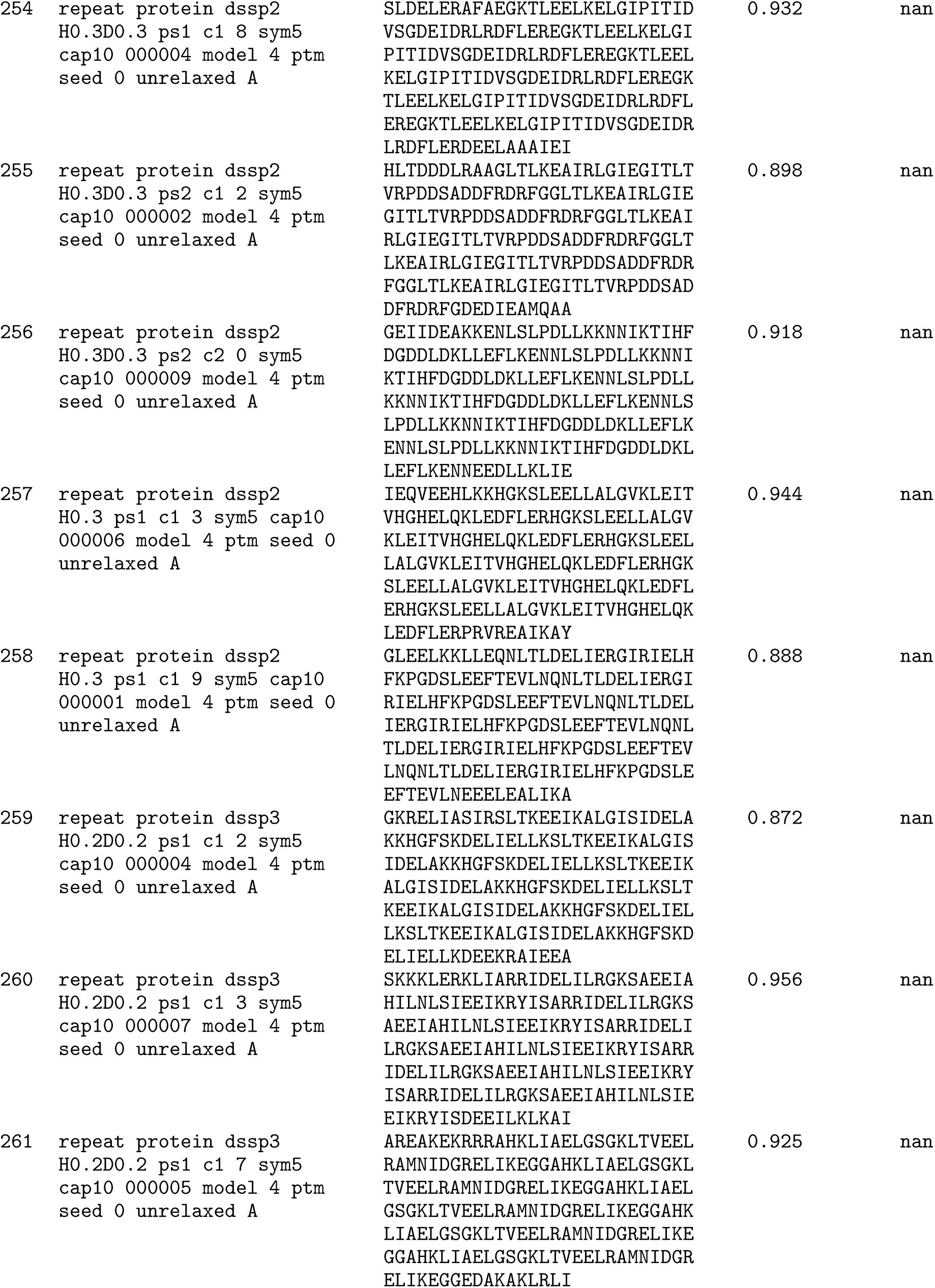
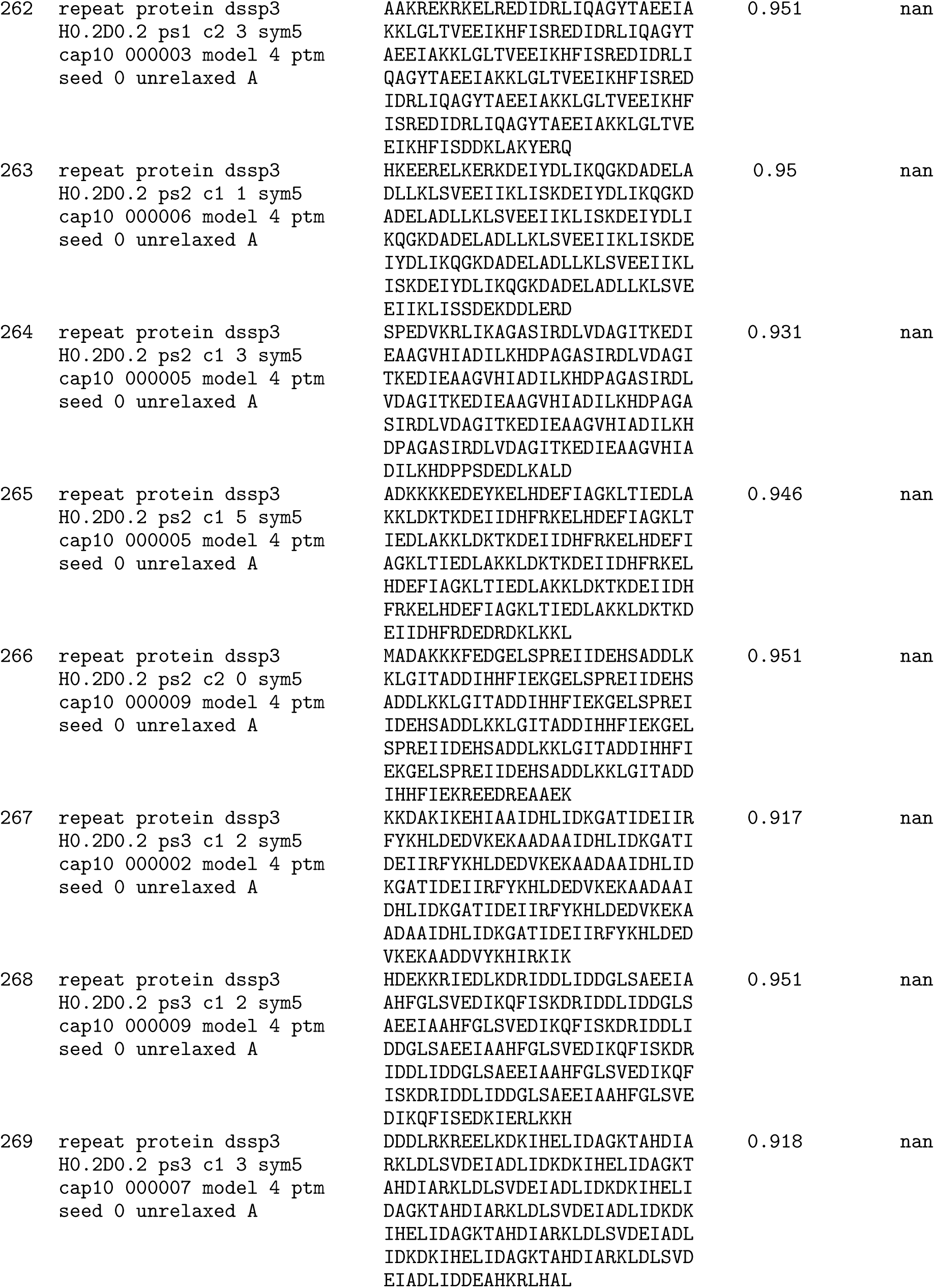
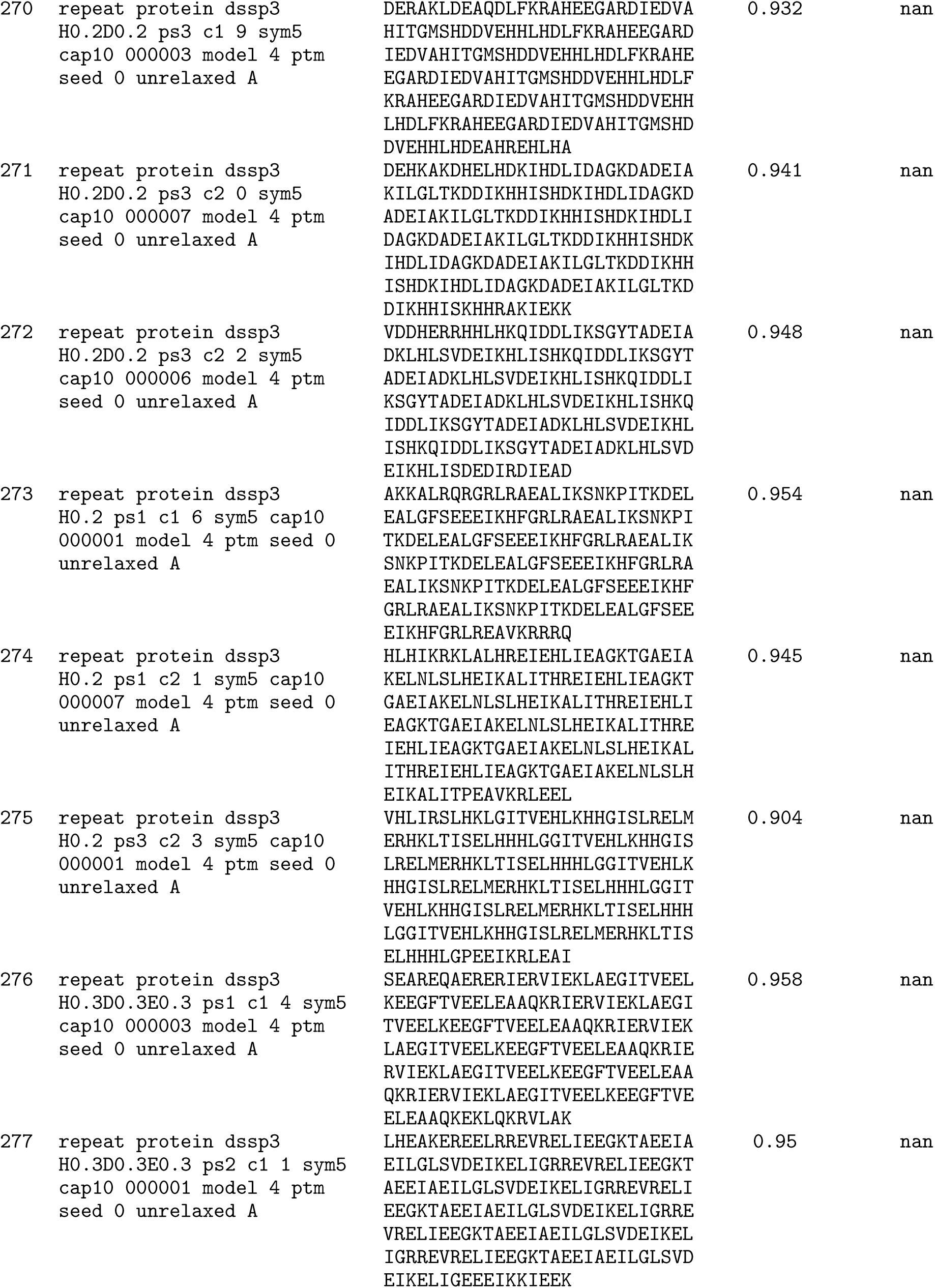
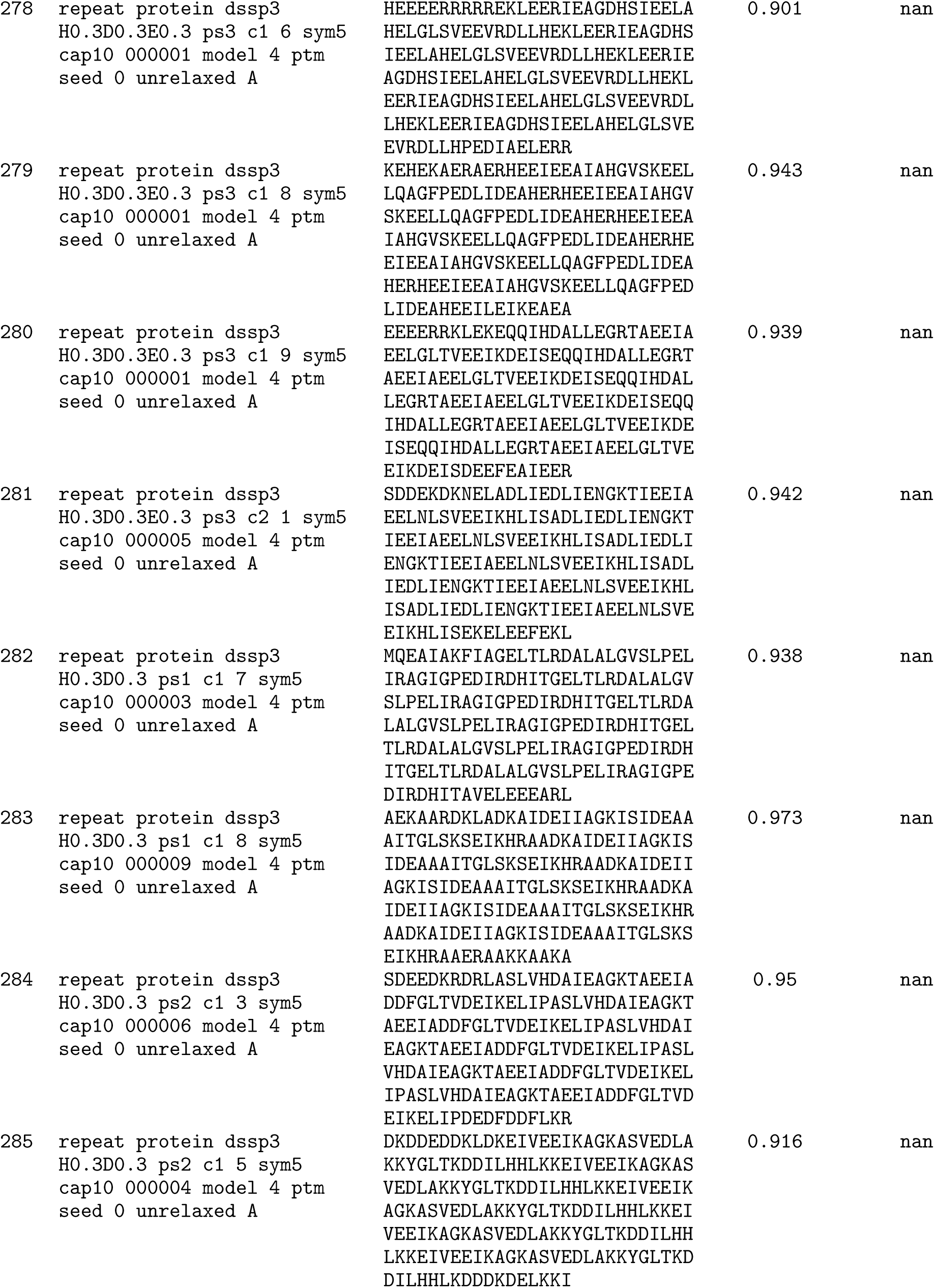
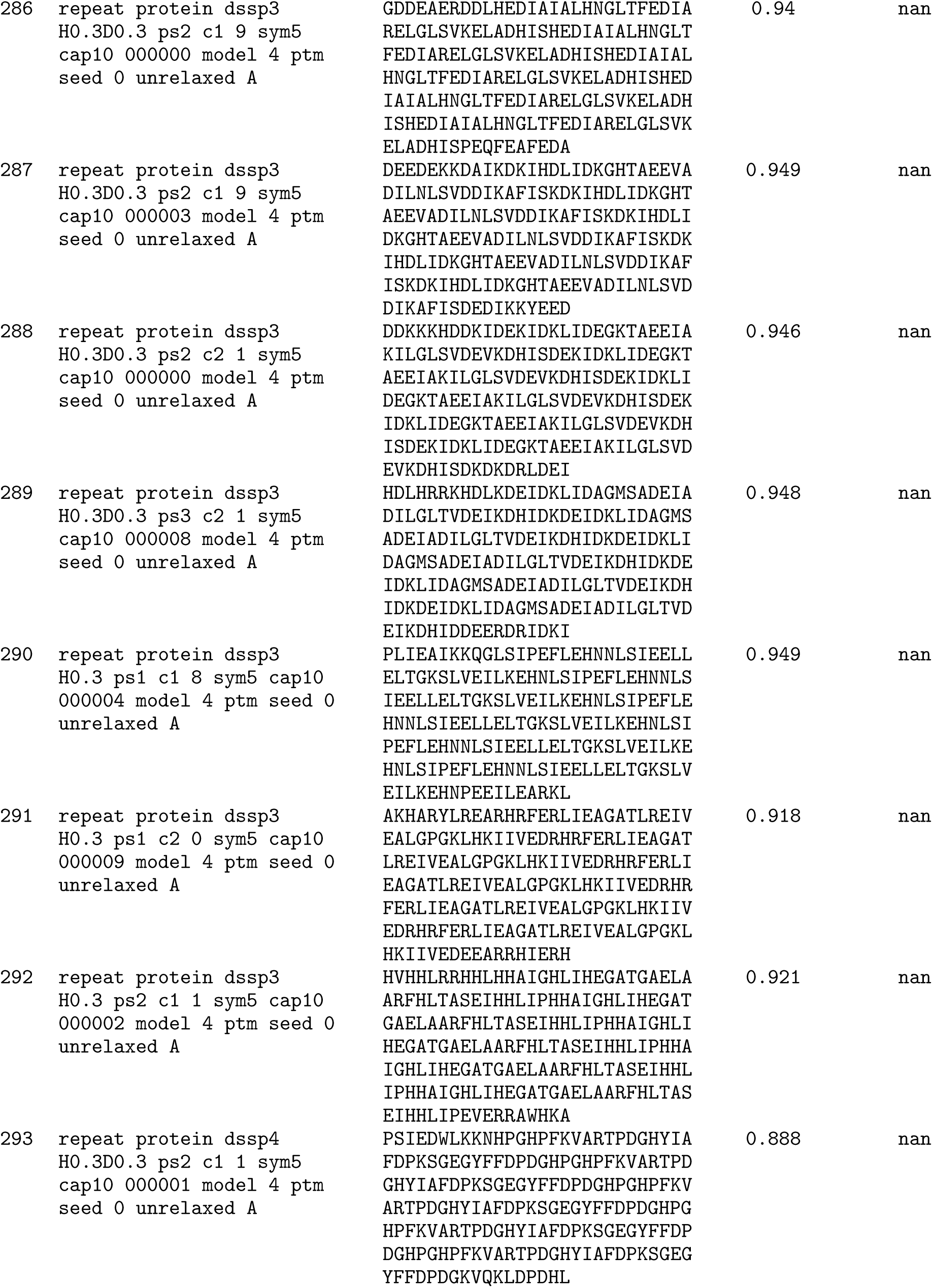
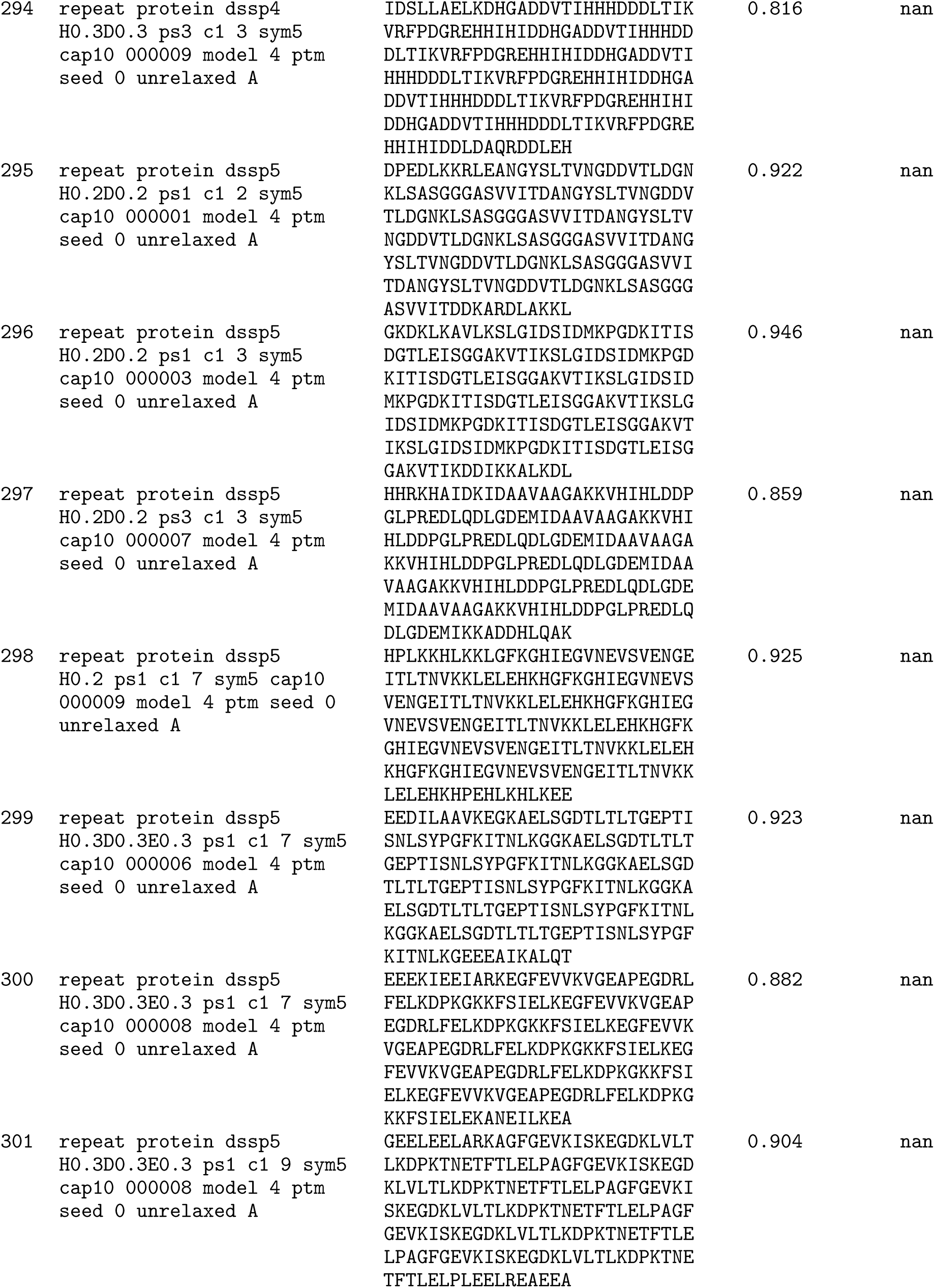
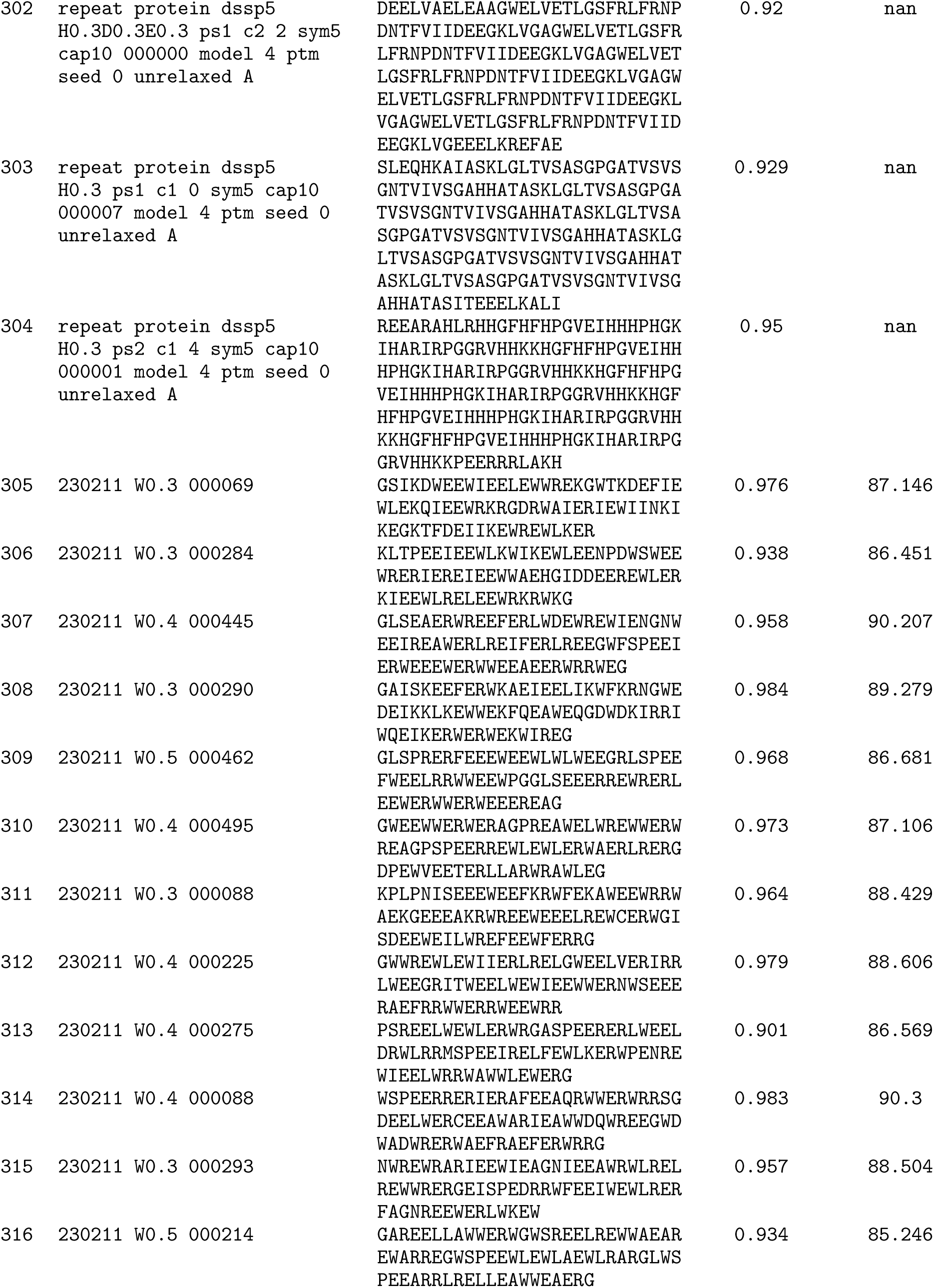
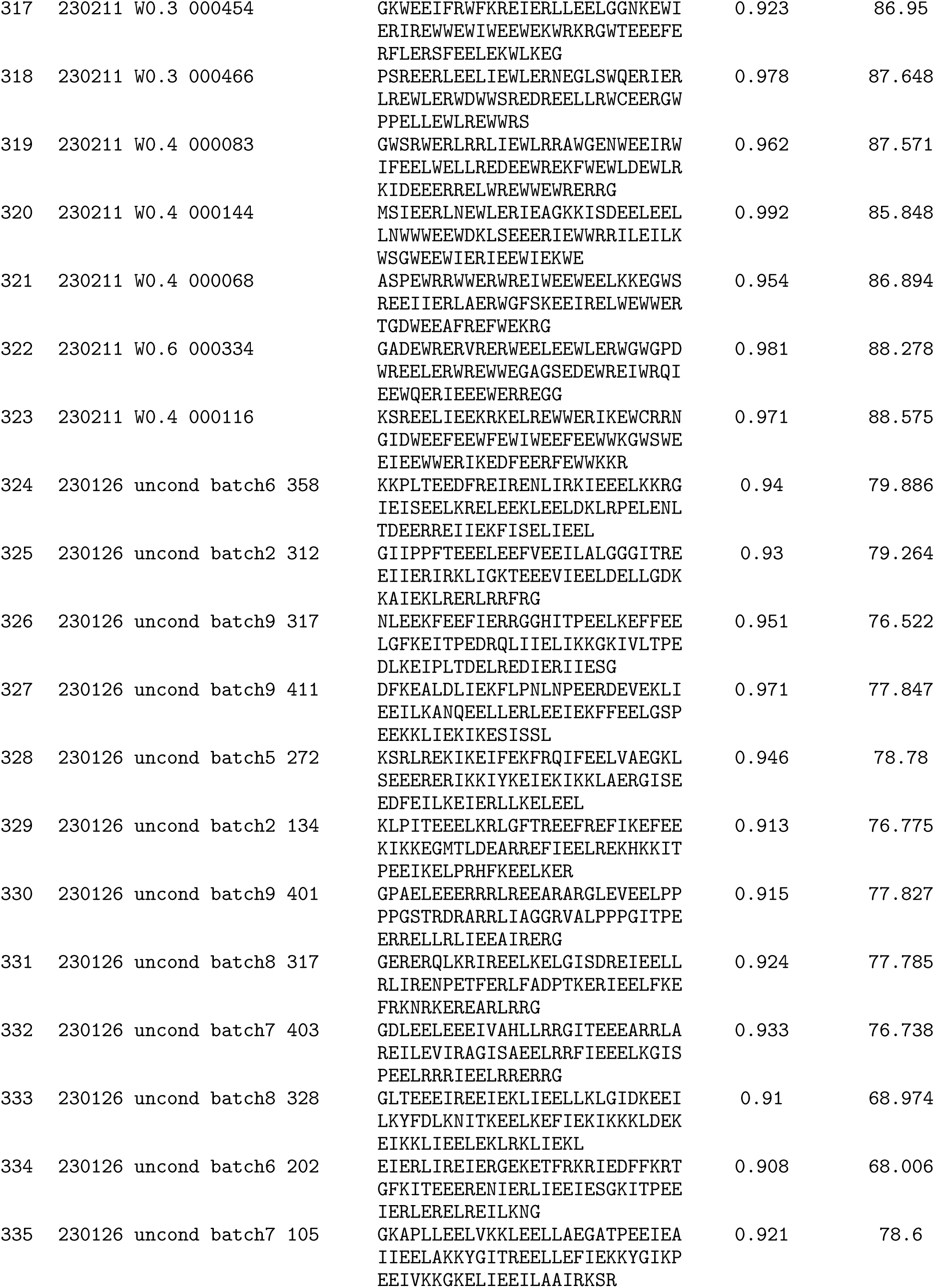
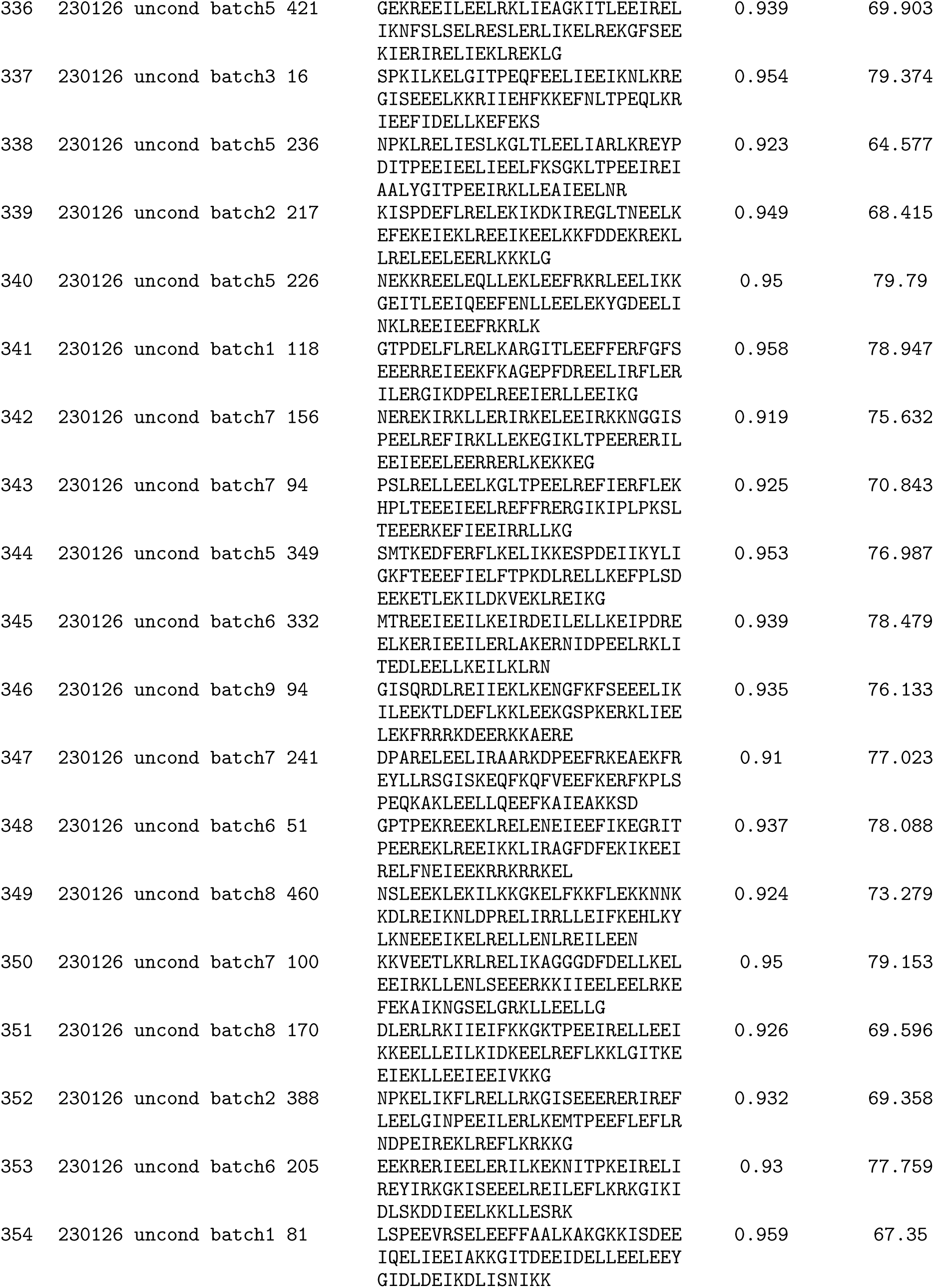
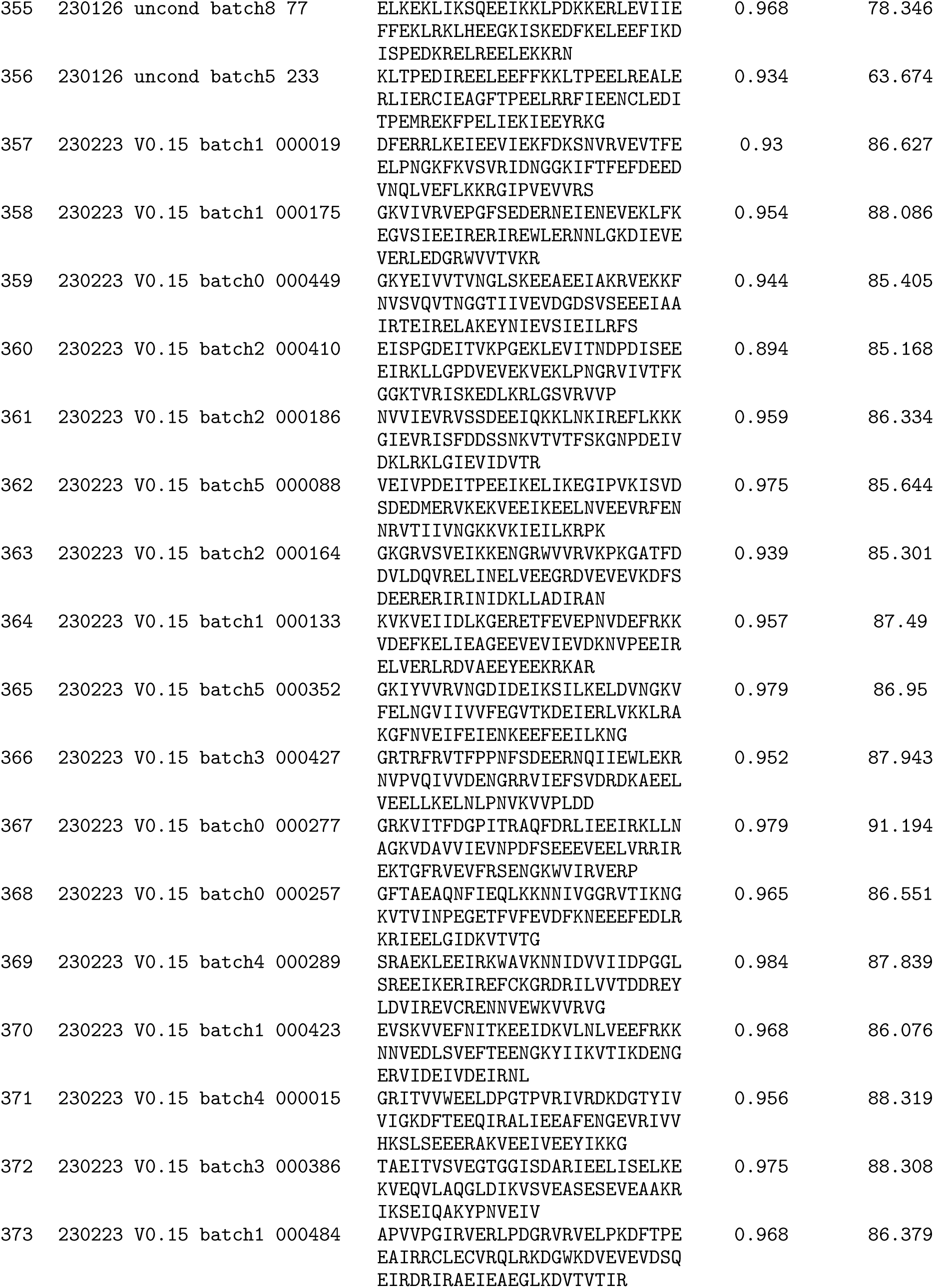
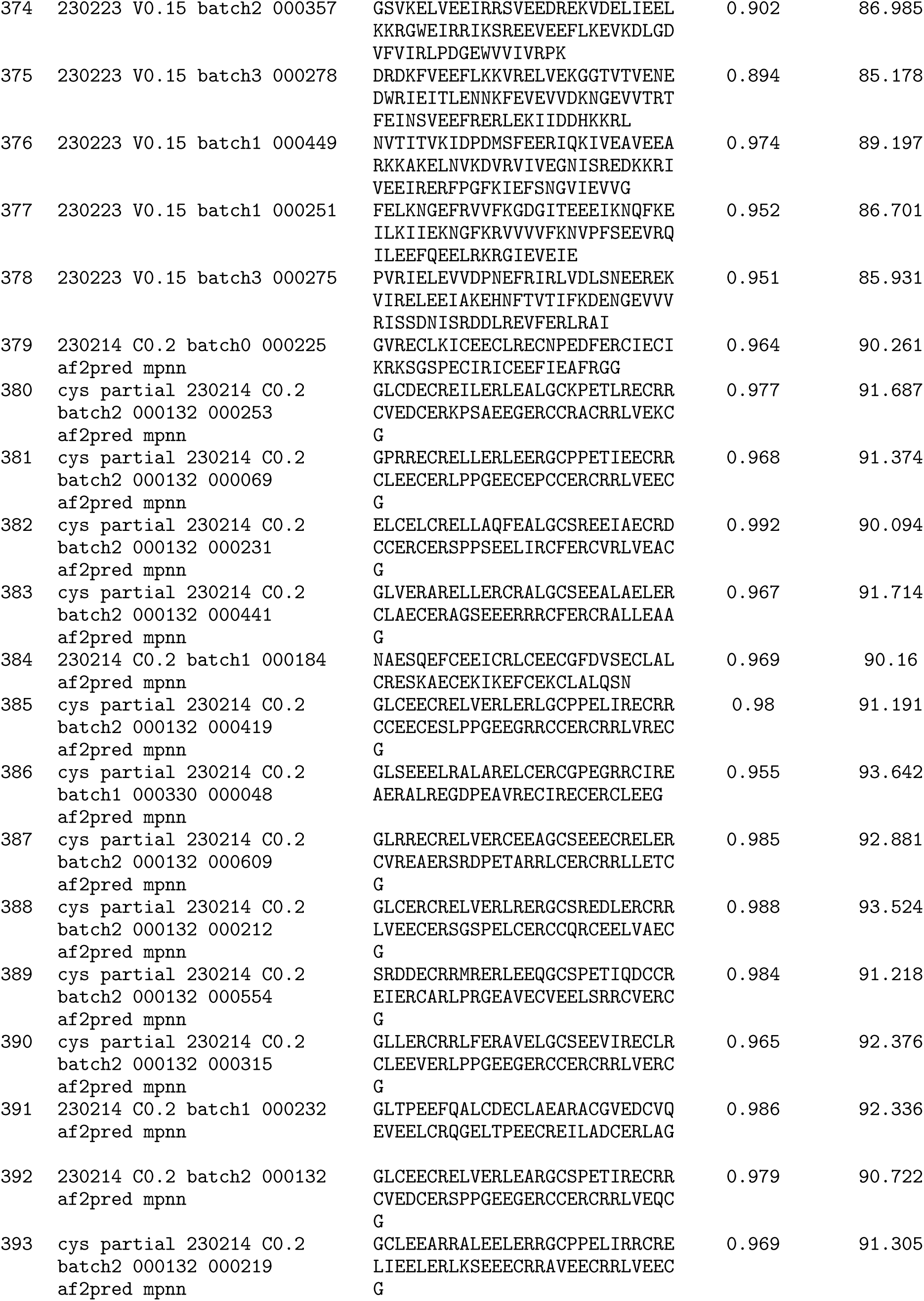
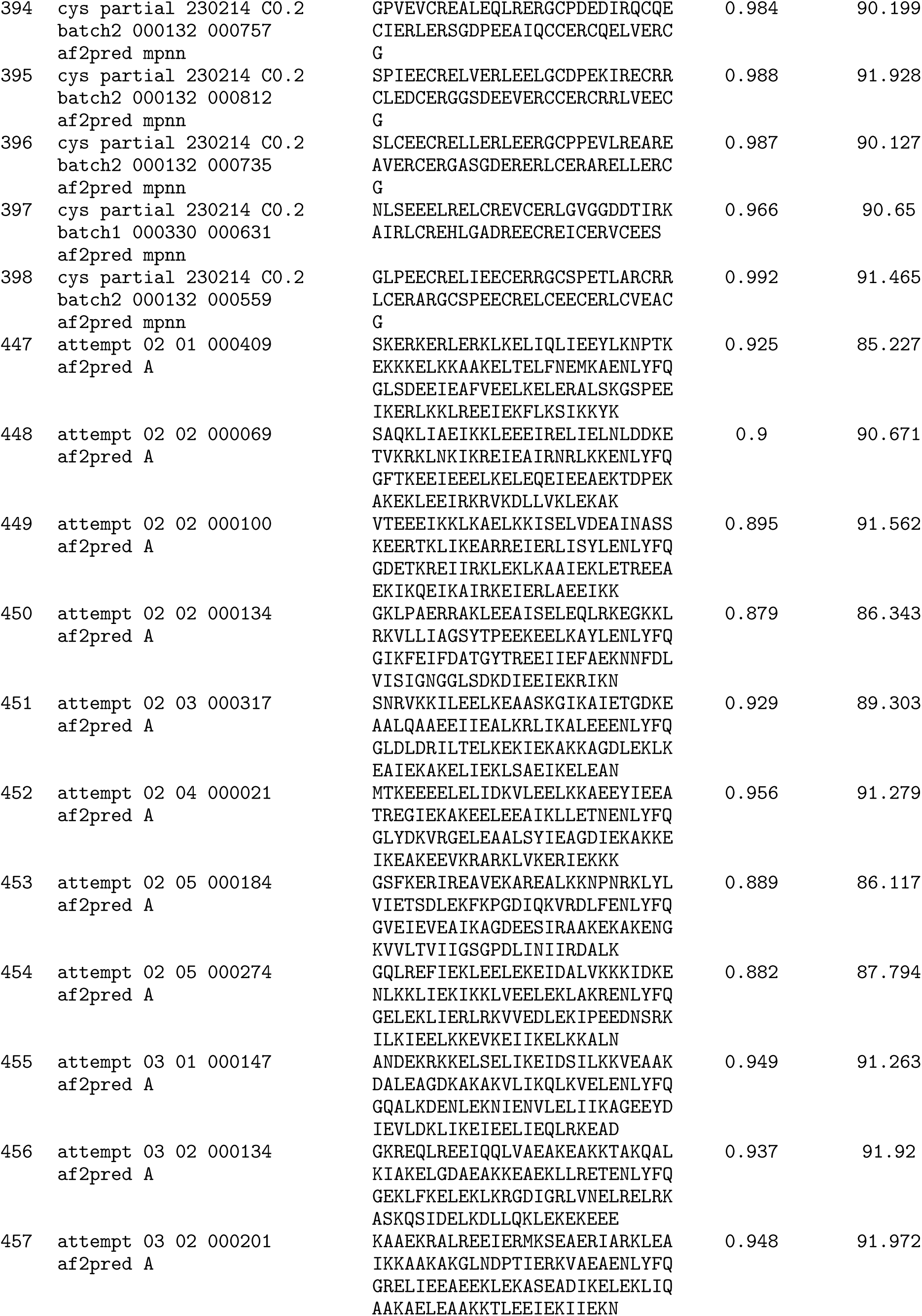
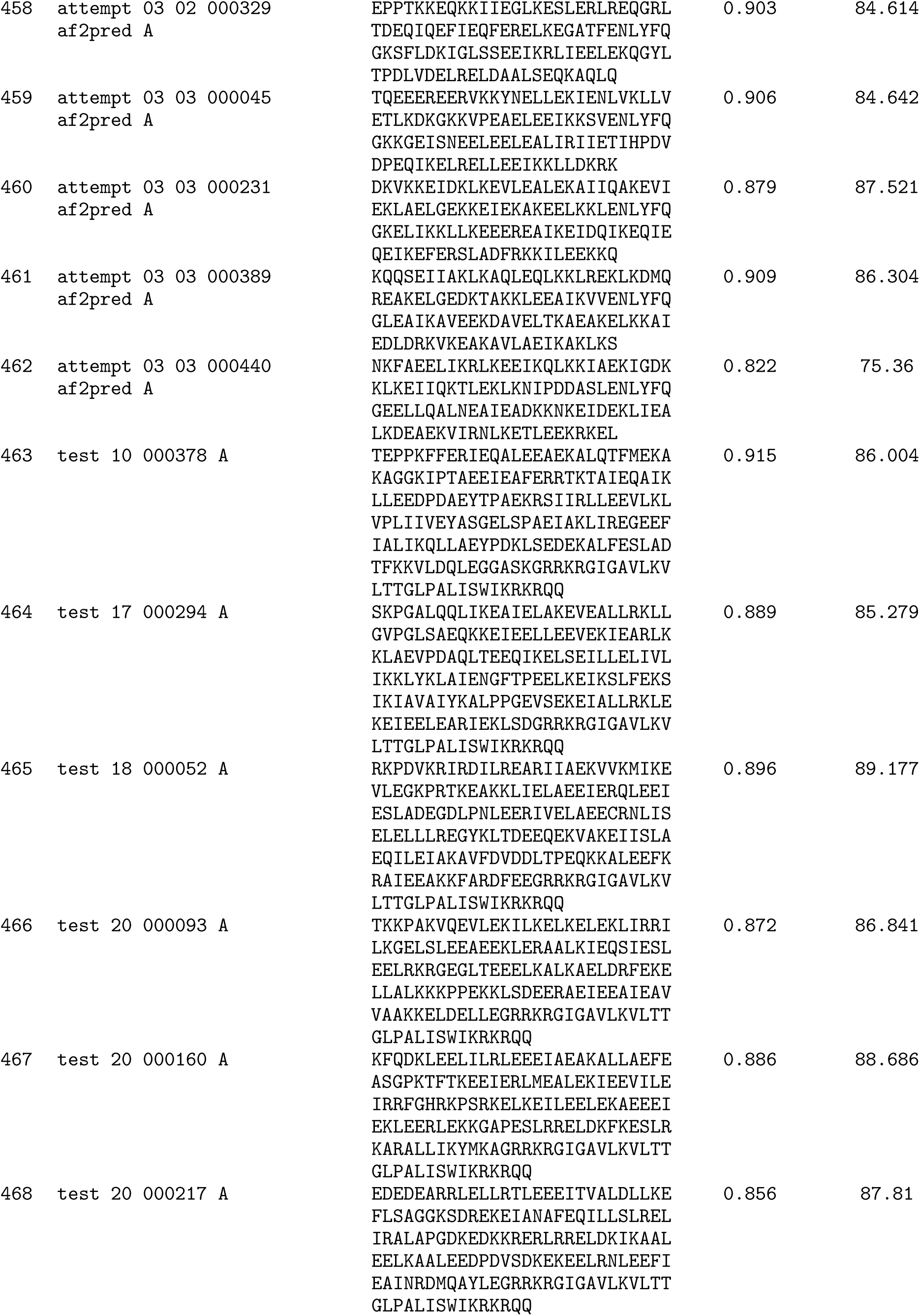
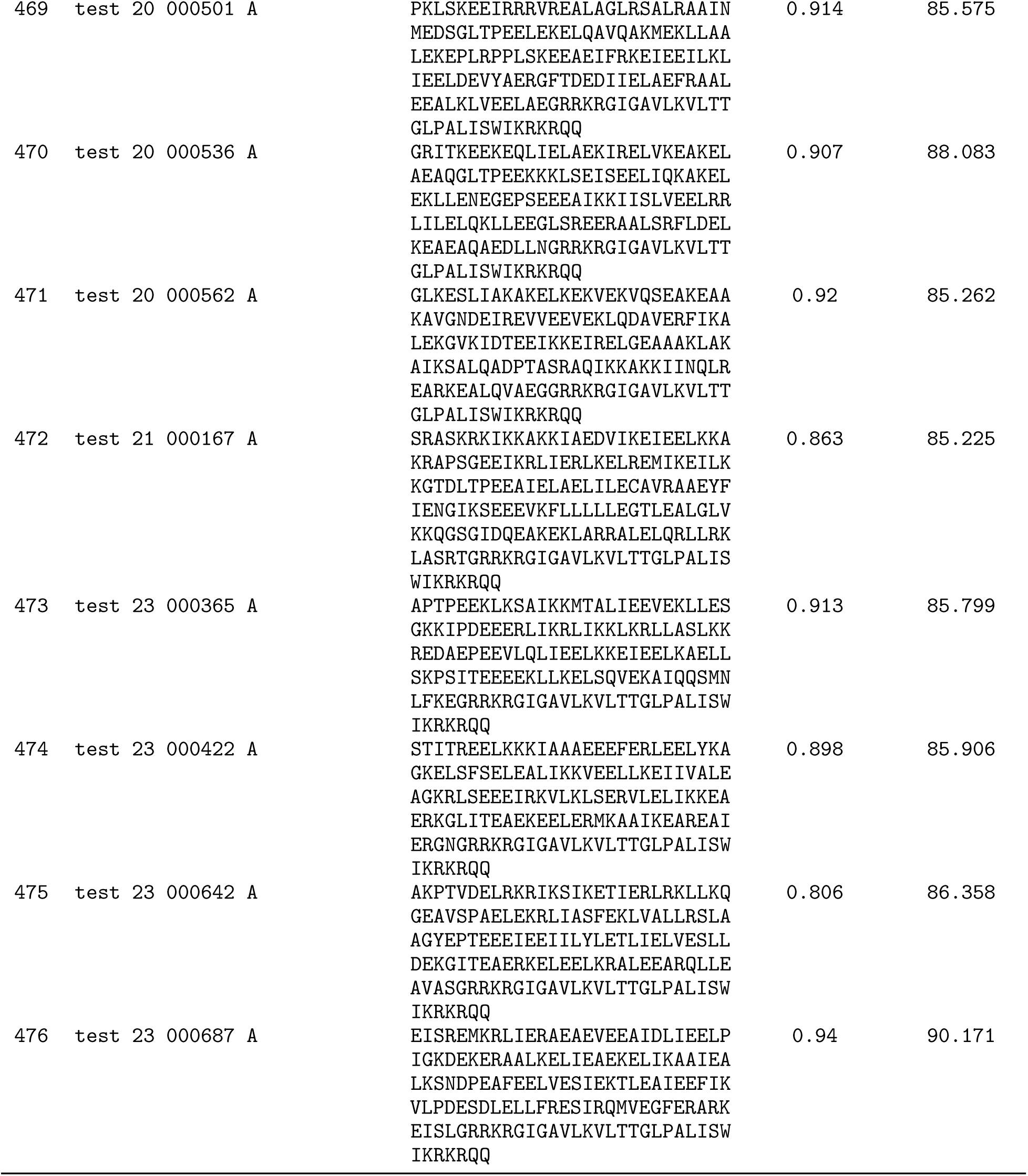
Ordered protein sequences, design confidence, and AF2 confidence metrics.

